# Modulating backbone flexibility in hydroxamate siderophores for improved iron chelation and peptide nucleic acid (PNA) delivery into bacteria

**DOI:** 10.1101/2025.09.22.677772

**Authors:** Uladzislava Tsylents, Piotr Maj, Mateusz Wdowiak, Jan Stadnicki, Adam Mieczkowski, Monika Wojciechowska, Joanna Trylska

## Abstract

Peptide nucleic acid (PNA) is a synthetic oligonucleotide analog with a peptide-based backbone that selectively binds with high affinity to natural nucleic acids. PNA is a valuable tool in antisense technology with potential antibacterial applications. However, PNA cannot penetrate bacterial cells alone. To address this, we explored iron chelators - siderophores - as PNA carriers. Bacteria acquire iron through siderophores, which are transported via specific TonB-dependent receptors in the bacterial envelope. Previously, we demonstrated that a synthetic hydroxamate-type siderophore (S_L_) exploited this transport system to deliver PNA into bacterial cells, achieving a gene-silencing effect. However, this transport was limited to an *Escherichia coli* mutant with continuous iron uptake, and was not observed in wild-type *E. coli*. In this study, we developed a new synthetic siderophore (S_GLY_) with glycine spacers between modified ornithine residues for enhanced flexibility and iron-binding. We also synthesized marine siderophore analogs (M_GLY_ and M_ALA_) inspired by natural moanachelins. Using circular dichroism spectroscopy, spectrophotometric assays, and molecular dynamics simulations, we confirmed iron binding. Growth recovery experiments showed S_GLY_ recognition and internalization via the TonB-dependent transport system, likely using hydroxamate siderophore pathways. The M_GLY_ and M_ALA_ siderophores showed lower growth promotion than S_GLY_, indicating less efficient internalization. Molecular docking revealed high affinity of S_GLY_ for *E. coli* receptors involved in the uptake of hydroxamate siderophores. However, upon conjugation to PNA, all three siderophores effectively delivered PNA into *E. coli* cells. We confirmed PNA-mediated gene silencing using fluorescence measurements and confocal microscopy.

## 1. INTRODUCTION

Antibiotic resistance is an increasing global health threat as conventional antibiotics lose effectiveness, while the development of new antimicrobials remains slow. If this trend continues, a return to the pre-antibiotic era - where even minor injuries could result in fatal outcomes - becomes a realistic concern. Therefore, development of alternative antimicrobial strategies is critical (Abbas et al., 2024). Antisense therapy is a promising alternative in the fight against infectious diseases (Collotta et al., 2023). It uses chemically modified antisense oligonucleotides to modulate gene expression by targeting RNA encoding proteins essential for pathogen survival. Peptide nucleic acid (PNA) is a synthetic oligonucleotide analog often used as an antisense oligomer. In PNA, standard nucleobases are linked to an *N*-(2-aminoethyl)-glycine backbone instead of the sugar-phosphate backbone found in DNA or RNA (Nielsen et al., 1991). This neutrally charged, peptide-based backbone, designed by Peter Nielsen and colleagues, enhances the affinity and specificity of PNA for complementary nucleic acids (Egholm et al., 1993). As a peptide-nucleic acid hybrid, PNA is resistant to both proteolytic and nucleolytic enzymes. When binding to DNA or RNA, PNA forms stable duplexes or triplexes, effectively blocking expression of the target gene (Nielsen, 1999). PNA sequences have been established for bacterial mRNA targets (recently reviewed in (El-Fateh et al., 2024; Moreira et al., 2024; Pals et al., 2024; Tsylents et al., 2023)). New sequences for selected targets can be designed using different software tools (Eller et al., 2021; Górska et al., 2016; Jung et al., 2023).

Various PNA modifications have been introduced and tested over the past decades, as reviewed in (Brodyagin et al., 2021; Wojciechowska et al., 2020). Despite significant efforts, PNA cannot penetrate bacteria unaided due to the complex and charged bacterial envelope. A common solution is conjugating PNA to carriers to facilitate cellular uptake. Initially, Nielsen et al. attached PNA to a cell-penetrating peptide, (KFF)_3_K, which showed antibacterial activity in *E. coli* at 1 µM (Good et al., 2001). In that study, they introduced a PNA sequence targeting the *acpP* transcript encoding the essential acyl carrier protein involved in fatty acid biosynthesis. This PNA sequence, complementary to *acpP* mRNA, was later optimized (Goltermann et al., 2019). Unfortunately, poor peptide stability and hemolytic activity preclude (KFF)_3_K from medical applications (Vaara & Porro, 1996; Yavari et al., 2021). Nevertheless, PNA conjugates with (KFF)_3_K are still extensively used for antibacterial research (Campion et al., 2024; C. Ghosh et al., 2024; Goltermann et al., 2022; Hizume et al., 2023; Patenge et al., 2013; Sarkar et al., 2025; Siekierska et al., 2024). Conjugation site, linker length, and type were shown to be critical in PNA conjugates (Klabenkova et al., 2021; Siekierska et al., 2024). Recently, we conjugated hydrocarbon-stapled (KFF)_3_K and an antimicrobial peptide, anoplin, to PNA using either cleavable disulfide bridges or non-cleavable ethylene glycol linkers (Siekierska et al., 2024). PNA conjugates with stapled peptides did not improve antibacterial activity compared to stapled peptides alone, suggesting that structurally stabilized peptides exhibit lower carrier potential against gram-negative bacteria than unstapled analogs. These findings suggest that peptide internal flexibility may be a necessary feature to promote PNA delivery into bacterial cells.

Finding efficient delivery compounds is crucial for antibacterial PNA applications. For PNA delivery into bacteria, we turned to the Trojan horse strategy and used nutrients recognized by bacterial transport systems. Our group has employed the bacterial TonB-dependent transport (TBDT) system as an entry route for PNA. This specialized uptake system actively transports nutrients required by bacteria, such as vitamin B_12_, maltose, heme, and siderophores, via selective receptors (Silale & Van Den Berg, 2023). Indeed, we showed that vitamin B_12_ covalently conjugated with PNA delivers it to *E. coli* and *Salmonella enterica* serovar Typhimurium cells using TBDT (Równicki et al., 2017). However, although vitamin B_12_-PNA conjugates targeting the essential *acpP* gene inhibited *E. coli* growth, they were less efficient than the (KFF)_3_K peptide used as a PNA carrier (Równicki et al., 2019). Recently, a glucose polymer (GP) was used to exploit bacterial sugar transporters to deliver PNA conjugated to silicon nanoparticles (SiNPs) (Liu et al., 2023). The GP-SiNPs-PNA conjugate showed a minimum inhibitory concentration (MIC) of 0.8 µM against multidrug-resistant *E. coli* and methicillin-resistant *Staphylococcus aureus* strains and demonstrated effective *in vivo* infection clearance in mouse models.

Therefore, to further investigate the potential of TBDT-dependent PNA uptake, we focused on siderophore receptor pathways (Figure 1). Bacteria produce siderophores to retrieve iron(III) from the environment. Siderophores are crucial for bacterial growth by overcoming limited iron solubility and bioavailability. They typically employ oxygen atoms in octahedral geometry to coordinate iron(III). Depending on the groups that bind iron(III), siderophores are classified as catecholate, hydroxamate, carboxylate, or mixed type. The TonB-dependent *E. coli* uptake system for natural catecholate (enterobactin and Fe^3+^-2,3-dihydroxybenzoylserine (Fe^3+^-DHBS)) and hydroxamate (ferrichrome and coprogen) siderophores is shown in Figure 1 (Andrews et al., 2003).

**Figure 1.**
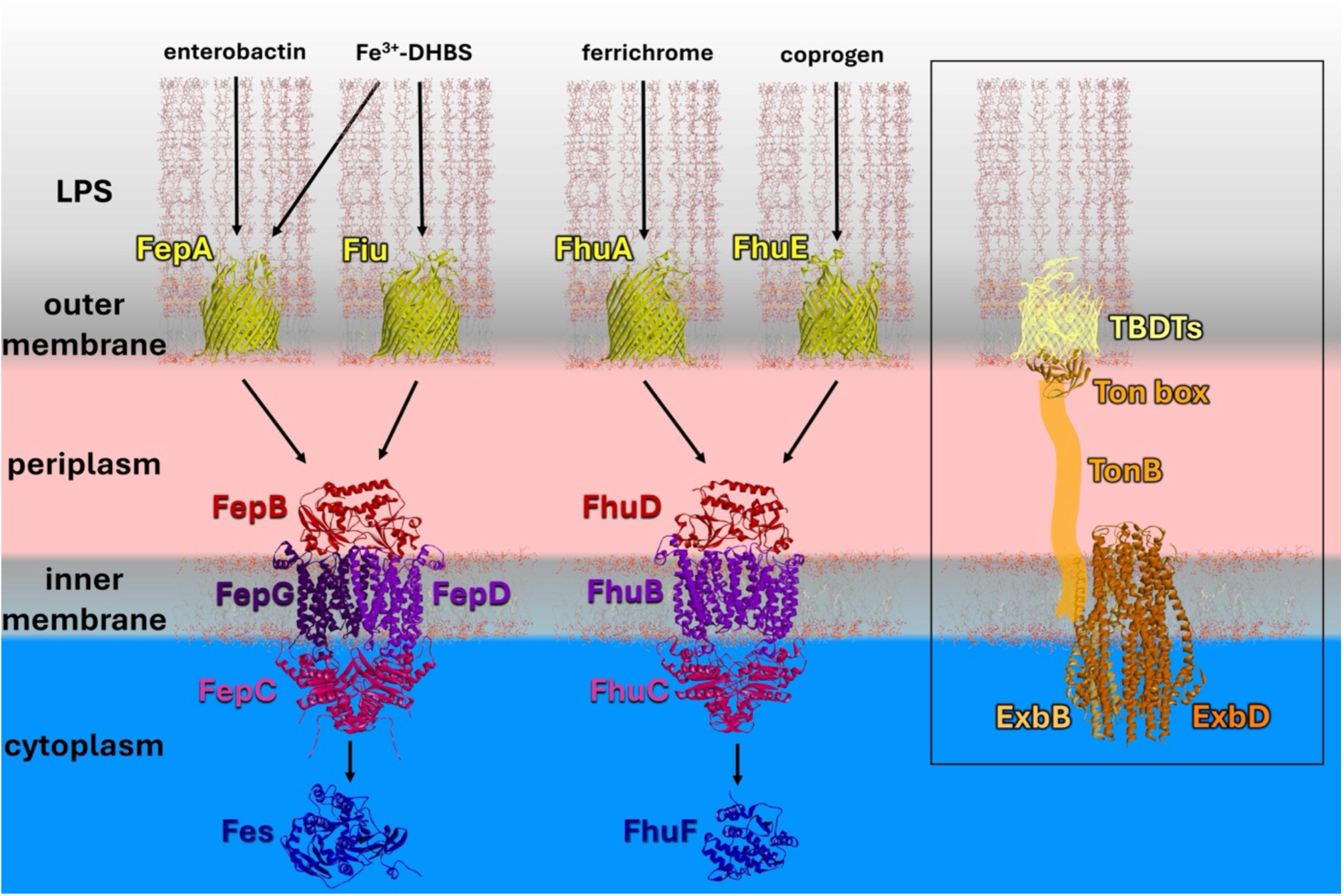
Schematic representation of the TonB-dependent *E. coli* uptake systems for catecholate and hydroxamate-type siderophores (Andrews et al., 2003). These ferric-siderophores selectively bind to outer-membrane receptors (TBDTs: FepA, Fiu, FhuA, or FhuE). Binding induces unfolding of the Ton-box, triggering the TonB-ExbB-ExbD complex, which transduces the energy generated from the proton motive force for active transport across the outer membrane. Once transported into the periplasm, the siderophore binds to a periplasmic binding protein (FepB or FhuD), which interacts with the ABC transporter complex (FepCDG or FhuBC) for translocation across the inner membrane. Within the cytoplasm, enzymes, such as esterase (Fes) and a reductase (FhuF), process ferric siderophores to reduce Fe^3+^ and release Fe^2+^ for metabolic processes. The boxed panel depicts the interaction between the outer-membrane TBDT receptor and the TonB-ExbB-ExbD complex common to all such receptors. Structures were derived from the PDB (FepA, PDB ID: 1fep (Deisenhofer et al., 1999); Fiu, PDB ID: 6bpn (Grinter & Lithgow, 2019b); FhuA, PDB ID: 1by3 (Locher et al., 1998); FhuE, PDB ID: 6e4v (Grinter & Lithgow, 2019a); FhuBCD, PDB ID: 7lb8 (Hu & Zheng, 2021); FhuF, PDB ID: 7qp5 (Trindade et al., 2023); and ExbB-ExbD, PDB ID: 6tyi (Celia et al., 2019)) or were modeled in AlphaFold3 (Abramson et al., 2024) (FepBCG, Fes, and the TBDT-Ton-box complex). The intrinsically disordered periplasmic region of TonB is depicted as a schematic tube. Proteins were embedded in membranes using CHARMM-GUI (Jo et al., 2008).

Naturally occurring sideromycins - antibiotics linked to siderophores - provided an early proof of concept for siderophore-mediated drug delivery (reviewed in (Miao et al., 2025)). This concept was further adapted for the uptake of antibiotics (Brochu et al., 1992; A. Ghosh et al., 1996) and resulted in the FDA-approved cefiderocol, a conjugate of cephalosporin with a catecholate-type siderophore (Aoki et al., 2018; El-Lababidi & Rizk, 2020; Liu et al., 2023). We previously applied hydroxamate-type tripeptide siderophores based on *N*^δ^- acetyl-*N*^δ^-hydroxy-L-ornithines for antisense PNA delivery into *E. coli* (Tsylents et al., 2024). We confirmed their iron-binding properties and TonB-dependent uptake pathway and showed that the linear siderophore transported PNA into *E. coli* mutant cells with continuous iron uptake. However, the siderophore-PNA conjugates targeting the *acpP* gene did not inhibit wild-type *E. coli* growth, probably because insufficient number of PNA molecules reached the cytoplasm. More recently, a synthetic tris-catecholate siderophore mimic was conjugated to PNA and showed a concentration-dependent growth inhibition of both *E. coli* and *Acinetobacter baumannii* (Pals, Wijnberg, et al., 2024).

Encouraged by these results confirming that synthetic siderophores can carry PNA into cells via the TonB-dependent pathway, in this study, we developed new siderophore-based carriers. We synthesized new hydroxamate-type siderophores, including a flexible analog of our previous tri-peptide with incorporated glycine residues and marine siderophores (moanachelins) containing glycine and alanine. We analyzed their iron-binding properties and compared them to natural siderophores (coprogen and ferrichrome), assessed their interactions with TBDT receptors, and, importantly, confirmed their potential as PNA carriers.

## 2. RESULTS AND DISCUSSION

### 2.1 SIDEROPHORE SELECTION, DESIGN AND SYNTHESIS

To develop new siderophore-based PNA carriers that can be synthesized on solid phase like peptides, we investigated three new siderophores. We previously used the *N*^δ^-acetyl-*N*^δ^-hydroxy-L-ornithine trimer as a siderophore mimic (S_L_, Figure 2). However, its carrier properties were insufficient to observe an effective antisense effect of PNA in wild-type *E. coli* (Tsylents et al., 2024). Our molecular dynamics (MD) simulations of S_L_ indicated that only two of the three hydroxamate groups preferentially chelated iron(III) (Tsylents et al., 2024). We attributed this to backbone stiffness, which prevented the hydroxamate groups from freely rotating to efficiently coordinate iron(III). Therefore, in this study, to enhance flexibility and improve iron(III) coordination, we modified the S_L_ siderophore by inserting glycines between the hydroxamate-bearing modified ornithines. This modification was intended to increase backbone flexibility, consistent with the residual flexibility scale (Huang & Nau, 2003). This resulted in the S_GLY_ siderophore mimic shown in Figure 2. We inserted only single glycines between the modified ornithines to minimize the number of backbone oxygens, which could compete with hydroxamate groups for iron binding. We noted a similar separation between iron-binding residues by either Gly or Ala in natural siderophores produced by *Vibrio* species, known as moanachelins (Gauglitz & Butler, 2013). Interestingly, Gly enhances the conformational flexibility of peptides, whereas Ala promotes more ordered structures (Myers et al., 1997; Pace & Scholtz, 1998). Although only one amino acid is present between the iron-binding moieties in moanachelins, we aimed to understand the functional differences between siderophores containing Gly or Ala. The structures (Gauglitz & Butler, 2013) and synthesis (Cherkupally et al., 2015) of moanachelins have been studied, but their functionality and potential applications as carriers remain unexplored.

**Figure 2.**
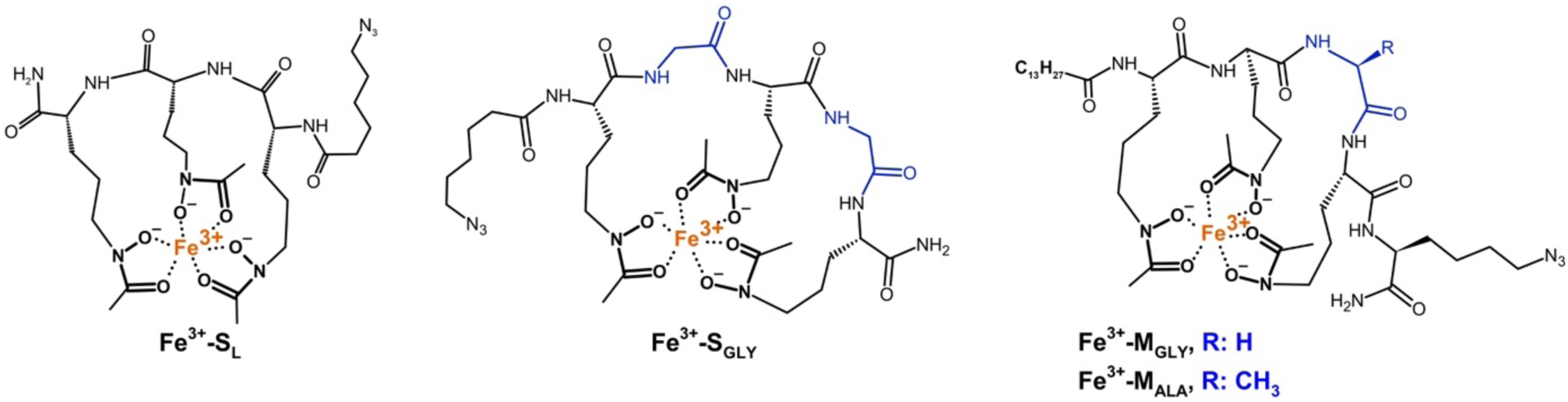
Schematic structures of iron(III) complexes with the siderophores and siderophore mimics studied in this work, with their corresponding names. The azide (N3) group, to which a PNA oligomer was attached, was introduced either at the N-terminus (for SL and SGLY), as 6-azidohexanoic acid (N3-Hx-OH), or at the C-terminus (for MGLY and MALA), as an additional azido-lysine, due to the presence of a fatty acid (-C13H27; tetradecanoic acid residue, TDA) at their N-terminus. Gly and Ala residues are shown in dark blue.

### 2.2. IRON(III) BINDING BY SIDEROPHORES

#### Verifying iron coordination by siderophores

Upon forming a siderophore-iron(III) complex, three hydroxamate groups can adopt one of two possible optical isomeric configurations – Δ or Λ (van der Helm et al., 1980). Specific isomers are recognized by the bacterial receptors responsible for siderophore transport (Raymond et al., 2015; Thulasiraman et al., 1998; Winkelmann, 1979). The predominant isomeric configuration of a siderophore in the presence of Fe^3+^ ions can be determined using circular dichroism (CD) spectroscopy. The CD approach was first applied to the natural hydroxamate siderophore, ferrichrome, which adopts the Λ configuration (van der Helm et al., 1980) (Figure 3). In CD spectra, a negative band preceding a positive band, in the 350-490 nm range, signifies the Λ configuration. Conversely, since the Δ configuration is a mirror image of the Λ configuration, the positive band appears before the negative one. As an example of a Δ-isomer structure, we used the natural hydroxamate siderophore, coprogen, derived from *Neurospora crassa* (Tóth et al., 2009; Wong et al., 1983) (Figure 3).

**Figure 3.**
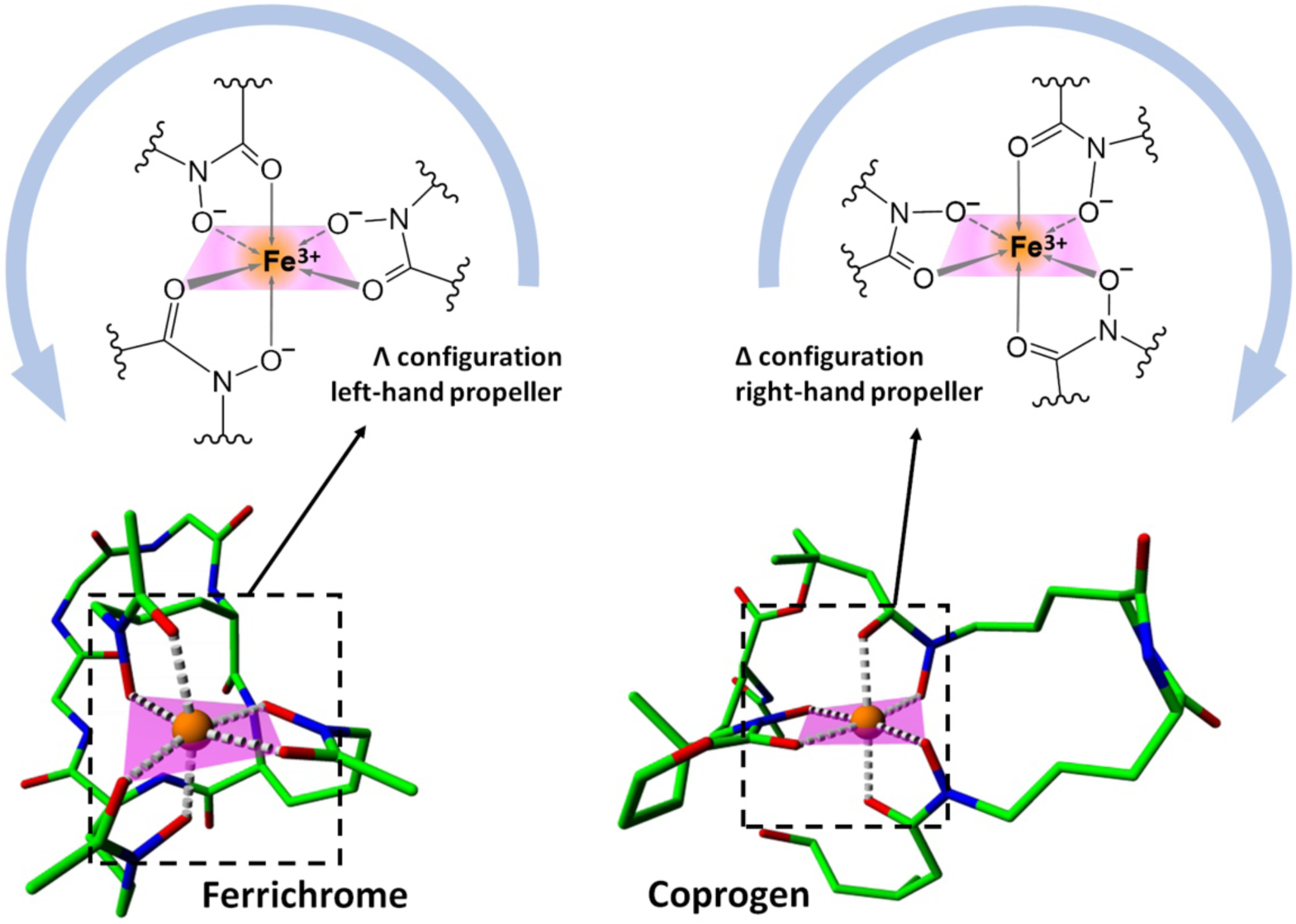
Structures of Λ and Δ stereoisomers formed by the natural siderophores, ferrichrome and coprogen, showing the hydroxamate groups coordinating Fe^3+^ (orange). The crystal structure of ferrichrome was obtained from the CCDC database (ID: 1154895; van der Helm et al., 1980), and the coprogen structure was adapted from the neocoprogen I crystal structure (CCDC ID: 1128509; Hossain et al., 1987).

While ferrichrome, coprogen, S_L_, and S_GLY_ were soluble in water, M_GLY_ and M_ALA_ solutions had to be prepared in acetonitrile. In the presence of Fe^3+^ ions, the siderophore solutions changed from colorless to yellow (for ferrichrome, S_L_, S_GLY_, M_GLY_, and M_ALA_) or orange (for coprogen) within several minutes, which indicates the formation of ferric-siderophore complexes.

CD spectra for all siderophore solutions (except coprogen) show a negative band followed by a positive band (Figure 4), indicating that their ferric complexes adopt the Λ configuration. As expected, coprogen adopts the Δ configuration, with a maximum at 376 nm and a minimum at 469 nm (Figure 4, Supplementary Table S1), consistent with previous reports (Wong et al., 1983). Notably, the Fe^3+^-S_GLY_ complex showed more pronounced extrema compared to Fe^3+^-S_L_, with its maximum shifted from 458 nm for Fe^3+^-S_L_ to 466 nm for Fe^3+^-S_GLY_ (Figure 4, Supplementary Table S1). This red shift suggests that all three hydroxamate groups in S_GLY_ participate in iron chelation, leading to improved iron-binding for this Gly-extended siderophore, compared to S_L_, which contains three modified ornithines in a row.

**Figure 4.**
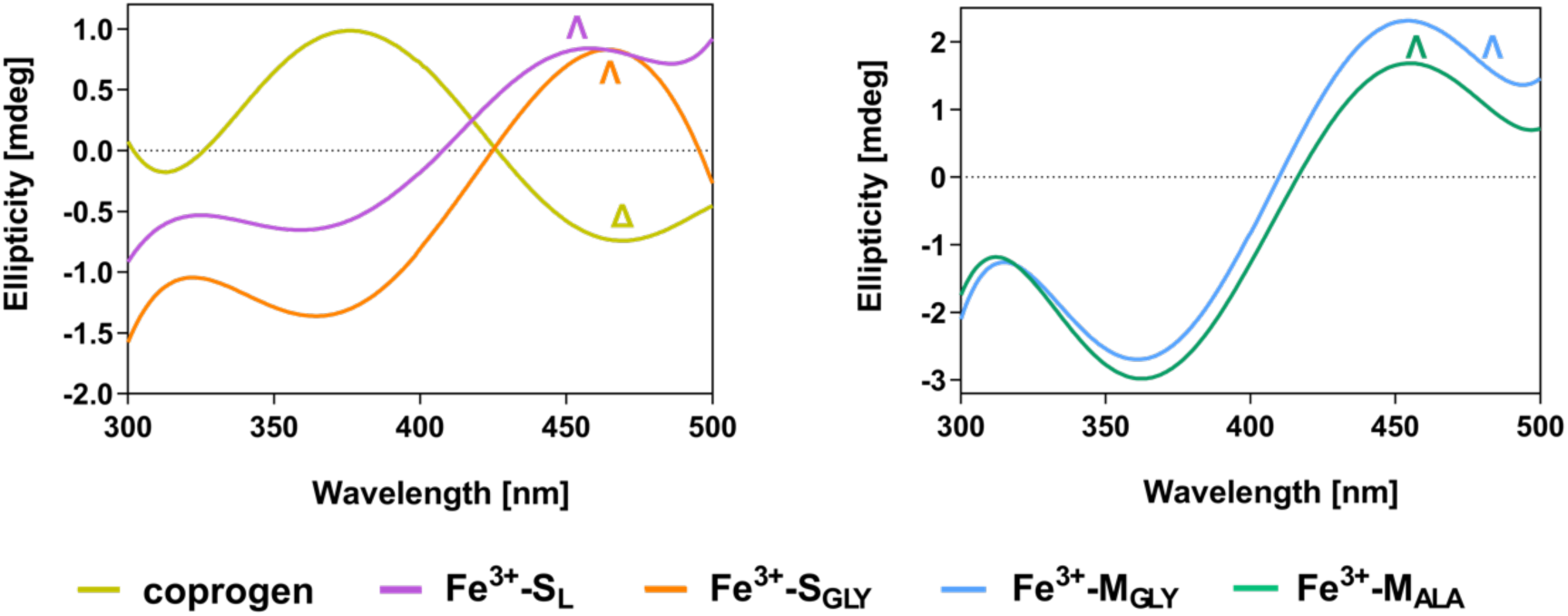
CD spectra of siderophores: left panel - coprogen, SL, and SGLY in phosphate buffer (pH 7.0); right panel - moanachelins (MGLY and MALA) in a 1:1 (v/v) mixture of phosphate buffer and acetonitrile. Spectra were recorded in the presence of the Fe^3+^ salt to assign the Λ or Δ configuration of the ferric-siderophore complexes.

#### The stability of the ferric siderophore complexes

After determining that S_GLY_ coordinates iron(III) in the Λ configuration, we assessed the stability of the resulting complex using a competition experiment with ethylenediaminetetraacetic acid (EDTA). The principle of this assay is that ferric-siderophore complexes are colored and detectable by UV-Vis spectroscopy. In particular, hydroxamate siderophores form yellow to orange ferric complexes, with maximum absorbance in the 420–440 nm range (Winkelmann, 1991). In contrast, neither colorless EDTA nor ferric-EDTA complex (colorless under these pH and concentration conditions) absorb in this UV-Vis range, which allows selective detection of ferric-siderophore complexes. This approach, described by Alvin L. Crumbliss and colleagues (Crumbliss & Harrington, 2009; Mies et al., 2008), relies on the equilibrium reaction

[Fe^3+^ − Sid] + [EDTA] ⇌ [Fe^3+^ − EDTA] + [Sid],

where the species concentrations are: [Fe^3+^-Sid] is the ferric-siderophore complex; [EDTA] is the colorless competitor; [Fe^3+^-EDTA] is the colorless ferric-EDTA complex; and [Sid] is the colorless iron-free siderophore. Upon adding EDTA, it competes with the siderophore for Fe^3+^ ions, partially removing them from the siderophore complex, which leads to color fading and a decreased absorbance for [Fe^3+^-Sid]. A siderophore with stronger iron-binding affinity will retain more Fe^3+^ ions, with absorbance decreasing only slightly upon EDTA addition (and only a subtle color change). The absorbance changes are related to the stability of the ferric-siderophore complex and indicate either weak or strong iron-binding affinity.

The reaction described above reached equilibrium only after 2-3 days. Therefore, the concentration of the ferric-siderophore complex was measured by its absorbance at λ_max_ before the addition of EDTA ([Fe^3+^-Sid]_before_) and after the EDTA incubation period ([Fe^3+^-Sid]_after_). The percentage of the remaining complex [Fe^3+^-Sid]_remaining_ was calculated using the formula:

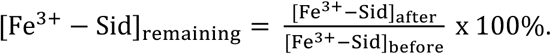

Using this method, we compared the relative stability of the Fe^3+^-S_GLY_ complex to that of Fe^3+^-S_L_ and the natural siderophores in the presence of EDTA (Figure 5). Unfortunately, the limited solubility of M_GLY_ and M_ALA_ excluded these siderophores from this experiment.

**Figure 5.**
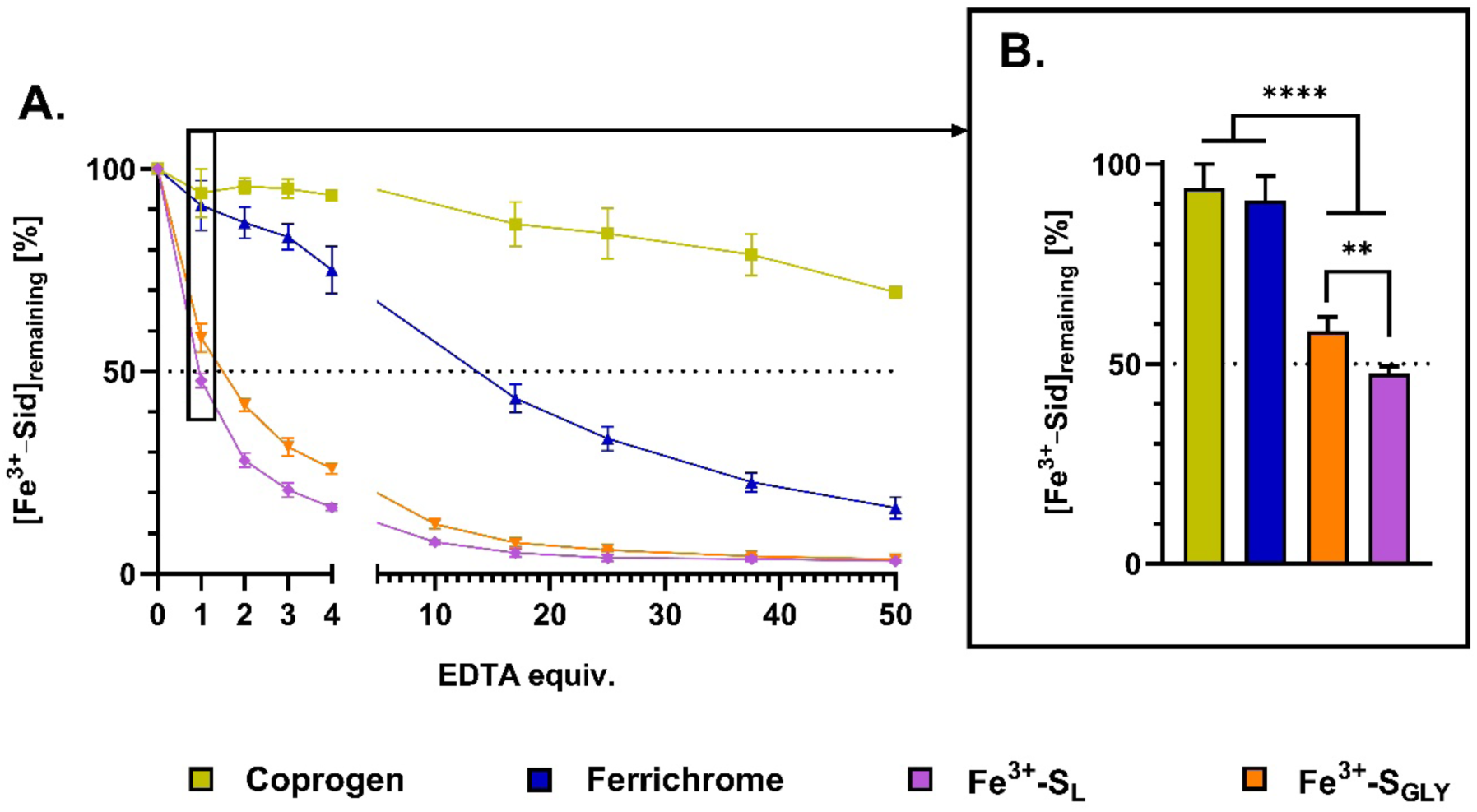
Relative stability of ferric-siderophore complexes determined by spectrophotometric competition experiments. (A) Complexes were incubated with concentrations ranging from 0 to 50 molar equivalents of EDTA; (B) Complexes were incubated with an equimolar (1 equiv.) concentration of EDTA. Solutions were prepared in phosphate buffer (pH 7.4). The data show the mean ± standard deviation (n = 3, each in duplicate). Statistical significance was calculated using a one-way ANOVA and is shown as ** p < 0.01 and **** p < 0.0001.

Based on the EDTA competition assay, the natural siderophores coprogen and ferrichrome are more stable than the synthetic siderophores S_GLY_ and S_L_. When incubated with an equimolar amount of EDTA, about 90% of the ferric-coprogen and ferric-ferrichrome complexes remain (Figure 5B). At higher EDTA concentrations, the amount of the ferric-ferrichrome complex drops to about 20%, while that of the ferric-coprogen complex remains at about 75%, corroborating that coprogen is a stronger iron chelator than ferrichrome (Figure 5A) (Crumbliss & Harrington, 2009).

In the presence of 1 equivalent of EDTA, the Fe^3+^-S_GLY_ complex retained about 60% of its concentration, while the Fe^3+^-S_L_ complex retained less than 50% (Figure 5B), indicating that S_GLY_, unlike S_L_, binds Fe^3+^ more strongly than EDTA. This difference in Fe^3+^ binding affinity between S_GLY_ and S_L_ persists as the concentration of EDTA increases (Figure 5A).

### 2.3. SIDEROPHORE UPTAKE PATHWAY

After confirming that S_GLY_, M_GLY_, and M_ALA_ bind iron(III), we tested whether their ferric complexes are recognized and internalized by receptors of the *E. coli* TonB-dependent system (Silale & Van Den Berg, 2023). Note, that moanachelins are xenosiderophores for *E. coli* as they are produced by *Vibrio* species. Their uptake by *E. coli* has not been studied, and therefore we do not know whether they can be taken up and used by *E. coli* strains. As shown in Figure 1, each TBDT pathway consists of at least one outer membrane receptor, periplasmic binding protein, inner-membrane transporter, and cytoplasmic reductase (Andrews et al., 2003). To investigate uptake, we used *E. coli* mutants with deletions in the catecholate uptake system, ensuring that any observed siderophore uptake occurs exclusively via the hydroxamate pathway (Table 1). Mutant strains were cultured in iron-limited medium, which hindered bacterial growth. The growth of cultures with and without the S_GLY_, M_GLY_, or M_ALA_ siderophores was compared to assess their uptake and subsequent iron(II) release in the cytoplasm, which is necessary for cell growth. Effective recognition and internalization of these siderophores by TBDT receptors should restore bacterial growth compared to cultures lacking the siderophores.

**Table 1.**
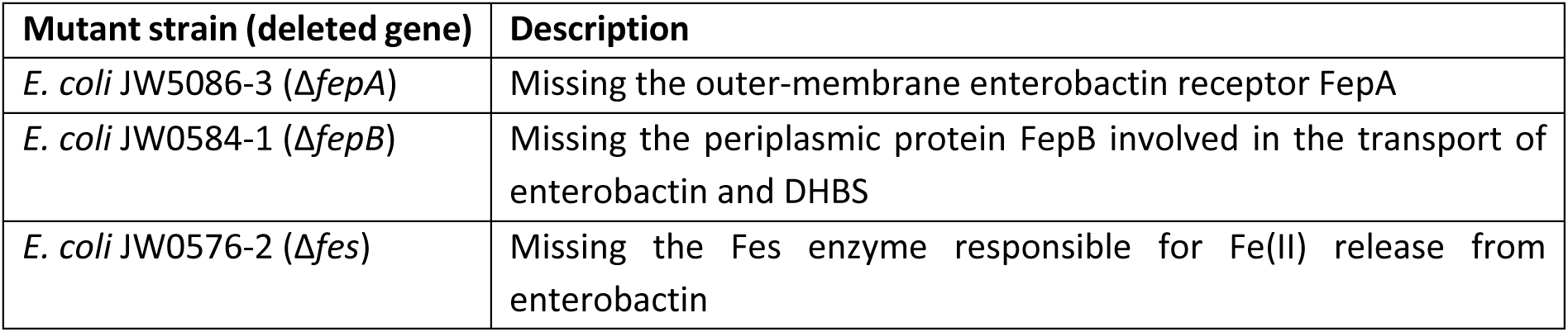
Bacterial strains used in growth recovery experiments.

For all *E. coli* mutants of the catecholate pathway, supplemented with S_GLY_, we observed statistically significant growth recovery (Figure 6A). These results suggest that S_GLY_ is recognized and transported by the TBDT hydroxamate-pathway. This is consistent with our previous studies on S_L_, which required the coprogen outer-membrane receptor for cellular entry (Tsylents et al., 2024). In contrast, growth recovery was less pronounced in cultures treated with the moanachelin analogs M_GLY_ and M_ALA_ (Figure 6B). Specifically, M_ALA_ supplementation resulted in a statistically significant growth increase only in the mutant lacking the FepA receptor, whereas M_GLY_ promoted growth in the mutants lacking the FepB and Fes proteins. These results suggest that M_GLY_ may be more effectively recognized and internalized by *E. coli* than M_ALA_, but the uptake efficiency for both M_GLY_ and M_ALA_ siderophores remains inconclusive. These differences in M_GLY_ and M_ALA_ uptake arise from a single amino acid substitution (Gly vs Ala), which underscores the high specificity of the TBDT receptors. Notably, M_GLY_ and M_ALA_ exhibit limited solubility and were dissolved in DMSO before use. The final DMSO concentration in the cultures was maintained at 3%, which may have influenced the results, preventing direct comparisons with the water-soluble S_GLY_.

**Figure 6.**
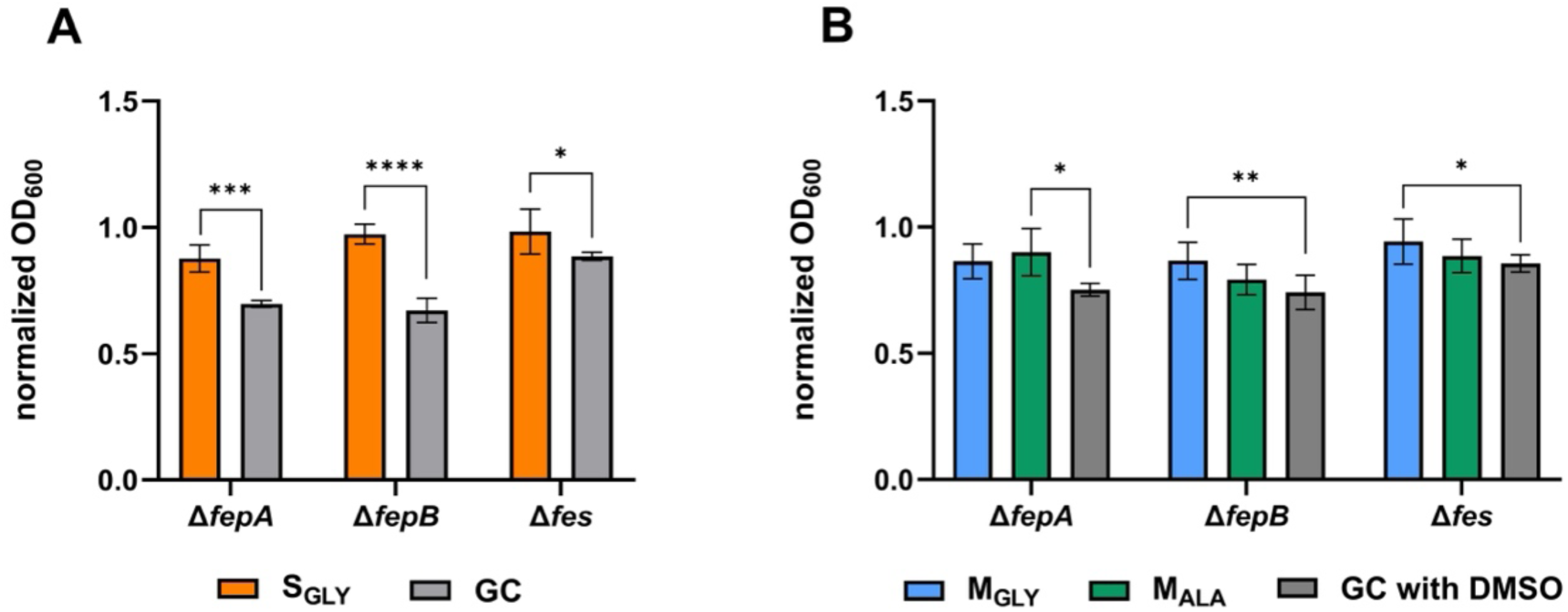
Growth of *E. coli* mutant strains incubated with (A) SGLY and (B) MGLY or MALA. *E. coli* mutants were cultured under iron-limiting conditions in the absence or presence of 16 μM of the indicated siderophores. MGLY and MALA were dissolved in DMSO, which was included in the corresponding growth control (GC) cultures. Statistical significance between the GC and the sample was calculated using either a two-tailed *t*-test (A) or a one-way ANOVA (B): * *p* < 0.05, ** *p* < 0.01, *** *p* < 0.001 and **** *p* < 0.0001. Unmarked differences are not statistically significant. Data are shown as mean ±SD from three independent biological experiments (n = 3), each performed in duplicate.

#### *In vivo* fluorescence silencing with siderophore-PNA conjugates

After confirming that S_GLY_, M_GLY_, and M_ALA_ are recognized by *E. coli*, we investigated their potential as PNA carriers. We conjugated them to a PNA sequence (PNA_anti-*rfp*_) designed to hybridize with a fragment of the *mrfp1* gene transcript encoding red fluorescent protein (RFP) from a plasmid introduced to *E. coli* cells (Figure 7A) (Równicki et al., 2017). If a siderophore-PNA conjugate efficiently delivers PNA_anti-*rfp*_ into *E. coli* cells expressing RFP, the PNA would bind to the target mRNA, decreasing RFP production and, ultimately, its fluorescence. As a control, we used a scrambled PNA sequence (PNA_SCR_) with 7 mismatches relative to the targeted mRNA fragment (Figure 7B). The PNA conjugates were synthesized via the copper-catalyzed azide-alkyne cycloaddition (CuAAC) reaction (Figure 7C) (Rostovtsev et al., 2002; Tornøe et al., 2002; Zhu et al., 2016).

**Figure 7.**
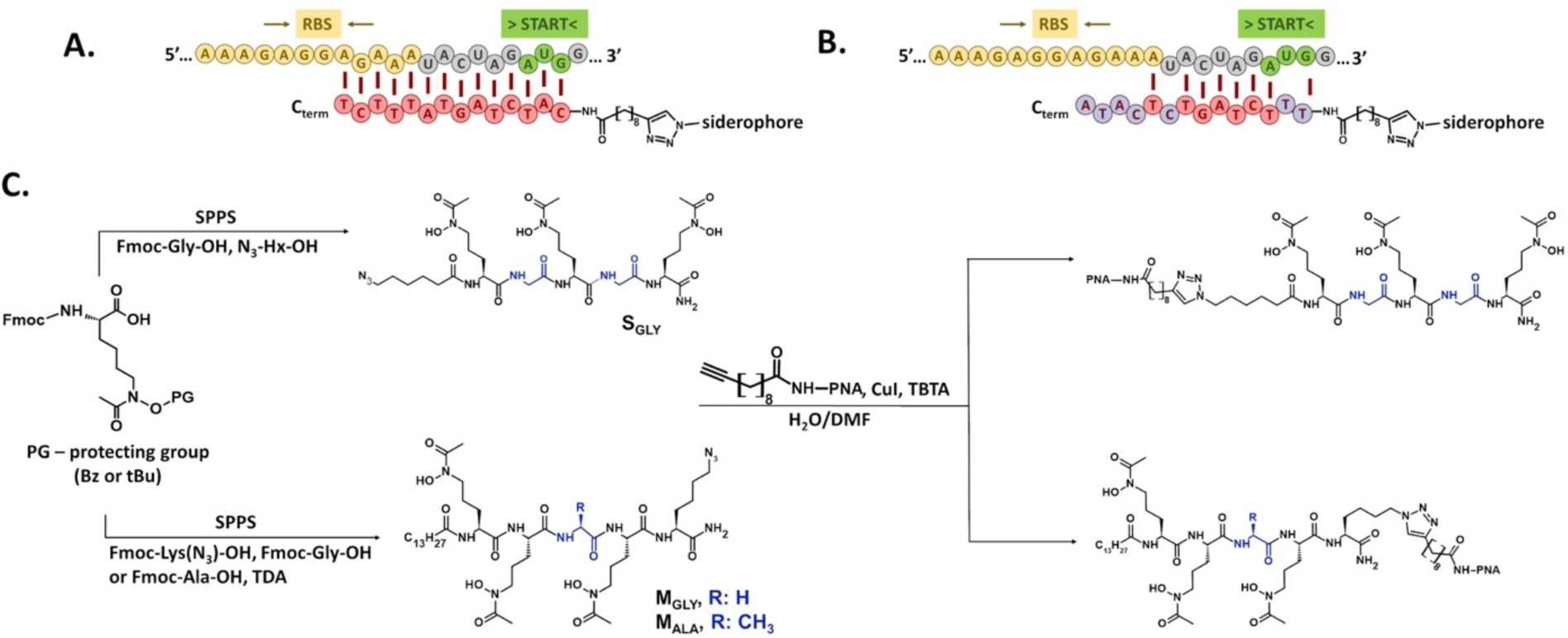
Schematic representation of the *mrfp1* mRNA sequence targeted by (A) PNAanti-*rfp* and (B) PNASCR sequences. Matching PNA nucleobases are shown in red, mismatched in blue. The ribosome binding site (RBS, in green) and start codon (START, in yellow) of the mRNA are shown. (C) The schematic pathway of the siderophore-PNA synthesis. N3-Hx-OH: 6-azidohexanoic acid; Fmoc-Lys(N3)-OH: azido-lysine; TDA: tetradecanoic acid; SPPS; solid-phase peptide synthesis; CuI; copper(I) iodide; TBTA: tris[(1-benzyl-1H-1,2,3-triazol-4-yl)methyl]amine.

To detect fluorescence changes induced by siderophore-PNA conjugates, we used the protocols established in our previous studies (Równicki et al., 2017; Tsylents et al., 2024). Bacteria expressing RFP were cultured under iron-limiting conditions and incubated with iron-preloaded siderophores and their PNA conjugates. As we had not previously observed significant RFP silencing in wild-type *E. coli*, we used an *E. coli* Δ*fur* mutant. This strain lacks the Fur regulator, which is the transcriptional repressor of genes responsible for expression of siderophore receptors, resulting in unlimited iron uptake.

The RFP fluorescence intensity measured for bacteria exposed to the siderophore-PNA conjugates is shown in Figure 8. For all three PNA_anti-*rfp*_ conjugates, which are fully complementary to the mRNA target, we observed a statistically significant drop in fluorescence intensity. Treatment with M_GLY_-PNA_anti-*rfp*_ and M_ALA_- PNA_anti-*rfp*_ reduced fluorescence to the levels comparable to that of the negative control (bacteria without the plasmid encoding RFP). The conjugates with PNA_SCR_, resulted in a less pronounced reduction of fluorescence (to ∼75% of the normalized control). This partial silencing by the PNA_SCR_ conjugates (both scrambled and mismatched PNA) can be attributed to the sequence containing 7 bases complementary to the mRNA target, including a 6-base stretch that overlaps with one base of the start codon (Figure 7B). Therefore, PNA_SCR_ may still bind RNA tightly (Katkevics et al., 2024) and exert some silencing effect. Nevertheless, the silencing efficacy of the siderophore-PNA_anti-*rfp*_ conjugates, which are fully complementary to the target mRNA fragment, was significantly greater than that of their corresponding PNA_SCR_ controls, indicating sequence-specific silencing.

**Figure 8.**
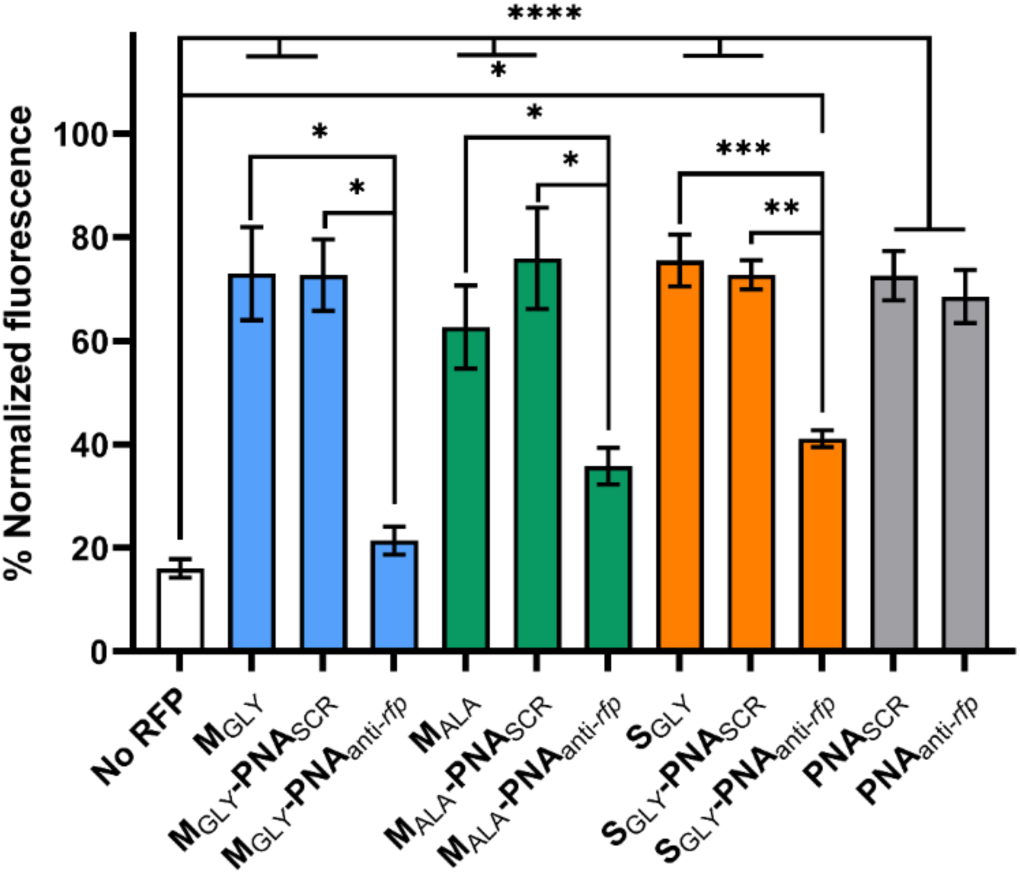
RFP silencing by siderophore-PNA conjugates in an *E. coli* Δ*fur* mutant. Bacteria were cultured in iron-limiting conditions and treated with 16 μM of the indicated compounds. RFP fluorescence was normalized to the fluorescence intensity recorded for untreated bacteria carrying a plasmid expressing RFP. Data are shown as the mean ± SEM from three independent experiments (n = 3). The “No RFP” bar denotes cells lacking the RFP-encoding plasmid. Statistical significance was determined by a one-way ANOVA (**** *p* < 0.0001, *** *p* < 0.001, \*\**p* < 0.01, \**p* < 0.05). Unmarked differences are not statistically significant.

To further confirm the uptake of siderophore-PNA conjugates and directly observe fluorescence silencing in single bacterial cells, we performed confocal microscopy imaging (Supplementary Figure S1). The images show a reduced RFP signal in *E. coli* cells treated with the conjugates. The calculated fluorescence levels in bacteria incubated with M_GLY_-PNA_anti-*rfp*_ (about 20%) and M_ALA_-PNA_anti-*rfp*_ (about 40%) are consistent with the results of the RFP assay shown in Figure 8. In confocal imaging, we also observed a substantial decrease in RFP fluorescence in bacteria treated with S_GLY_-PNA_anti-*rfp*_ (about 20%). Overall, results from both the plate reader assay and microscopy confirm that the conjugates with PNA_anti-*rfp*_ are recognized and transported into *E. coli* cells, leading to the expected gene silencing effect.

### 2.4. SIDEROPHORE FLEXIBILITY FROM MOLECULAR DYNAMICS

We experimentally found that extending the siderophore backbone with glycine improves iron binding and PNA delivery. Therefore, to investigate whether this modification increased backbone flexibility and facilitated iron(III) coordination, we applied all-atom molecular dynamics (MD) simulations. Trajectory-derived root-mean square fluctuations (RMSF) of free S_L_ and S_GLY_ indicate that separation of *N*^δ^–acetyl–*N*^δ^– hydroxy–L–ornithines (AHOs) by Gly increases the mobility of the hydroxamate groups (Figure 9). RMSF values are by 0.4, 1.2 and 0.8 Å higher for the S_GLY_ AHO(1), AHO(2) and AHO(3) residues, respectively, than the RMSF of the corresponding groups in S_L_. Also, the average RMSF is higher for S_GLY_ (3.0 ± 0.3 Å) than for S_L_ (2.3 ± 0.1 Å) and the differences are statistically significant with p-value < 0.05 (for per-atom RMSF, see Supplementary Figure S2). These differences in flexibility explain the CD results showing that S_GLY_ binds iron(III) better than S_L_.

**Figure 9.**
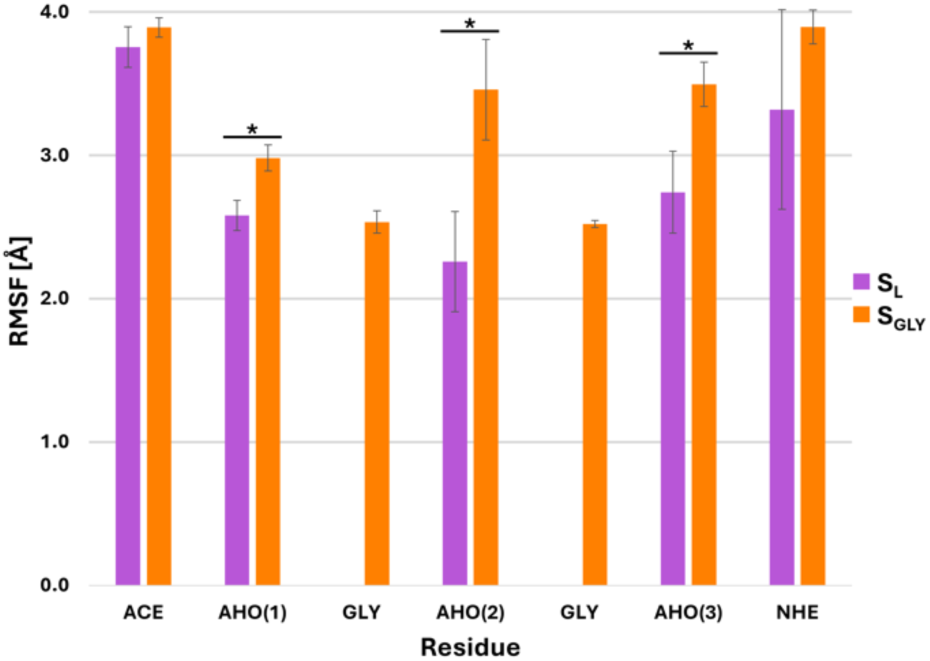
RMSF of SL and SGLY residues derived from three 200 ns trajectories in the absence of iron(III). Mean values ± SEM are shown. Statistical significance was calculated using one-tailed t-test assuming two-sample equal variance: * p < 0.05. AHO stands for *N*^δ^–acetyl–*N*^δ^–hydroxy–L–ornithines, numbered from N- to C-terminus, ACE is acetyl group at the N-terminus, and NHE is amide group at the C-terminus (Supplementary Figure S3).

To generate the ferric-siderophore complexes and to recreate the dative bonds between iron(III) and oxygen atoms in these siderophores, for the interactions involving iron(III), we applied the modified 12-6-4 Lennard-Jones (LJ 12-6-4) potential (Li et al., 2015; Li & Merz, 2014) with the following form:

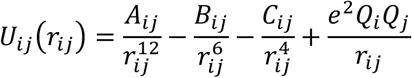

where *U_ij_* is the interaction energy of atoms *i* and *j* dependent on the distance *r*, *A*, *B*, and *C* are atom-type specific coefficients assigned to each atom pair. Terms in power -12 and -6 correspond to repulsive and attractive pairwise interactions as in ordinary Lennard-Jones potential (LJ 12-6), and the 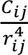 term represents the charge-induced dipole interaction. The 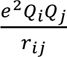 is a Coulombic term representing the electrostatic interactions between atoms, with 𝑄_𝑖_and 𝑄_j_ partial charges. MD simulations using the standard LJ 12-6 potential often do not reproduce the interactions of charged metal ions, *e.g.*, iron(III), with organic molecules. Therefore, to address this issue, the LJ 12-6-4 potential was developed with the aim of accurately reproducing hydration free energies, ion-oxygen distances, and coordination number with a single set of parameters (see Supplementary Section S1) (Li et al., 2015; Li & Merz, 2014).

In the first iron binding simulations using the modified LJ 12-6-4 potential, the iron(III) ion was placed at a distance at least 16 Å away from the deferred siderophores (Supplementary Figure S4). Using the ferrichrome structure (CCDC ID: 1154895; van der Helm et al., 1980) as a starting conformation, we tested multiple sets of *C_ij_* parameters of the LJ 12-6-4 potential against the experimental free energy of binding of iron(III) to ferrichrome (Anderegg et al., 1963). However, even the parameter sets with favorable binding free energies obtained via thermodynamic integration (TI) (see Methods), compared to the experimental value, failed to

reproduce the proper geometry of iron(III) coordination sphere. One or more hydroxamate groups would twist, and the coordination number of iron(III) extended to seven due to water molecules interacting with the cation (Supplementary Figure S5). We thus chose *C_ij_* parameters that reproduced both the free energy, derived from the experimental association constants, and geometry (Supplementary Table S2). MD simulations with the LJ 12-6-4 nonbonded potential and this parameter set were carried out for S_L_, S_GLY_, M_GLY_, and M_ALA_. Trajectories showed iron(III) association and binding by three hydroxamate groups of M_ALA_ in two (out of three) replicas, and in one replica for S_L_, S_GLY_, and M_GLY_ (Figure 10).

**Figure 10.**
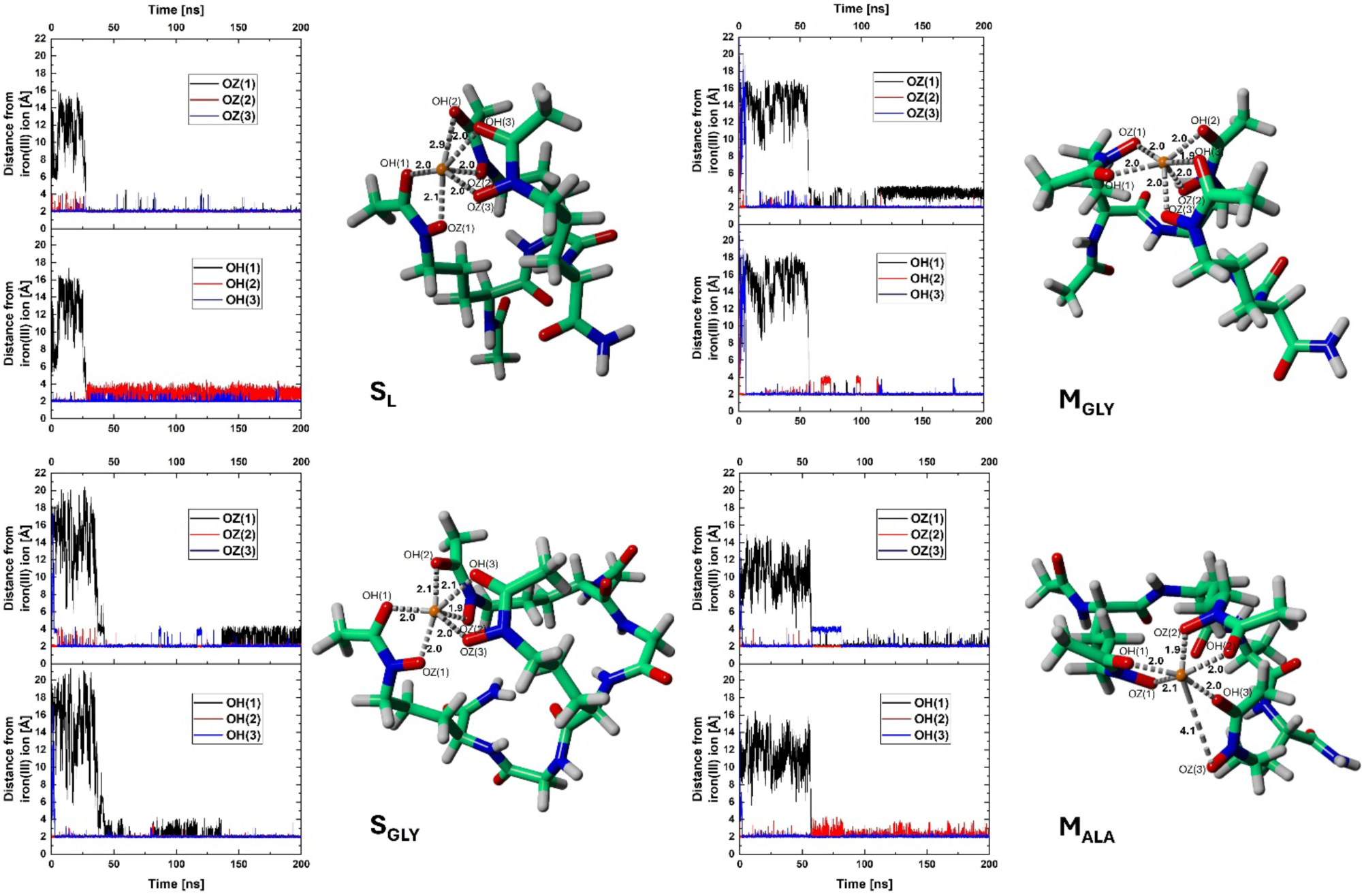
Distances between iron(III) and the oxygen atoms of hydroxamate groups from MD trajectories, with representative structures showing the six oxygen atoms coordinating the iron(III) ion. OH and OZ denote the hydroxamate oxygens, with numbers in parentheses denoting the AHO residues to which they belong (from N- to C-terminus). Siderophores are shown in a stick representation with iron(III) as an orange sphere. Distances are in Å. The aliphatic tails of MGLY and MALA are omitted (see Methods).

Notably, the S_L_ structure obtained from the simulations applying the LJ 12-6-4 nonbonded model has the iron(III) ion well-coordinated by all three hydroxamate groups of the siderophore. This lies in contrast to our previous results, where preferentially only two hydroxamate groups chelated iron(III) (Tsylents et al., 2024). This difference can be explained by more extensive parametrization of the system performed in the present work, namely the use of LJ 12-6-4 potential. Previously, we applied the standard LJ 12-6 form. However, weaker chelation of iron(III) by S_L_ in our previous simulations made us extend its side chain with glycines and develop S_GLY_. The latter siderophore mimic turned out to have improved performance in CD and EDTA competition assays, as well as molecular docking.

Selected representative structures from the LJ 12-6-4 nonbonded simulations served as starting points for MD simulations with covalently bound iron(III), in the ’bonded model’, with harmonic bonds applied between iron(III) and hydroxamate oxygens (see Methods). The representative structure from the most populated cluster of the latter bonded model served as an input structure for molecular docking (see next subsection).

### 2.5 PREDICTIONS OF THE COMPLEXES BETWEEN SIDEROPHORES AND RECEPTORS OF THE TONB-DEPENDENT PATHWAY

Growth recovery assays and siderophore-PNA uptake experiments on *E. coli* mutant strains indicated that S_L_ (Tsylents et al., 2024), S_GLY_, M_ALA_, and M_GLY_ siderophores are taken up by *E. coli* cells through the TonB-dependent hydroxamate pathway (Figure 1). To gain insight into their interactions, we docked these siderophores to the TBDT receptors involved in the hydroxamate-uptake pathway. The iron(III)-bound conformations for docking were taken from MD trajectories and are shown in Figure 10. Control re-docking of crystallographic ligands (except for FhuF, which was deposited in the PDB only as an *apo*-structure) confirmed the correctness of the docking protocol with the ligand heavy atom root-mean-square deviations (RMSD) between the re-docked ligand and its crystallographic pose below 1.0 Å (RMSD ≤1.0 Å indicates excellent docking performance (Castro-Alvarez et al., 2017) (Supplementary Figure S6). The docking scores (a metric roughly resembling the free energy of binding of a ligand) for S_L_ and S_GLY_ are similar to those of crystallographic ligands, with the scores for S_GLY_ being more favorable, by 0.5 to 2.7 kcal/mol, depending on the receptor (Table 2). M_GLY_ and M_ALA_ consistently showed 1-2 kcal/mol worse docking scores, likely due to their hydrophobic chains (Table 2). The docking scores and binding poses indicate that the ligands can bind to proteins from the hydroxamate pathway of the *E. coli* TBDT system. S_GLY_ is the only tested siderophore that exhibited equal or higher affinity toward a protein than its *native* ligand (crystallographic coprogen in the case of FhuE or docked ferrichrome in the case of FhuF). Its docking poses reveal a number of interactions with the receptor proteins, similar to those identified within the crystal structures (Figure 11, Supplementary Figures S7-8).

**Figure 11.**
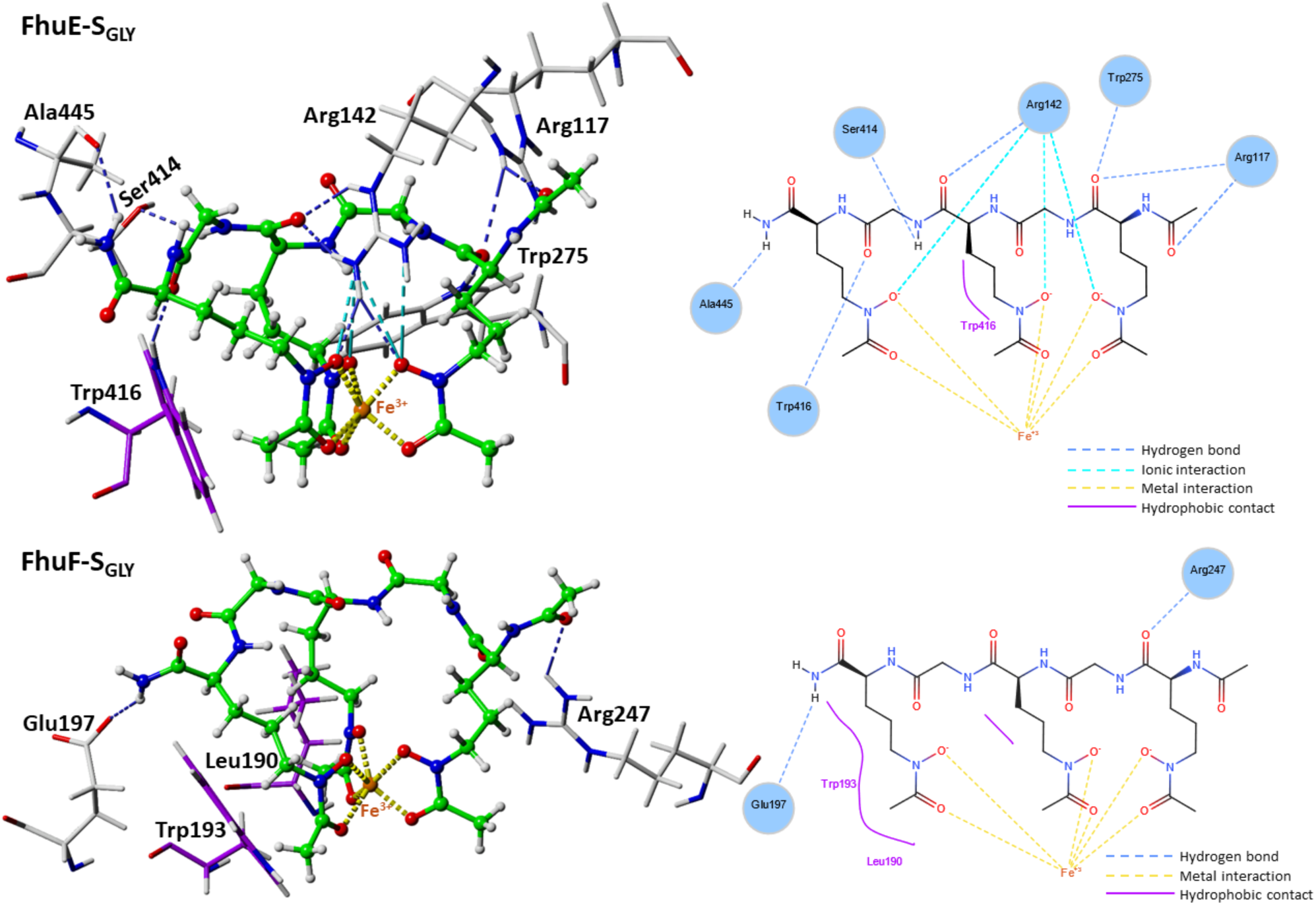
Interactions between SGLY and FhuE or FhuF in the top-scoring docking poses. Amino acids participating in hydrophobic contacts are in purple.

**Table 2.** Top-scoring docking scores of natural and synthetic siderophores docked to the crystal structures of *E. coli* proteins involved in the uptake of hydroxamate siderophores. PDB codes of the original structures are shown in parentheses. ^a^ Coprogen structure with PDB ligand ID: CPO. ^b^ Coprogen structure with PDB ligand ID: HWS.

| Target protein | Docking score for ligand [kcal/mol] |  |  |  |  |  |
| --- | --- | --- | --- | --- | --- | --- |
| | $S_L$ | $S_{GLY}$ | $M_{GLY}$ | $M_{ALA}$ | Coprogen | Ferrichrome |
| FhuA (1by5) | -11.7 | -12.5 | -9.9 | -9.0 | -12.9<sup>a</sup> | -14.5 |
| FhuE (6e4v) | -9.9 | -12.6 | -9.7 | -9.1 | -11.5<sup>b</sup> | -12.7 |
| FhuD (1esz) | -9.7 | -10.3 | -8.9 | -9.6 | -12.2<sup>a</sup> | -12.2 |
| FhuF (7qp5) | -10.3 | -11.6 | -9.6 | -8.4 | -10.8<sup>a</sup> | -11.6 |

However, other protein-siderophore docked complexes also form a similar number of contacts (Supplementary Figures S9-S15). The applied docking boxes were large enough to accommodate many diverse binding modes (Supplementary Figure S16). Despite this, we observed an overlap between receptor residues (within 4 Å of the ligand) that interact with the siderophore in the crystal structures and in docked poses. Between 23% and 100% (61% on average) of the residues near the top-scoring pose of the docked ligand are also found near the crystallographic ligand (Supplementary Figures S17-S20). The docking results, together with the experimental assays, indicate that these *E. coli* TBDTs may transport and process S_L_, S_GLY_, M_GLY_, and M_ALA_.

## 3. CONCLUSIONS

We investigated the effect of backbone flexibility of hydroxamate-type siderophores on their iron(III) binding and PNA transport into *E. coli*. All siderophores form complexes with iron in the Λ-isomeric configuration, except coprogen, which adopts a Δ-isomeric structure. Due to improved iron(III) binding, S_GLY_ CD spectrum showed a red-shifted maximum, compared to S_L_. This suggests that all three hydroxamate groups participate in iron(III) coordination, unlike in S_L_, where only two were preferentially involved. We also observed enhanced iron-binding by S_GLY_ in competition assays with EDTA. Although the iron-binding affinity of natural siderophores, coprogen and ferrichrome, remained higher, the backbone flexibility introduced into the synthetic S_GLY_ shows promise for its further optimization. CD spectra and growth recovery assays confirmed that introducing Gly into the S_GLY_ backbone did not disrupt complex formation or recognition by the TBDT receptors of *E. coli*, similar to S_L_. These findings, together with docking and MD simulations, suggest that S_GLY_ is a promising candidate for PNA delivery. While the M_GLY_ and M_ALA_ moanachelin analogs were less efficiently internalized, compared to S_GLY_, upon conjugation to PNA, these siderophores also delivered PNA into *E. coli* cells. The gene-silencing effect was confirmed for these siderophore-PNA_anti-*rfp*_ conjugates using confocal microscopy.

These findings highlight the potential of hydroxamate siderophores as carriers for antisense PNA into bacterial cells. The S_GLY_ siderophore with a flexible backbone showed improved iron-binding affinity, TBDT recognition, and PNA delivery efficiency. Although M_GLY_ and M_ALA_ were less efficiently internalized alone, when conjugated with PNA, these siderophores also enabled delivery of antisense PNA. These results demonstrate that hydroxamate siderophores, in combination with PNA, are promising candidates for further exploration in Trojan horse strategy applications against bacteria.

## 4. MATERIALS AND METHODS

### Reagents and conditions

All reagents and solvents were used as received from their respective suppliers. The Rink-amide resin (TentaGel S RAM) for peptide synthesis was obtained from Merck. The Nα-Fmoc-protected hydroxamate-ornithine derivative (Fmoc-L-Orn(Ac,OBz)-OH) was supplied by Iris Biotech GmbH. The Nα-Fmoc protected L-amino acids (Fmoc-Gly-OH, Fmoc-Ala-OH), tetradecanoic acid (TDA), iron-free ferrichrome (desferrichrome), 2,2ʹ-dipyridyl (DP), and Amberlite XAD-2 resin were obtained from Merck. Fmoc-L-Lys(N_3_)-OH and N_3_-Hx-OH were obtained from Novabiochem. Lysogeny broth (LB) and malt extract were obtained from VWR, and Mueller-Hinton broth (MHB) from Difco. Biotin and 8-hydroxyquinoline were obtained from Carl Roth. Bio-Gel P-2 resin was purchased from Bio-Rad. All chemicals were of analytical or reagent grade. Buffers were prepared using distilled water from a Direct-Q Millipore (mQ) system.

Products were analyzed by reverse-phase high-performance liquid chromatography (RP-HPLC) using Knauer C18 columns (5 μm particles, 4.6 × 250 mm for analytical and 8 × 250 mm for semi-preparative chromatography). Peptides were purified using mobile phase gradients of buffer A (0.1% TFA in acetonitrile (ACN)) and buffer B (0.1% TFA in mQ water) at room temperature, with UV-Vis detection at 220 nm, and flow rates of 1.5 mL/min (analytical) and 4.5 mL/min (semi-preparative).

### Synthesis of the tBu-protected hydroxamate-ornithine derivative: Fmoc-Orn(Ac,OtBu)-OH

Fmoc-Orn(Ac,OtBu)-OH was synthesized from the commercially available α-benzyl ester of N-Cbz-protected glutamic acid, according to literature procedures (Jiang et al., 2006; Kobayashi et al., 2012).

### Synthesis of modified hydroxamate ornithine-based siderophores

The S_GLY_ (azido-Hx-(Orn(Ac,OH)-Gly-(Orn(Ac,OH)-Gly-Orn(Ac,OH)), M_GLY_ (TDA-(Orn(Ac,OH))_2_-Gly-Orn(Ac,OH)-Lys(N_3_)) and M_ALA_ (TDA-(Orn(Ac,OH))_2_-Ala-Orn(Ac,OH)-Lys(N_3_)) siderophores were synthesized manually on the solid phase using either Fmoc/tBu or Fmoc/Bz chemistry, as described previously (Tsylents et al., 2024). For M_GLY_ and M_ALA_, TDA was attached at the N-terminus and the azide group (from azido-lysine) at the C-terminus, both using standard amino acid coupling reactions. The remaining tBu protecting groups were removed by prolonged treatment with a TFA/triisopropylsilane/water (95:2.5:2.5, v/v/v) mixture for over 6 hours. The synthesized products were confirmed by mass spectrometry (MS) using the Q-TOF Premier mass spectrometer, then lyophilized and purified (Table 3, Supplementary Figures S21–S23).

**Table 3.**
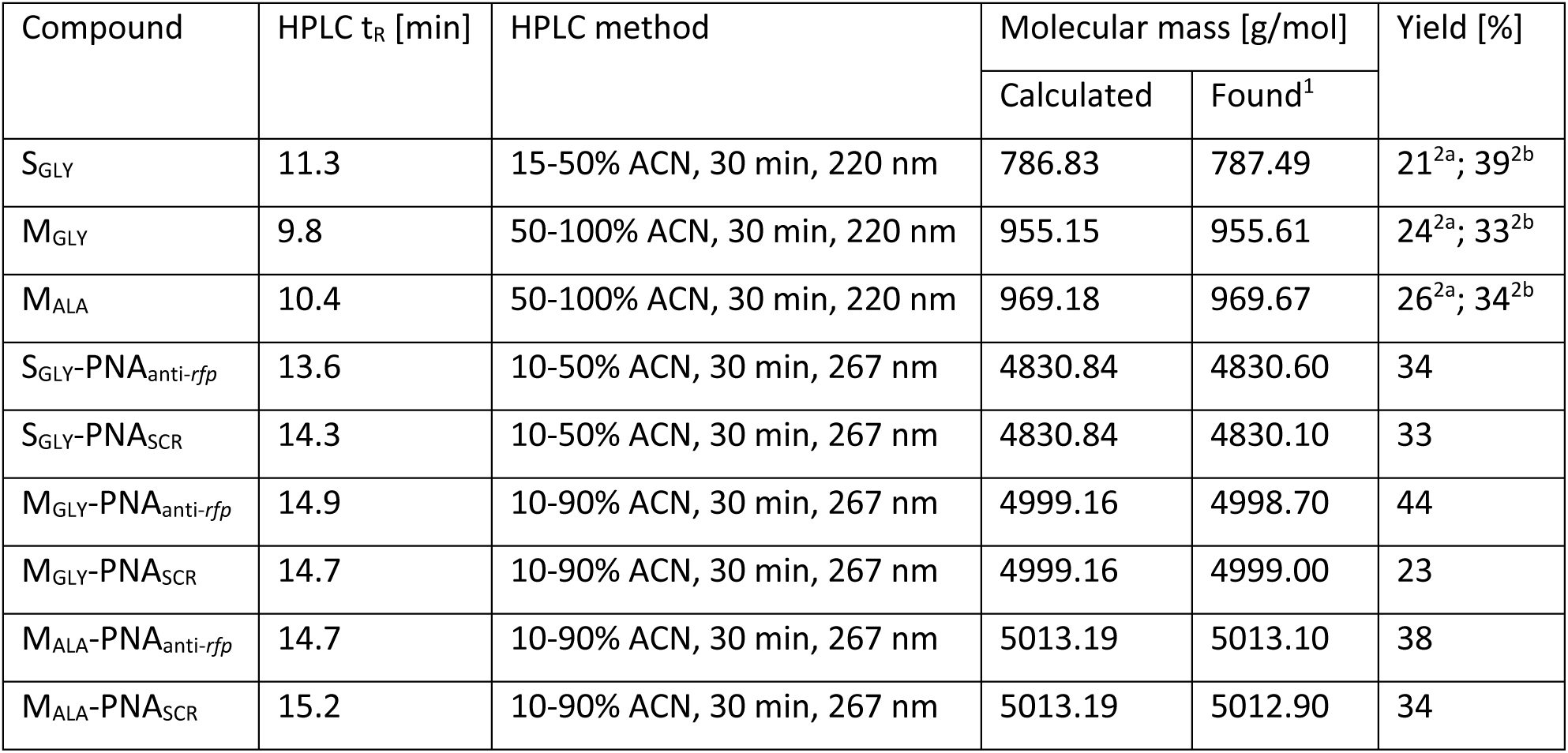
Retention times (tR) and molecular masses of synthesized siderophores and PNA-conjugates with yields calculated based on resin loading. ^1^Obtained from the Q-TOF Premier mass spectrometer. Yields for siderophores synthesized with ^2a^tBu- or ^2b^Bz-protected modified ornithine were calculated separately.

| Compound | HPLC $t_R$ [min] | HPLC method | Molecular mass [g/mol] | | Yield [%] |
| --- | --- | --- | --- | --- | --- |
|  |  |  | Calculated | Found<sup>1</sup> |  |
| S<sub>GLY</sub> | 11.3 | 15-50% ACN, 30 min, 220 nm | 786.83 | 787.49 | 21<sup>2a</sup>; 39<sup>2b</sup> |
| M<sub>GLY</sub> | 9.8 | 50-100% ACN, 30 min, 220 nm | 955.15 | 955.61 | 24<sup>2a</sup>; 33<sup>2b</sup> |
| M<sub>ALA</sub> | 10.4 | 50-100% ACN, 30 min, 220 nm | 969.18 | 969.67 | 26<sup>2a</sup>; 34<sup>2b</sup> |
| S<sub>GLY</sub>-PNA<sub>anti-rfp</sub> | 13.6 | 10-50% ACN, 30 min, 267 nm | 4830.84 | 4830.60 | 34 |
| S<sub>GLY</sub>-PNA<sub>SCR</sub> | 14.3 | 10-50% ACN, 30 min, 267 nm | 4830.84 | 4830.10 | 33 |
| M<sub>GLY</sub>-PNA<sub>anti-rfp</sub> | 14.9 | 10-90% ACN, 30 min, 267 nm | 4999.16 | 4998.70 | 44 |
| M<sub>GLY</sub>-PNA<sub>SCR</sub> | 14.7 | 10-90% ACN, 30 min, 267 nm | 4999.16 | 4999.00 | 23 |
| M<sub>ALA</sub>-PNA<sub>anti-rfp</sub> | 14.7 | 10-90% ACN, 30 min, 267 nm | 5013.19 | 5013.10 | 38 |
| M<sub>ALA</sub>-PNA<sub>SCR</sub> | 15.2 | 10-90% ACN, 30 min, 267 nm | 5013.19 | 5012.90 | 34 |

### Synthesis of PNA and their conjugates with siderophores

All PNA oligomers were synthesized with 10-undecynoic acid attached manually to the resin following the Fmoc strategy (Tsylents et al., 2024; Wojciechowska et al., 2014). PNA-siderophore conjugates were obtained using the CuAAC reaction as described in (Tsylents et al., 2024) with one modification: prolonged ultrasound treatment for up to 45 minutes. PNA sequences and their conjugates are shown in Figure 7. The products were lyophilized and purified (Table 3, Supplementary Figures S24–S29).

### Coprogen production and purification

Coprogen was produced using the *Neurospora crassa* wild-type strain 74A (obtained from the FGSC (McCluskey et al., 2010)). *N. crassa* was maintained on yeast malt extract with glucose (YMG) agar plates according to (Huschka et al., 1985). Mycelia were scraped from approximately 15 cm^2^ of a fully covered plate and used to inoculate 100 mL of a synthetic medium optimized for coprogen production. The medium consisted of glucose 40 g/L, aspartate 21 g/L, K_2_HPO_4_ x 3H_2_O 1 g/L, MgSO_4_ x 7H_2_O 1 g/L, CaCl_2_ x 2H_2_O 0.05 g/L, ZnSO_4_ x 7H_2_O 0.02 g/L, and biotin 25 µg/L, with the pH adjusted to 7.2. Cultivation was performed for 10 days at 28°C with stirring at 250 rpm (Tóth et al., 2009). The culture was then filtered through a 0.22 μm bottle filter, and the mycelia were thoroughly washed with mQ water. The filtrate was supplemented with 6 mM FeCl_3_, and the pH was adjusted to 6.5 (Tóth et al., 2009). The solution was aliquoted into smaller fractions and subsequently used for coprogen purification.

Coprogen was purified using previously published methods (Amsri et al., 2022; Pócsi et al., 2008; Wong et al., 1983) with minor adjustments. Briefly, aliquots of post-culture liquid were loaded onto an Amberlite XAD-2 column (approx. 80 mL of resin per 1 L of liquid) equilibrated with water. The column was washed with 3-5 column volumes (CVs) of water, and siderophores were eluted with 1 CV of 50% (v/v) acetone. The acetone was evaporated. The sample was concentrated before further purification *via* size exclusion chromatography (SEC) to remove remaining impurities, including other siderophores, such as the hydroxamate dimerum acid (a precursor or a degradation product of coprogen). SEC was performed on a Bio-Gel P-2 column (CV ≈ 78 mL) equilibrated with 5 mM Tris-HCl buffer (pH 7.0) containing 15 mM NaCl. The same buffer was used for elution. Purity was assessed using RP-HPLC and MS. Fractions with the highest purity were used for experiments with ferric coprogen.

Desferric coprogen fractions were obtained using a protocol similar to iron removal from legiobactin (Allard et al., 2009). Siderophore solution was mixed with an 8% (w/v) solution of 8-hydroxyquinoline (8-HCQ) in dichloromethane (DCM) at a 1:3 ratio. The mixture was shaken vigorously at 4°C overnight. The organic fraction was separated using a separatory funnel and discarded. This procedure was repeated once with a fresh 8-HCQ solution and three times with DCM. The final aqueous fraction was purified and analyzed using RP-HPLC (using both analytical and semi-preparative chromatography) with a gradient proposed by Genaxxon BioScience GmbH (Ulm, Germany) designed for use with the Coprogen HPLC Calibration Kit (discontinued): 4%-10% ACN, 10 min; 10%-20% ACN, 15 min; 20%-60% ACN, 25 min; 60%-100% ACN, 30 min. The detector wavelength was set to 220 nm (see Supplementary Figure S30).

### CD measurements

CD spectra of iron-free coprogen and S_GLY_ were recorded in 100 mM sodium phosphate buffer, pH 7.0, with iron(III) chloride (FeCl_3_) as previously described (Tsylents et al., 2024). M_GLY_ and M_ALA_ siderophores were dissolved in ACN/phosphate buffer (1:1, v/v) and incubated with FeCl_3_ for 30 minutes with shaking. In all cases, the siderophore/Fe^3+^ molar ratio was maintained at 1:1. The optimal incubation time with Fe^3+^ was adjusted for each siderophore (Supplementary Figure S31). Spectra were recorded on the Biokine MOS-450/AF-CD spectrometer equipped with a Xe lamp, using a 0.1 cm CD cell. Acquisition parameters included a resolution of 1 nm and a scan time of 2 s per data point. Measurements were performed at room temperature over the wavelength range of 300–500 nm. Spectra were smoothed with the Savitzky–Golay method using GraphPad, and the spectra represent the averages of three scans. Each experiment was repeated twice to confirm reproducibility.

### Competition assay with EDTA

The spectrophotometric protocol for the EDTA competition experiments was adapted from (Mies et al., 2008). Stock solutions of siderophores were incubated overnight in 100 mM sodium phosphate buffer, pH 7.4, with FeCl_3_ (siderophore/Fe^3+^ molar ratio of 1:1) at room temperature. Spectrophotometric measurements used a 96-well, flat-bottom, transparent plate with 200 µL volume per well. Calibration curves used serial dilutions of ferric-siderophore complexes, with concentrations from 0.222 mM to 0.00222 mM (Supplementary Figure S32). Competition assays used a fixed ferric-siderophore concentration (0.222 mM) with varying EDTA amounts (stock concentration: 37 mM) added to the buffer solution. After incubation for 2-3 days at room temperature, absorbance was recorded using a Perkin Elmer 2300 Multilabel Reader with Enspire software. Experiments were repeated with three independent ferric-siderophore stocks, and duplicate measurements were taken for each condition. Statistical significance was calculated using a one-way ANOVA.

### Growth recovery assay

The growth recovery experiments followed previous protocols (Tsylents et al., 2024; Zheng et al., 2012). *E. coli* mutants were grown in LB medium overnight and then diluted 1:100 in fresh 50% MHB medium (2 mL) with or without 200 μM DP. Cultures were incubated at 37°C with shaking until reaching an OD_600_ of 0.2, then diluted to an OD_600_ of 0.001. These diluted cultures were aliquoted (90 μL per well) into a 96-well flat-bottom, fully-transparent plate. Siderophores were dissolved in water (S_GLY_) or DMSO (M_GLY_ and M_ALA_), preloaded (overnight incubation) with iron (1:1 siderophore: FeCl_3_ ratio), and added to each well at 16 μM (10 μL per well). DMSO was kept at 3% (v/v) where applicable. Growth Control (GC) wells included additional iron introduced through siderophore preloading, and GCs containing DMSO if needed. Plates were sealed with adhesive foil and incubated at 37°C with shaking for 19 h. OD_600_ was measured hourly using the Biotek Microplate Reader (Supplementary Figure S33). Each condition had duplicate measurements, with three independent biological replicates conducted on separate days. Sterility Control (SC) OD_600_ values were subtracted from all readings, and measurements were normalized to the appropriate GC.

### Fluorescence silencing assay

2 mL of MHB medium was inoculated with a freezer stock of *E. coli* JW0669 Δ*fur* strain (from the Keio Collection, Yale University, New Haven, CT, USA) carrying the pBBR1MCS5(*rfp*) plasmid, and cultured overnight at 37°C with shaking (600 rpm). The culture was then diluted 1:100 in 2 mL of fresh 50% MHB medium with 200 μM DP. Bacteria were cultured at 37°C with shaking until OD_600_ = 0.5, then transferred to a sterile 2 mL Eppendorf-type tube, centrifuged (10000 rpm, 10 minutes) at 4°C, and resuspended in 1 mL of fresh 50% MHB medium with 200 μM DP. 90 µL of this suspension was transferred into black U-bottom 96-well plates (BRAND GmbH, Wertheim, Germany), and mixed with 10 µL of 160 µM compound solutions: M_GLY_, M_ALA_, S_GLY,_ M_GLY_–PNA_SCR_, M_ALA_–PNA_SCR_, S_GLY_–PNA_SCR,_ M_GLY_–PNA_anti-*rfp*,_ M_ALA_–PNA_anti-*rfp*_, S_GLY_–PNA_anti-*rfp*_, PNA_SCR_, and PNA_anti-*rfp*_. Controls included *E. coli* JW0669 Δ*fur* pBBR1MCS5(rfp) in 50% MHB without DP as growth control, and *E. coli* JW0669 Δ*fur* as a negative control. After 20 h incubation at 37°C with shaking (200 rpm), RFP fluorescence (excitation 555 nm, emission 583 nm) and bacterial OD_600_ were measured using a BIOTEK Synergy H1MFDG Microplate Reader. To reduce excessive fluorescence from the siderophore’s growth promotion effect, the fluorescence intensity, measured in each well, was normalized by dividing by the OD_600_ value in the adequate wells. The experiment was performed in triplicate, on different days. Results are presented as relative fluorescence intensity, normalized to adequate OD_600_ values. Statistical significance was determined by a one-way ANOVA.

### Confocal microscopy imaging

Confocal microscopy imaging was used to monitor fluorescence silencing in *E. coli* by siderophore-PNA conjugates. *E. coli* JW0669 Δ*fur* pBBR1MCS5(*rfp*) samples were prepared according to the protocol described in the *Fluorescence silencing assay*, with minor modifications. Namely, bacteria were mixed with tested compounds in 1.5 mL Eppendorf-type tubes (90 µL of bacterial suspension + 10 µL of 160 µM compound solution), then incubated overnight with shaking (37°C, 600 rpm). After incubation, bacteria were centrifuged, suspended in 100 µL of fresh PBS buffer, and each sample was supplemented with 0.25 µL of SYTO 9 green fluorescent nucleic acid stain (Thermo Fisher Scientific, Waltham, MA, USA). After 30 min of incubation at room temperature, in darkness, 8 µL of each suspension was transferred onto a glass microscopic slide and covered with a cover slip. For imaging, an Axio Imager Z2 LSM 700 (Carl Zeiss, Oberkochen, Germany) was used. Samples were excited with 488 nm (SYTO 9 fluorophore) and 555 nm (RFP fluorescence) wavelengths. To compare fluorescence intensities, images were processed using ImageJ software, and the percentage of the mean integrated density was determined.

### Molecular modeling

*In silico* methods were employed to obtain structures of iron(III)-bound siderophores used for their docking to receptor proteins of the *E. coli* TBDT pathways. Initially, the ’nonbonded model’ employing the Lennard Jones 12-6-4 potential was used in MD simulations to model dative bonds between the iron(III) ion and siderophore oxygen atoms, with Cij parameters optimized *via* thermodynamic integration (TI). Subsequently, simulations using a harmonic potential, as in the bonded terms of classical force fields, with covalent bonds between iron(III) and relevant oxygen atoms, were carried out (’bonded model’). For simulations, siderophores lacked the azide-containing moieties, which were present in the synthesis to enable their conjugation with PNA using “click” chemistry. Simulations of iron(III) association using the LJ 12-6-4 nonbonded potential that were performed for M_GLY_ and M_ALA_ did not include their N-terminal aliphatic tails, but the full M_ALA_ and M_GLY_ structures (with tails) were included in all other simulations and in docking.

### Molecular dynamics simulations protocol

The simulation system and force field were prepared using AmberTools23 (Case et al., 2023) and AmberTools24 (Case et al., 2024). MD simulations were performed in Amber20 (Case et al., 2020). Ferrichrome was used as a control in the parameterization of *N*^δ^–acetyl–*N*^δ^–hydroxy–L–ornithine. Force field parameters were tested to reproduce the geometry of the coordination sphere of iron(III) in ferrichrome and then were transferred to the corresponding residues in S_L_, S_GLY_, M_ALA_, and M_GLY_ molecules, since they also have a Λ configuration according to CD measurements. *N*^δ^–acetyl–*N*^δ^–hydroxy–L–ornithine (AHO) was built manually using YASARA Structure (Ozvoldik et al., 2023). Hydroxamate groups were built in the *cis* conformation with OZ oxygens deprotonated as necessary for iron(III) coordination (Figure 12). Siderophores were built using tleap.

**Figure 12.**
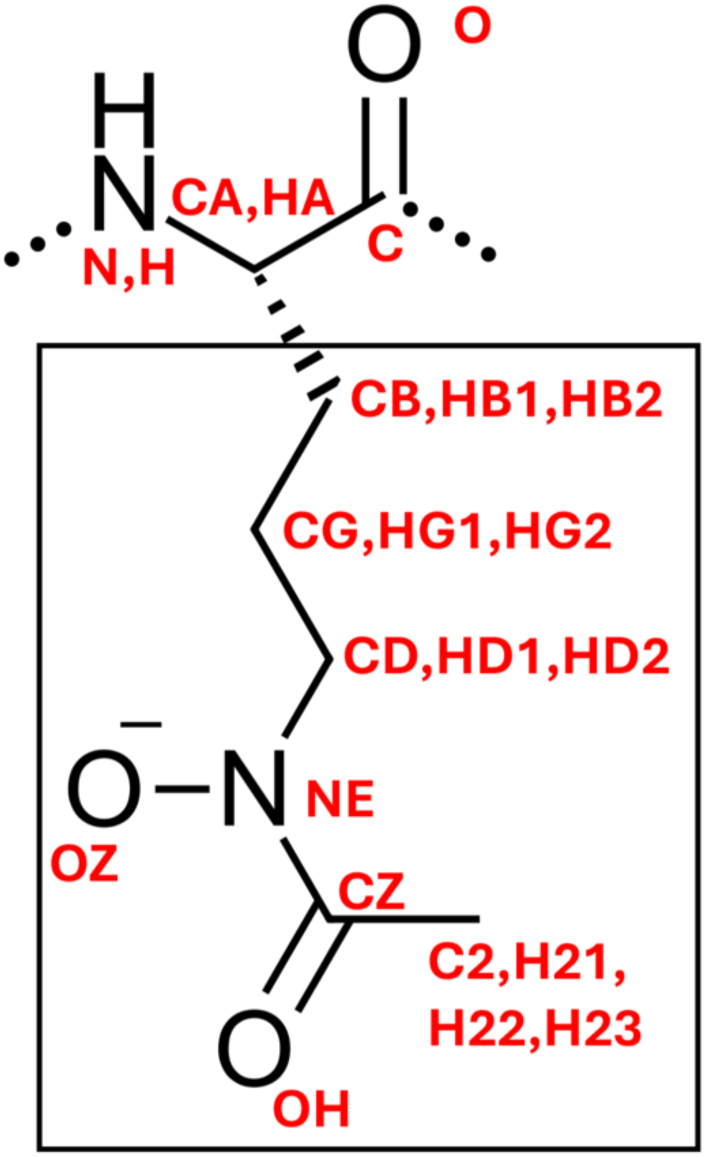
Atom names assigned to the *N*^δ^–acetyl–*N*^δ^–hydroxy–L–ornithine (AHO) residue. The frame denotes the AHO side chain used for force field parameterization, which was capped with the methyl group (the CB carbon with HB1, HB2, and HB3 hydrogens).

Siderophores were solvated in a 15 Å thick layer of SPC/E water molecules (Berendsen et al., 1987), and a random water molecule was replaced with an iron(III) ion. For ferrichrome, a crystal structure (van der Helm et al., 1980) from the CCDC database (structure ID: 1154895) with its iron ion removed was used as the initial structure. Periodic boundary conditions were used in all simulations. The systems with deferrated siderophores were neutralized with 3 Na^+^ ions.

Minimization was performed with sander.MPI and all other steps with pmemd.cuda (Götz et al., 2012; Le Grand et al., 2013; Salomon-Ferrer et al., 2012). The MD simulations protocol is detailed in Supplementary Material Section S1. An integration time step of 2 fs was used with SHAKE for bonds involving hydrogen atoms (Miyamoto & Kollman, 1992; Ryckaert et al., 1977). Particle Mesh Ewald (PME) summation was used for long-range electrostatics, and a cutoff of 9 Å for nonbonded interactions (Cheatham et al., 1995). A Langevin thermostat was used with a collision frequency of 1 ps^-1^. Pressure at 1 atm was controlled with the Monte Carlo barostat. Trajectories were analyzed using CPPTRAJ (Roe & Cheatham, 2013). Production runs (performed in triplicate with different initial velocities) in the nonbonded model were 200 ns. For siderophores bound covalently to iron(III), the production stage was 100 ns.

### Thermodynamic integration

Thermodynamic integration (TI) was carried out in pmemd.cuda to calculate the free energy of binding of iron(III) to ferrichrome dependent on a set of *C_ij_* parameters of the nonbonded LJ 12-6-4 potential form. One system was composed of an iron(III) ion solvated in a 16 Å layer of SPC/E water. The iron(III) was transformed into a dummy atom not interacting with the system, using a one-step transformation, in which electrostatic and van der Waals interactions were eliminated simultaneously. The free energy of removing the ion from the system was calculated, and was taken with the opposite sign to obtain the free energy of adding iron to the system. Values calculated for iron(III) alone were subtracted from those determined for the system with ferrichrome, so that the difference corresponded to the free energy of binding of the ion initially solvated in water. The target value of the Gibbs free energy Δ*G* was calculated using the association constant *K_a_* derived from the literature (Anderegg et al., 1963) and was equal to -38.97 kcal/mol at 20°C. Details of the TI protocol and calculations are provided in Supplementary Material Section S2 with Table S3. Three series of TI calculations were performed for the iron(III) hydration free energy and the calculated value was averaged to -1020 kcal/mol. This value was then subtracted from the calculated values for the iron(III) ion in ferrichrome. The average TI-determined free energy of iron(III) binding to ferrichrome for the chosen set of *C_ij_* parameters was equal to -41.35 kcal/mol.

### Force field preparation

Atom names for *N*^δ^–acetyl–*N*^δ^–hydroxy–L–ornithine (AHO) are shown in Figure 12. Atom types and charges are listed in Supplementary Table S2. *C_ij_* values for atoms outside the hydroxamate groups were set to zero. Nonbonded and covalent bond parameters were taken from the ff19SB (Tian et al., 2020) AMBER force field. Missing bonded parameters were taken from the GAFF2 parameters (He et al., 2020). Charge fitting was performed using the RESP method (Cornell et al., 1993). Quantum calculations were performed using Gaussian16 (Frisch et al., 2016). First, calculations were performed for the AHO sidechain (Figure 12), which was optimized at the B3LYP/6–311+G** level of theory. Merz-Kollman population analysis was then performed at the HF/6–311+G** level of theory, generating data that were processed in espgen and used for charge fitting in resp. Subsequently, population analysis at the HF/6–311+G** level of theory was performed for the full *N*^δ^–acetyl–*N*^δ^–hydroxy–L–ornithine (AHO), terminated with an acetyl group at the N-terminus and a methylamide group at the C-terminus. Charges for atoms CD, HD1, HD2, NE, OZ, CZ, OH, C2, H21, H22, and H23 were set to the same values as calculated for the AHO sidechain. Backbone atom charges, except for CA and HA, were assigned values from the ff19SB force field. This procedure resulted in a set of *C_ij_* parameters (Supplementary Table S4) that were tested in MD simulations of ferrichrome and that yielded the binding free energy and molecule geometry resembling experimental values. These parameters were then also applied in MD simulations of iron(III) binding by S_GLY_, M_ALA_, and M_GLY_.

Parameterization of covalent bonds between the iron(III) ion and oxygen atoms of hydroxamate groups in the bonded model was carried out using MCPB.py (Li & Merz, 2016), Gaussian16 (Frisch et al., 2016), and the Seminario method (Seminario, 1996) for the AHO sidechain capped with a methyl group (the framed structure in Figure 12 and Supplementary Figure S34). The ferrichrome crystal structure (CCDC ID: 1154895; van der Helm et al., 1980) was used with the backbone removed, leaving only the AHO sidechain (Figure 12). The multiplicity was corrected to 6. The structure was optimized, and force constant calculations were performed to ensure that local minima were found. The B3LYP/6-31G* level of theory was used throughout. A fragment of the ferrichrome structure was also used for charge refitting. The ferrichrome backbone was present in these calculations, terminated with an acetyl group at the N-terminus and an amide group at the C-terminus. Charges were fitted using RESP. Backbone atoms were assigned the same charge as in the LJ 12-6-4 nonbonded model. All other atoms were subject to charge fitting, with atoms of the same type enforced to have identical charges when fitted, to obtain a universal force field for *N*^δ^–acetyl–*N*^δ^–hydroxy–L–ornithine (AHO). Atom names, types, charges, and the corresponding .frcmod file are provided in Supplementary Section S3.

The N-terminal myristic acid of M_ALA_ and M_GLY_ was assigned the lipid 21 AMBER force field parameters (Dickson et al., 2022), with refitted charges from .prepgen, and carbonyl group atom types from ff19SB, with missing parameters taken from GAFF2 (Supplementary Table S5). M_ALA_ and M_GLY_ used in the nonbonded model simulations did not contain the N-terminal myristic acid and were N-terminated with acyl groups (Figure 10).

### Clustering analysis

Starting conformations for MD simulations using the bonded model of iron(III) were taken from MD simulations of iron(III) association with siderophores using the LJ 12-6-4 nonbonded model. Conformations were selected where all three hydroxamate groups coordinated the iron(III) ion. These frames were chosen *via* k-means clustering (Shao et al., 2007) after water removal. The 50–200 ns time interval was used for S_L_ and S_GLY_, and the 100–200 ns for M_GLY_ and both M_ALA_ trajectories. The number of clusters was determined based on the Davies-Bouldin Index (DBI) and pseudo-F statistics (pSF) metrics. For each siderophore, a representative structure with a coordination sphere geometry closest to the Λ configuration was selected. Siderophore conformations used for docking were selected from clustering of the last 50 ns of a 100 ns MD trajectory simulated with the bonded model. Clustering metrics are given in Supplementary Figures S35 and S36.

### Molecular docking

Molecular docking was performed as described in (Tsylents et al., 2024), using AutoDock Vina (Eberhardt et al., 2021) with an updated 25.1.13 version of YASARA Structure (Ozvoldik et al., 2023). The number of runs in the dock_run macro was set to 100 and the number of local dockings per ligand in the dock_rescore macro was set to 25. During docking to FhuF, atoms forming the [2Fe-2S] cluster were allowed to move during the minimization of each ligand pose. Structures used as receptors were taken from PDB with IDs: FhuE, 6e4v (Grinter & Lithgow, 2019a); FhuA, 1by5 (Locher et al., 1998); FhuD, 1esz (Clarke et al., 2002); and FhuF, 7qp5 (Trindade et al., 2023). Crystal water molecules were deleted, and the residues were protonated as described in (Tsylents et al., 2024). The box length was set to either 30 Å (for FhuE and FhuA) or 25 Å (for FhuD and FhuF), and the box was centered on the crystal ligand. For the FhuF *apo*-structure, blind docking of ferrichrome was performed as a control with the docking box extended another 10 Å around FhuF atoms. Since the top-scoring ferrichrome pose was consistent with the binding mode proposed in the literature (Trindade et al., 2023), further docking calculations were performed with a box centered on the coordinates of ferrichrome obtained *via* blind docking. Coprogen and ferrichrome were extracted from the crystal structures and other ligands were obtained from the MD trajectories. Crystallographic ligands were protonated, and dative metal coordination bonds were added as in (Tsylents et al., 2024). Visualisation of the interactions in the docked complexes was based on default parameters in PoseEdit (Diedrich et al., 2023).

## Author Contributions

JT conceived and supervised the project. JT, MWo and UT designed modified siderophores. AM synthesized modified ornithine derivative. UT, PM, MWo synthesized S_GLY_. UT, PM synthesized M_GLY_ and M_ALA_ siderophores, UT performed CD, spectrophotometric competition and growth recovery experiments. UT synthesized PNA-siderophore conjugates. MWd performed fluorescence silencing experiments. PM obtained and purified coprogen, and performed molecular docking studies. JS performed force field parameterization, molecular dynamics simulations and trajectory analyses. UT, PM, and JT wrote the first manuscript draft. JT revised the manuscript. All authors discussed the results.

## Acknowledgements

The authors acknowledge support from the National Science Centre (OPUS 19, UMO-2020/37/B/NZ1/02904). We thank Jacek Olędzki from the Laboratory of Mass Spectrometry of the Institute of Biochemistry and Biophysics of the Polish Academy of Science for recording MS spectra. *N. crassa* 74A strain, originally obtained from The Fungal Genetics Stock Center (FGSC), was kindly donated by Prof. Joanna Kruszewska and Dr Sebastian Piłsyk from the Laboratory of Fungal Biology at the Institute of Biochemistry and Biophysics of the Polish Academy of Science.

## Additional Information

Supplementary Material is available as a separate file

## Data availability statement

Data obtained or analyzed in this study are either in the Supplementary Material or can be provided upon reasonable request.

