## Supporting Information File for "Modulating backbone flexibility in hydroxamate siderophores for improved iron chelation and peptide nucleic acid (PNA) delivery into bacteria"

**Table S1.** Summary of the CD experiments for iron(III)-siderophore complexes.

| Siderophore | Wavelength [nm] | | Optimal $Fe^{3+}$ complex configuration found | Ref. |
|---|---|---|---|---|
| | Min. | Max. | | |
| **Coprogen** | 469 | 376 | Δ | (Wong et al., 1983) |
| **$Fe^{3+}$-$S_L$** | 360 | 458 | Λ | (Tsylents et al., 2024) |
| **$Fe^{3+}$-$S_{GLY}$** | 358 | 466 | Λ | This study |
| **$Fe^{3+}$-$M_{ALA}$** | 362 | 455 | Λ | |
| **$Fe^{3+}$-$M_{GLY}$** | 361 | 454 | Λ | |

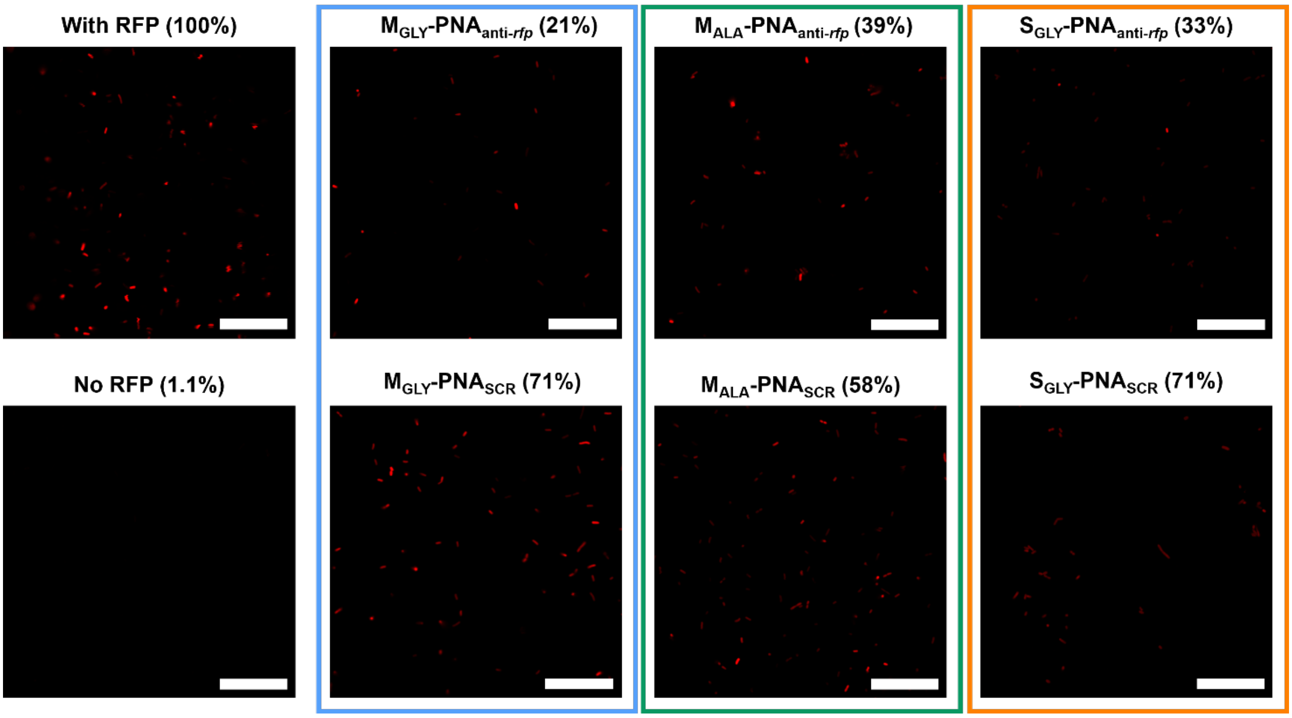


**Supplementary Figure S1.** Confocal microscopy images showing RFP fluorescence silencing of siderophore-PNA conjugates, at a concentration of 16 µM, in the *E. coli* Δ*fur* mutant. RFP fluorescence was excited at 555 nm wavelength. Fluorescence intensity was quantified as the mean integrated density from the entire image and is expressed as a percentage relative to untreated, RFP-expressing cells (set to 100% intensity). The scale bar is 20 µm.

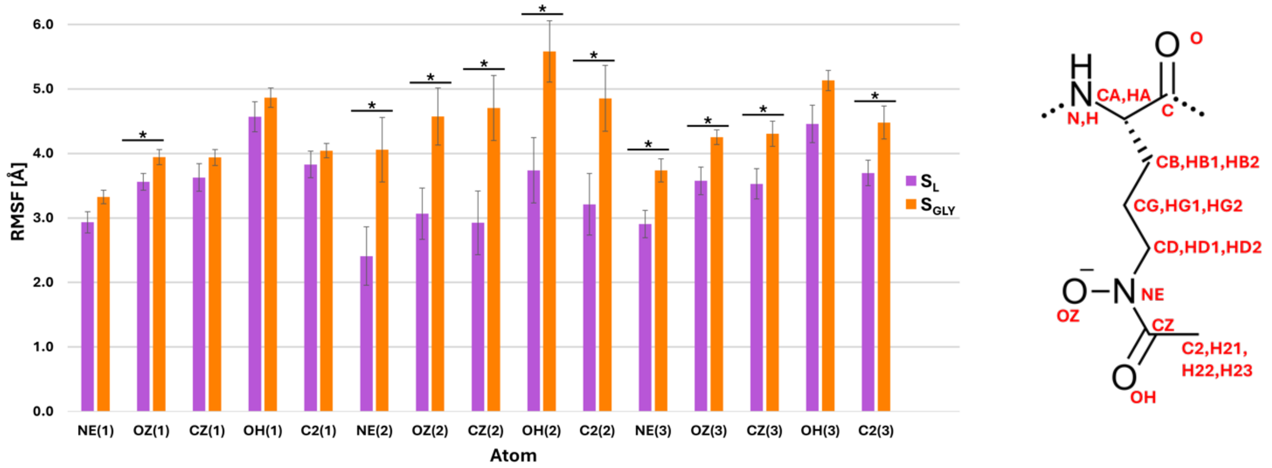


**Supplementary Figure S2.** RMSF of $S_L$ and $S_{GLY}$ heavy atoms from hydroxamate groups derived from three 200 ns trajectories in the absence of iron(III). Mean values ± SEM are shown. Statistical significance was calculated using a one-tailed t-test assuming two-sample equal variance: * $p < 0.05$. Atom names for the AHO residue are shown on the right. Numbers in parentheses number the AHO residues from the siderophore's N-terminus.

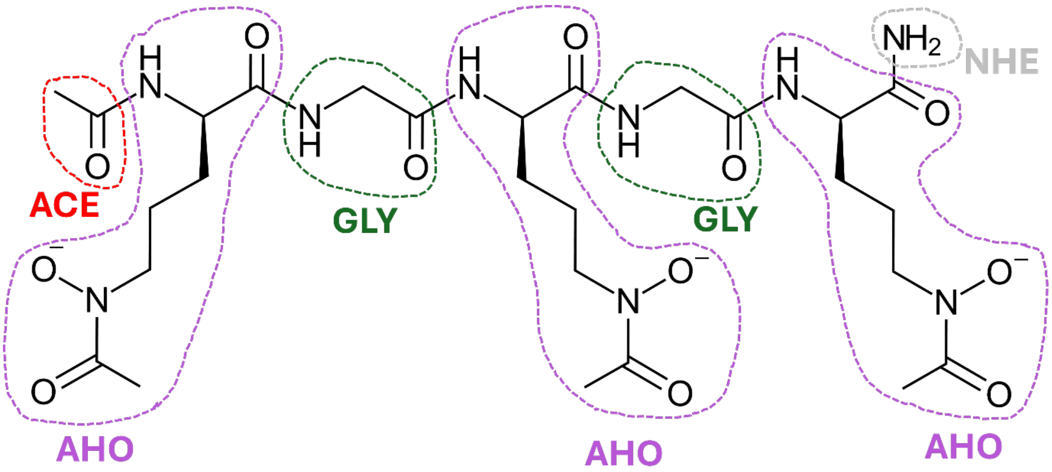


**Supplementary Figure S3.** Schematic structure of the iron-free form of the $S_{GLY}$ siderophore as used in the simulations. Residue names are consistent with those of investigated siderophores.

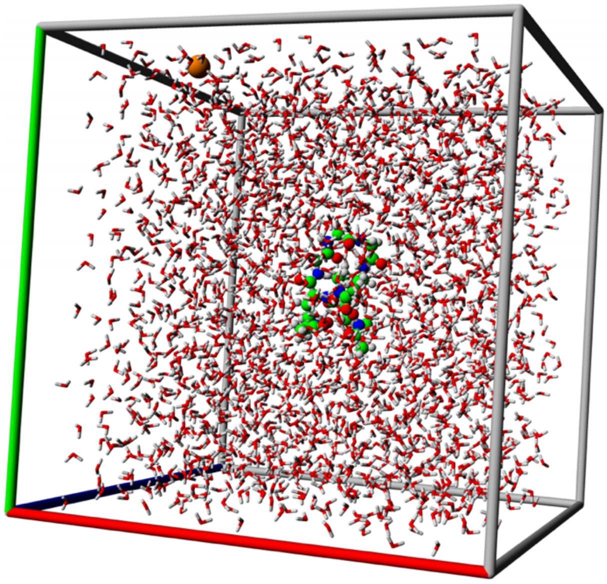

**Supplementary Figure S4.** The simulation box of 46.4 Å$^3$ showing the starting structure of desferrichrome surrounded by explicit water molecules for molecular dynamics simulations. The iron(III) cation is in orange and its initial starting position was at 16-18 Å away from the siderophore.

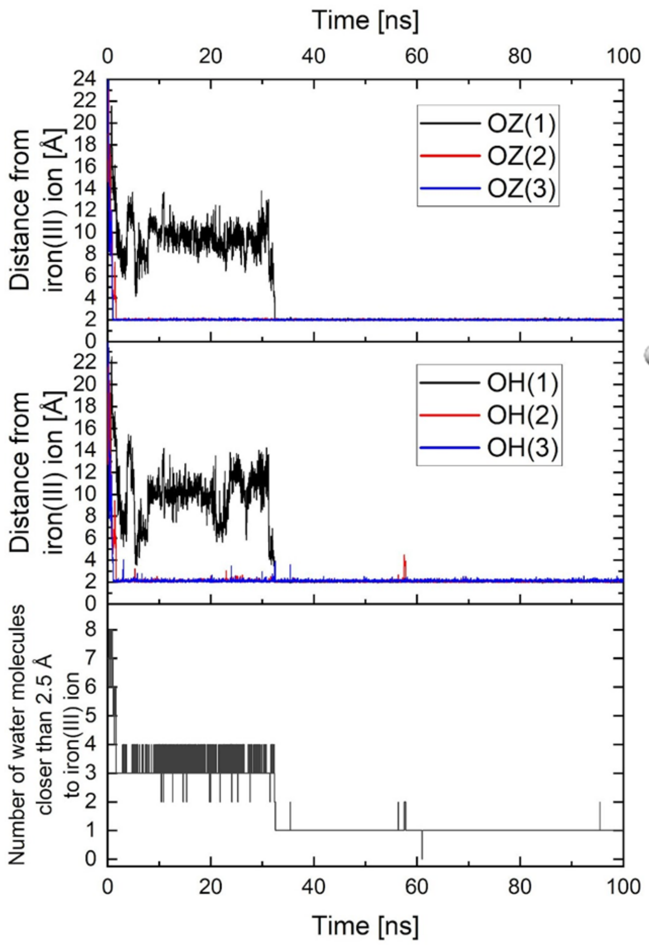


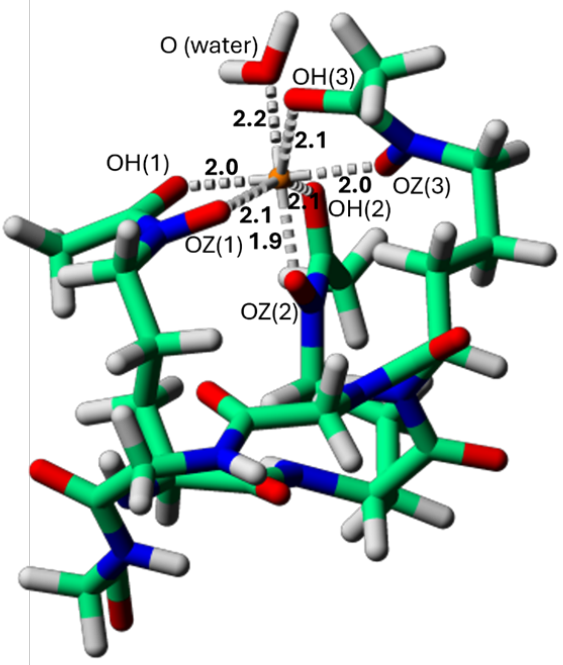


**Supplementary Figure S5.** (Left) Distances separating iron(III) and oxygen atoms from hydroxamate groups in a test simulation starting from *desferri*-ferrichrome. The number of water molecules coordinating the ion as a function of the simulation is included in the bottom graph. (Right) Structure of ferrichrome at the end of the molecular dynamics simulation (100 ns). When the parameters are assigned to correspond to the targeted value of the free energy of iron binding, at least one water molecule interacts with iron(III). $C_{ij}$ parameters were set to default values, except for OH atom types, where the parameter was set to 350 Å$^4$ × kcal/mol. Names of oxygen atoms interacting with the iron(III) ion and distances between them (in Å) are provided.

**Supplementary Table S2.** Names, atom types, and partial charges of atoms in the $N^{\delta}$–acetyl–$N^{\delta}$–hydroxy–L–ornithine (AHO) residue in the LJ 12-6-4 nonbonded model for iron-solute interactions. Parameters for analogous atom types were taken from the ff19SB force field with missing parameters taken from GAFF2, except for different $C_{ij}$ values. These atom types were introduced to modify $C_{ij}$ values for atoms forming the hydroxamate moiety independently from other atoms in the molecule.

| Atom name | Atom type | Charge [e] |
|---|---|---|
| N | N | -0.415700 |
| H | H | 0.271900 |
| CA | CX | -0.236721 |
| HA | H1 | 0.177216 |
| CB | CT | -0.024334 |
| HB1, HB2 | HC | 0.032110 |
| CG | CT | 0.120276 |
| HG1, HG2 | HC | 0.006670 |
| CD | CT | 0.009017 |
| HD1, HD2 | H1 | 0.010861 |
| NE | NH (analogous to N) | -0.013429 |
| OZ | ON (analogous to O2) | -0.748329 |
| CZ | CH (analogous to C) | 0.547939 |
| OH | OK (analogous to O) | -0.730387 |
| C2 | CT | -0.127082 |
| H21, H22, H23 | HC | 0.013650 |
| C | C | 0.597300 |
| O | O | -0.567900 |

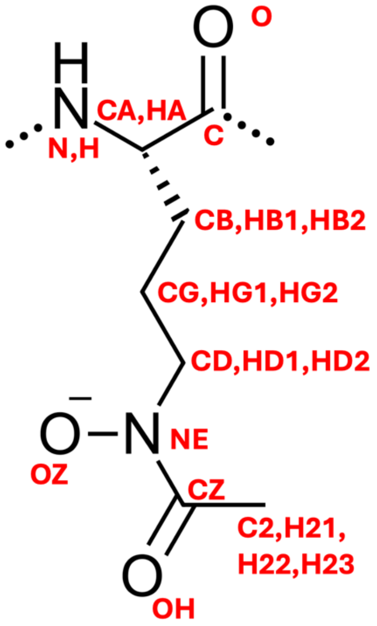


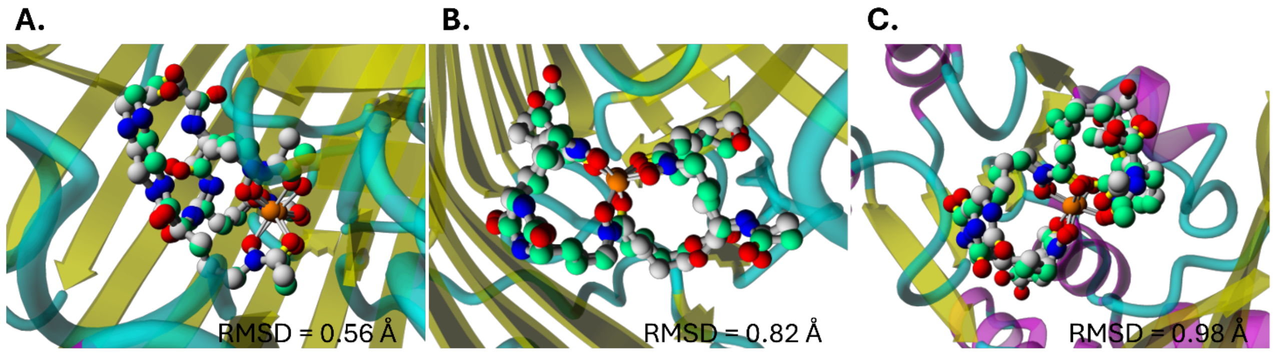


**Supplementary Figure S6.** Crystallographic (carbon atoms in grey) and re-docked (carbon atoms in green) orientations of siderophores within the binding sites of (A) FhuA (PDB ID: 1by5) and (B) FhuE (PDB ID: 6e4v), and (C) FhuD (PDB ID: 1esz) receptors. Top-scoring docking poses were superimposed onto their respective crystal structures. Root-mean-square deviations (RMSD) between the crystallographic and re-docked poses were calculated for heavy atoms. Hydrogen atoms were omitted for clarity of the superposition.

### FhuA-ferrichrome

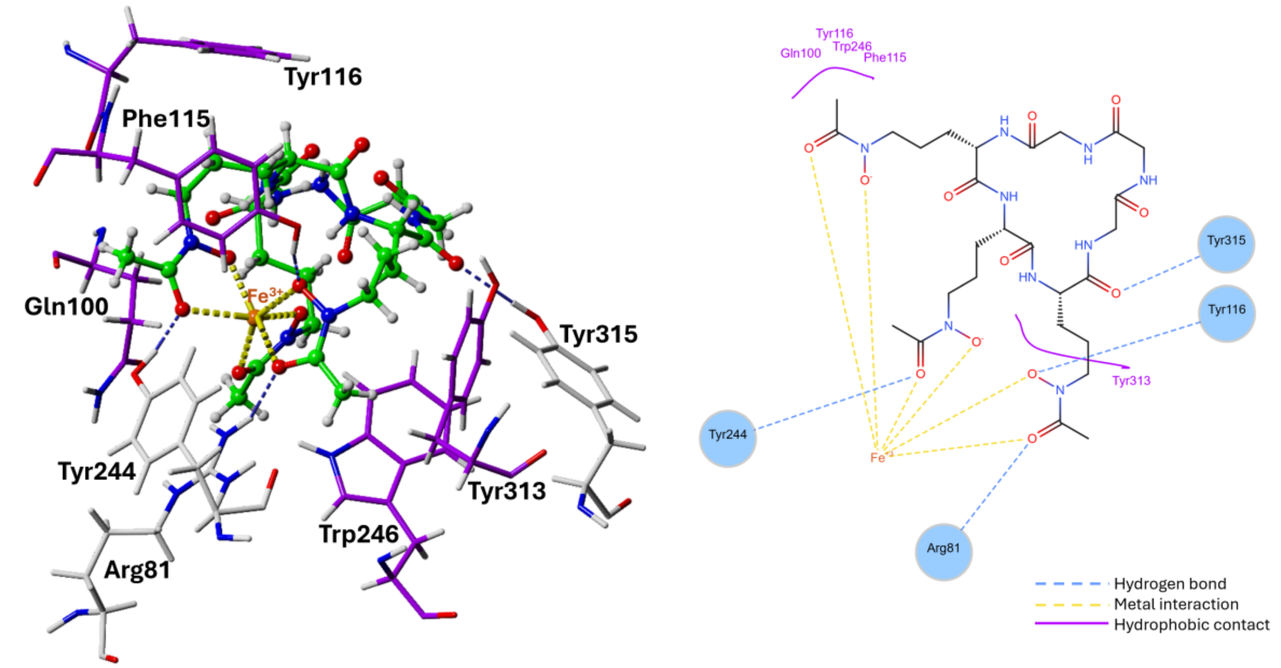


### FhuE-coprogen

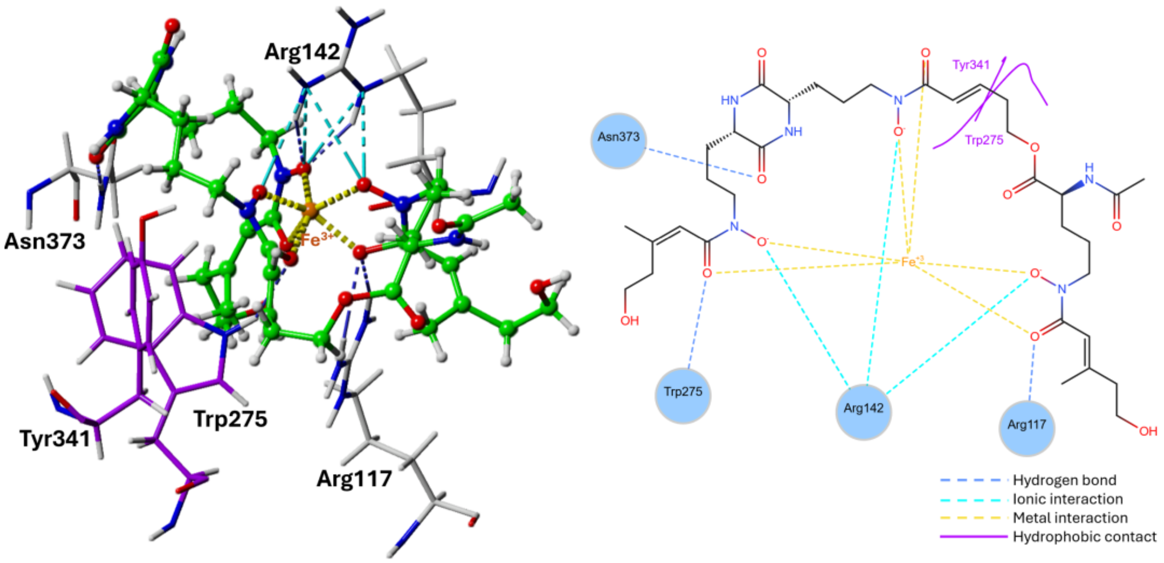


**Supplementary Figure S7.** 3D and 2D depictions of interactions between FhuA (top) or FhuE (bottom) and their respective crystallographic ligands (ferrichrome and coprogen, respectively). The interactions are based on default definitions used in PoseEdit (Diedrich et al., 2023). Amino acid residues participating in hydrophobic contacts are shown in purple.

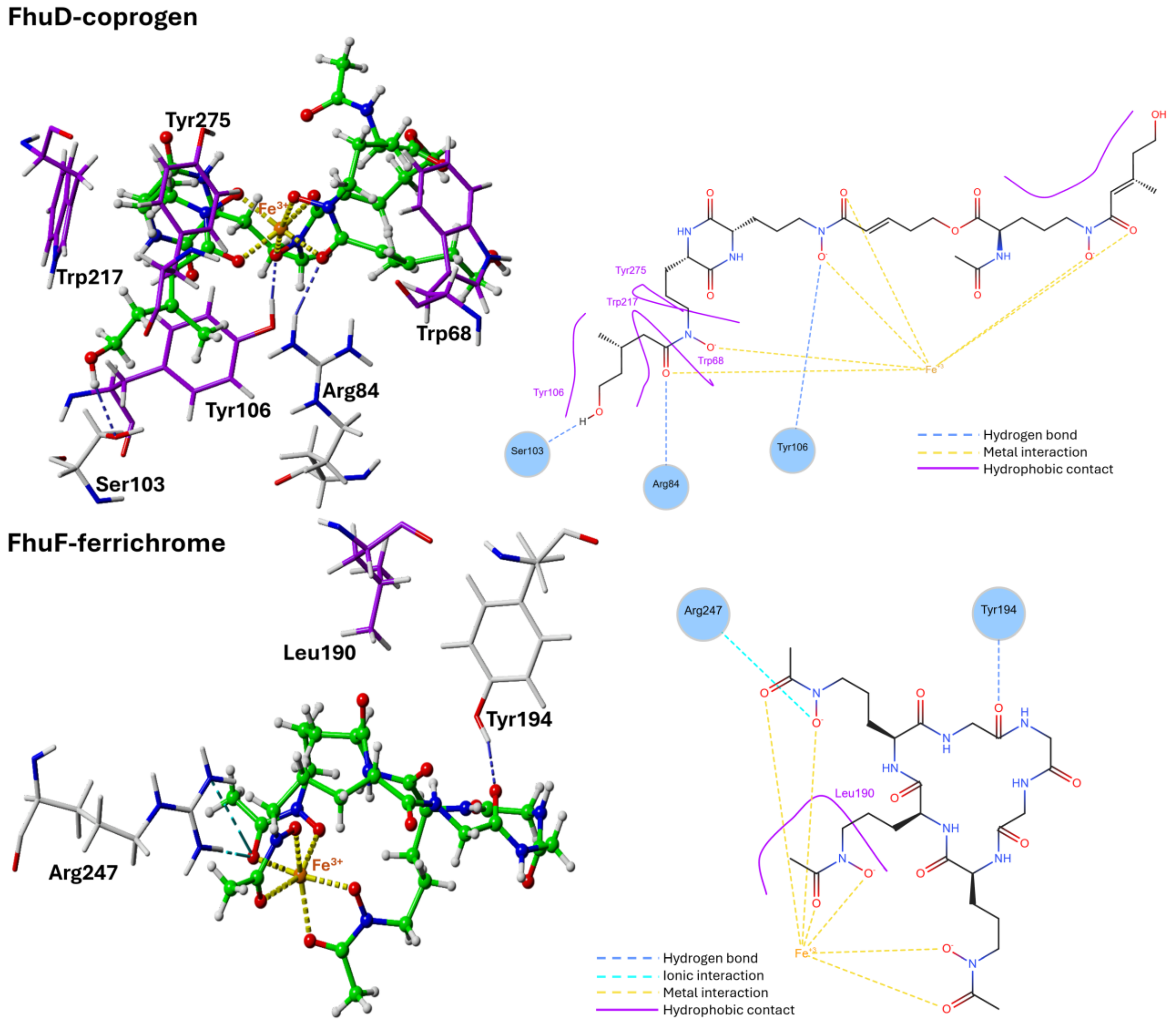


**Supplementary Figure S8.** 3D and 2D depictions of interactions between FhuD (top) or FhuF (bottom) and their ligands (crystallographic pose of coprogen and top-scoring docking pose of ferrichrome, respectively). The interactions are based on default definitions used in PoseEdit (Diedrich et al., 2023). Amino acid residues participating in hydrophobic contacts are in purple.

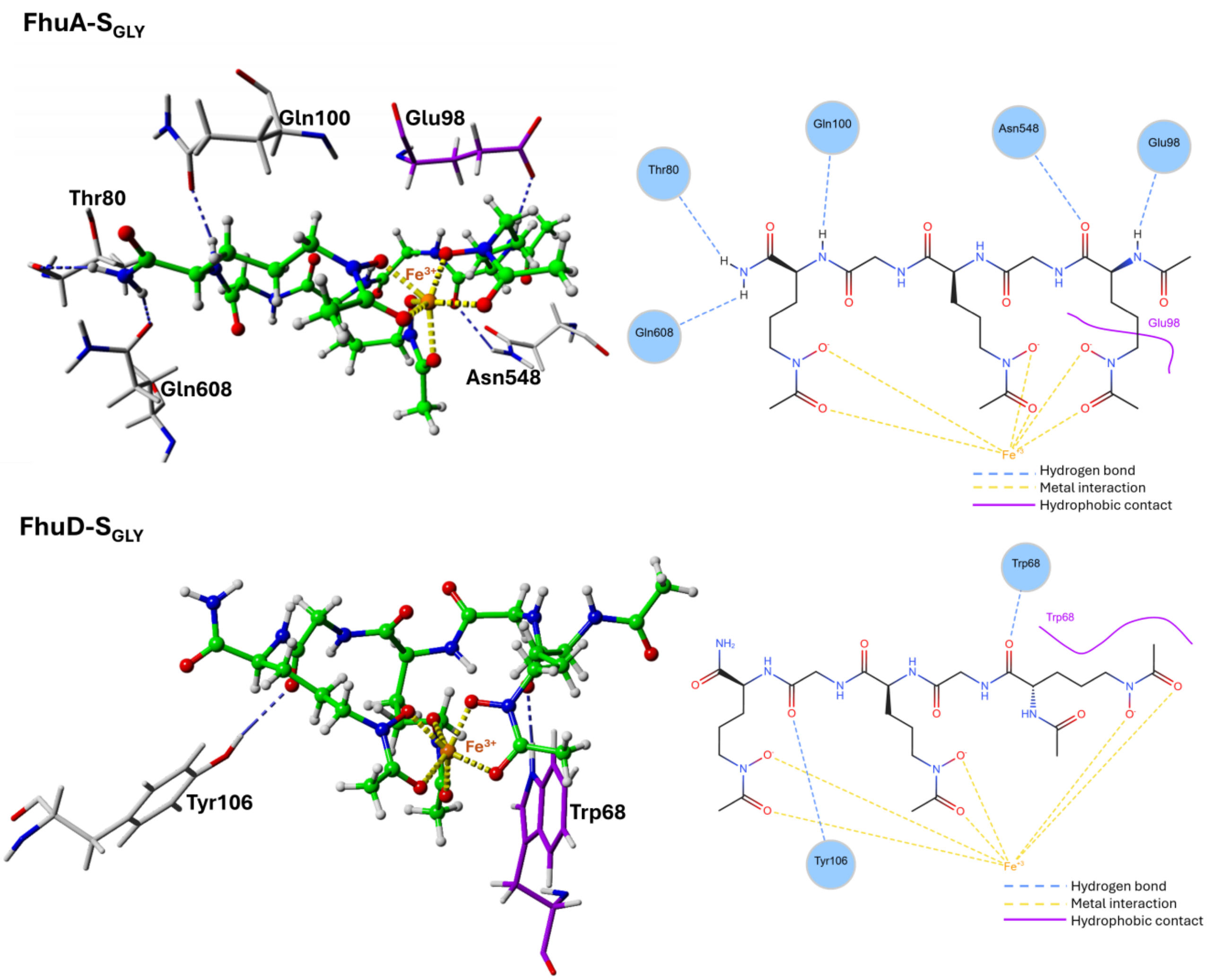


**Supplementary Figure S9.** 3D and 2D depictions of interactions between FhuA (top) or FhuD (bottom) and top-scoring docking poses of $S_{GLY}$. The interactions are based on default definitions used in PoseEdit (Diedrich et al., 2023). Amino acid residues participating in hydrophobic contacts are in purple.

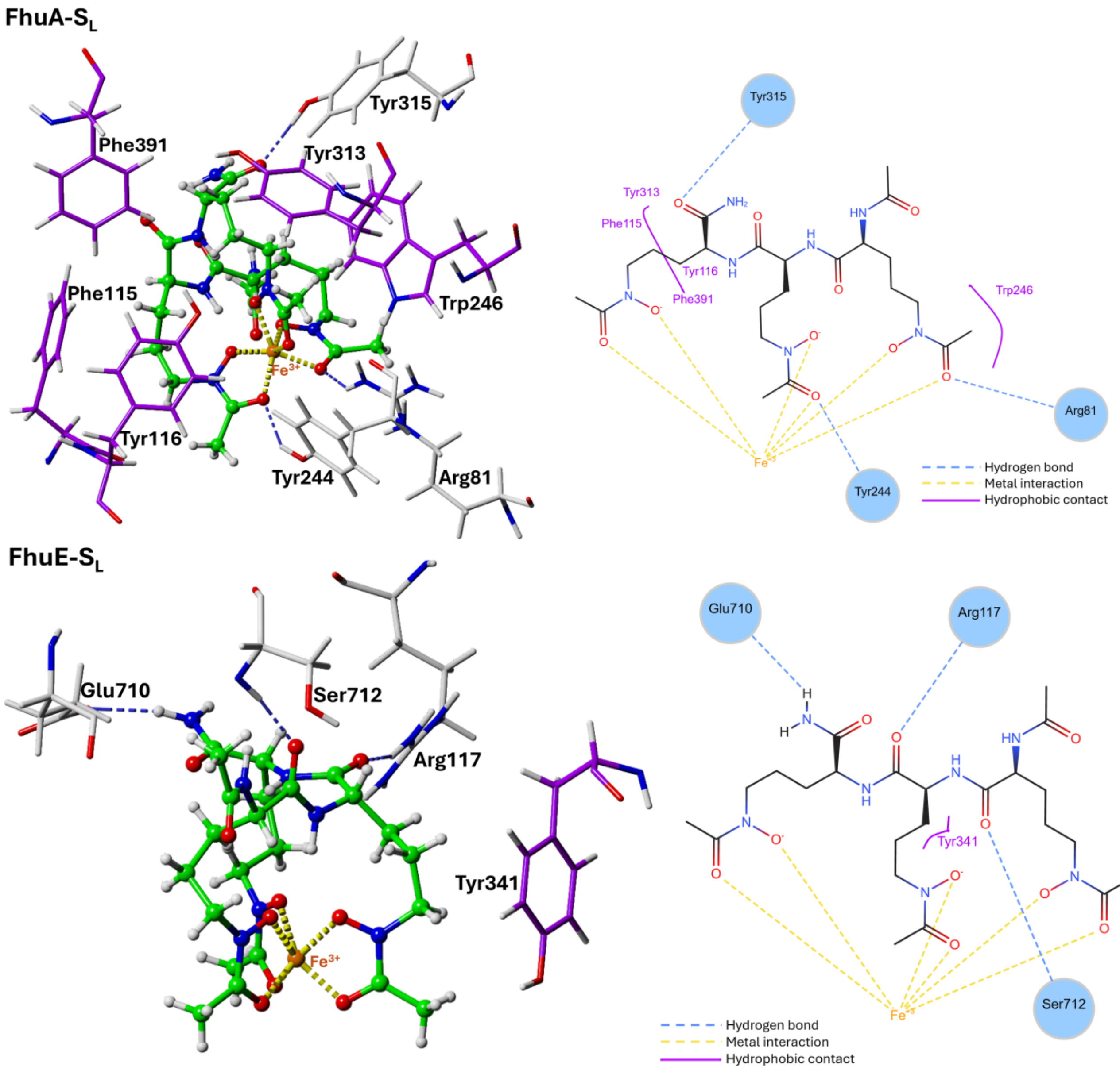


**Supplementary Figure S10.** 3D and 2D depictions of interactions between FhuA (top) or FhuE (bottom) and top-scoring docking poses of $S_L$. The interactions are based on default definitions used in PoseEdit (Diedrich et al., 2023). Amino acid residues participating in hydrophobic contacts are in purple.

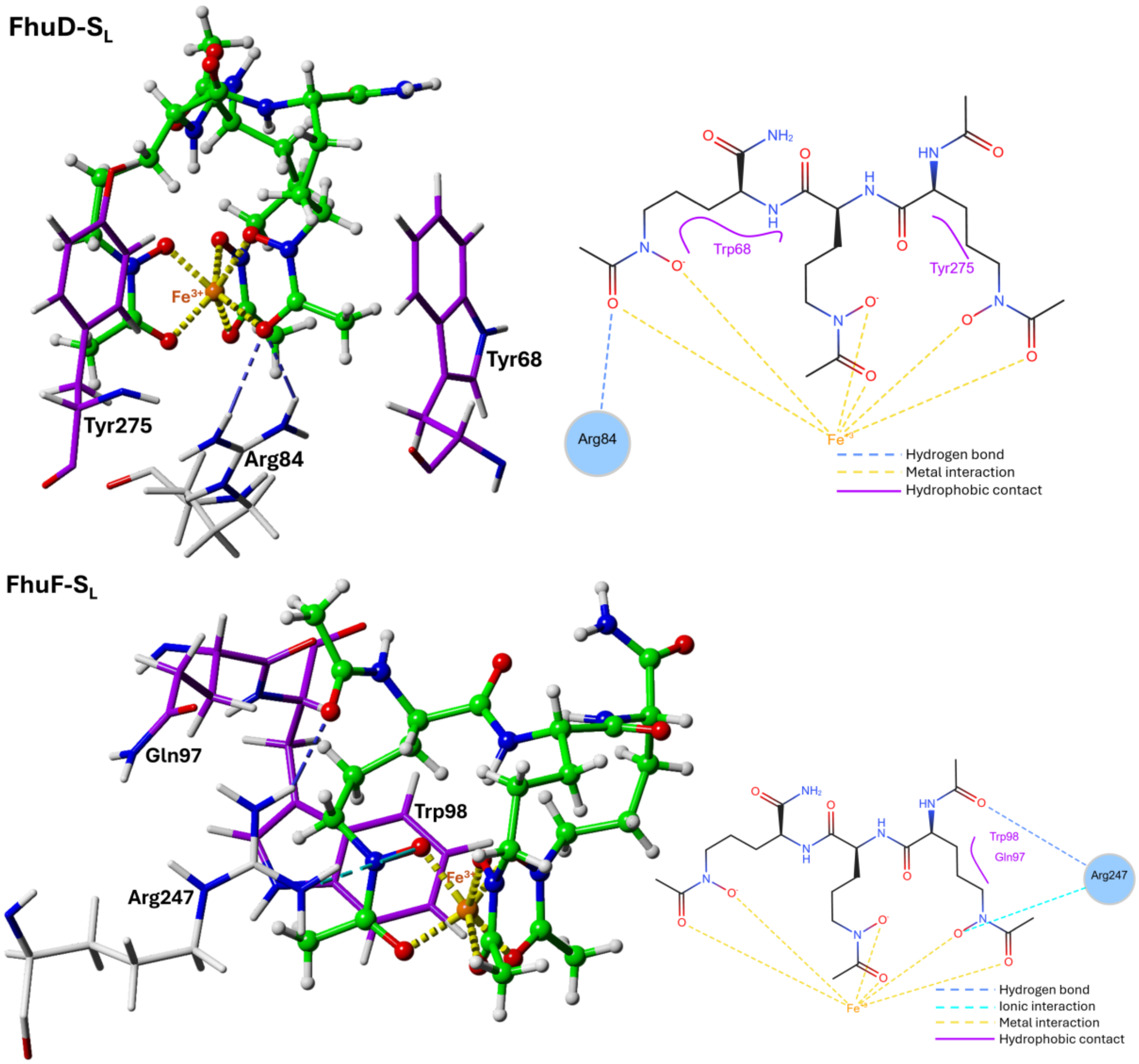


**Supplementary Figure S11.** 3D and 2D depictions of interactions between FhuD (top) or FhuF (bottom) and top-scoring docking poses of $S_L$. The interactions are based on default definitions used in PoseEdit (Diedrich et al., 2023). Amino acid residues participating in hydrophobic contacts are in purple.

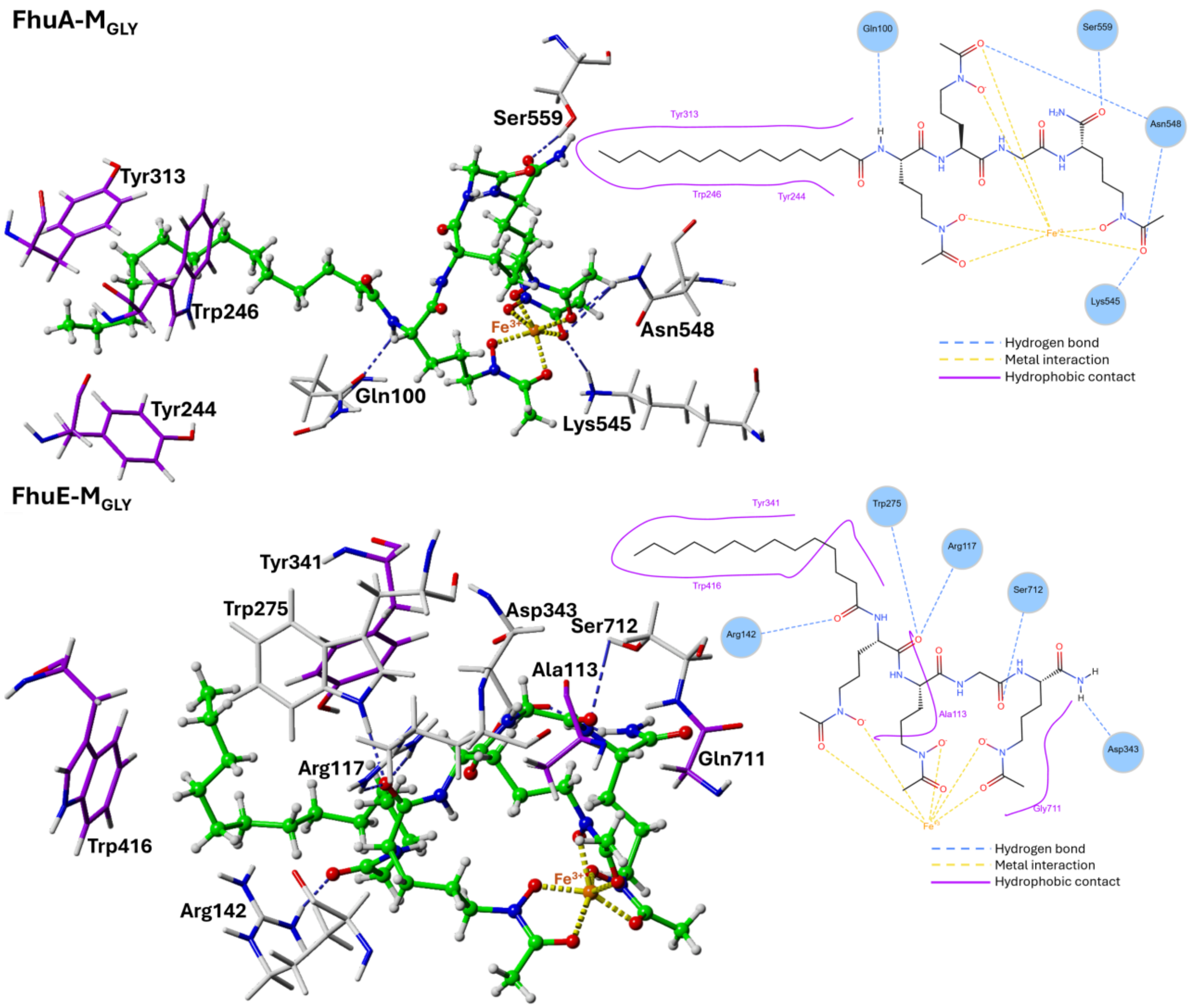


**Supplementary Figure S12.** 3D and 2D depictions of interactions between FhuA (top) or FhuE (bottom) and top-scoring docking poses of $M_{GLY}$. The interactions are based on default definitions used in PoseEdit (Diedrich et al., 2023). Amino acid residues participating in hydrophobic contacts are in purple.

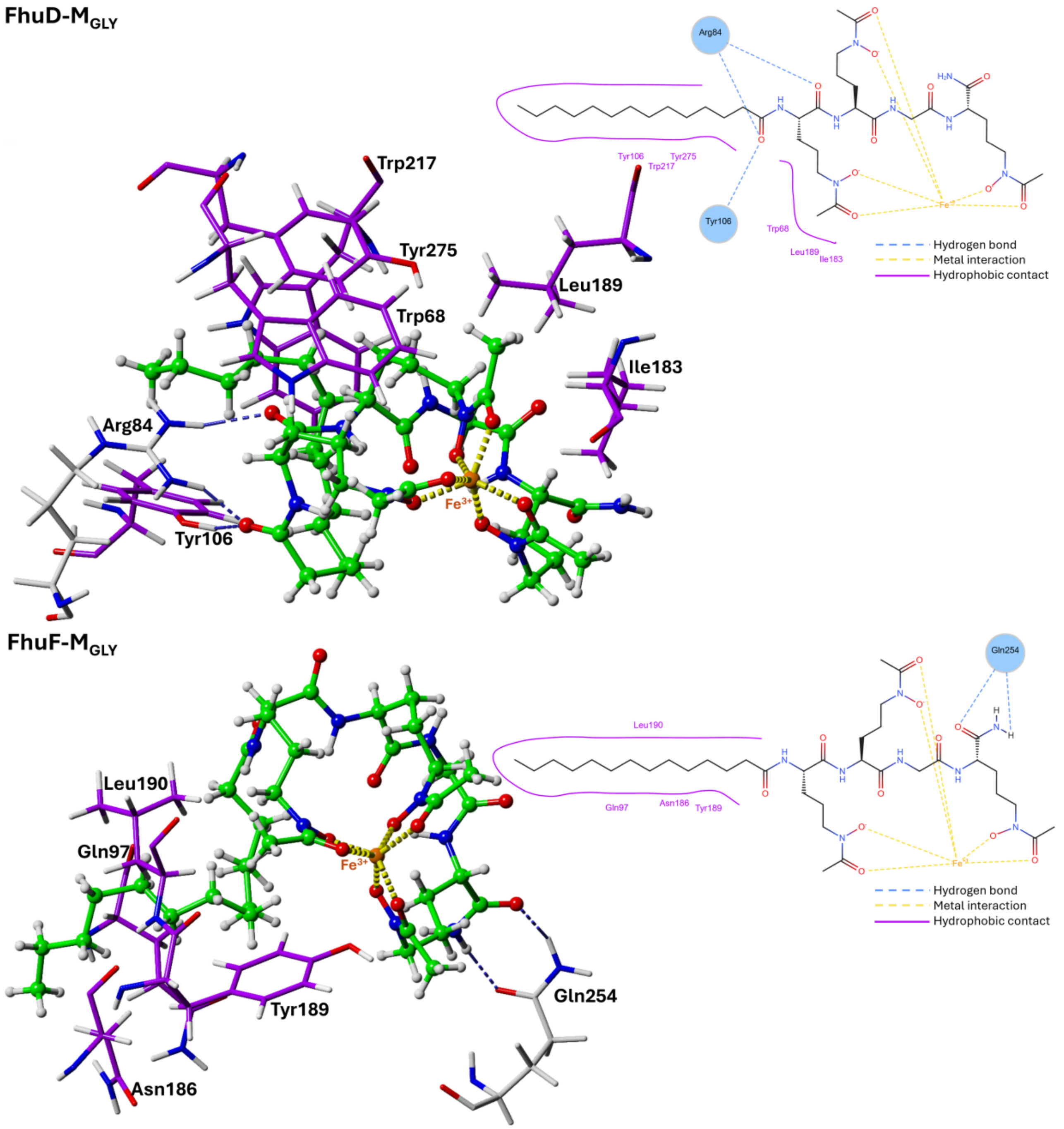


**Supplementary Figure S13.** 3D and 2D depictions of interactions between FhuD (top) or FhuF (bottom) and top-scoring docking poses of M$_{GLY}$. The interactions are based on default definitions used in PoseEdit (Diedrich et al., 2023). Amino acid residues participating in hydrophobic contacts are in purple.

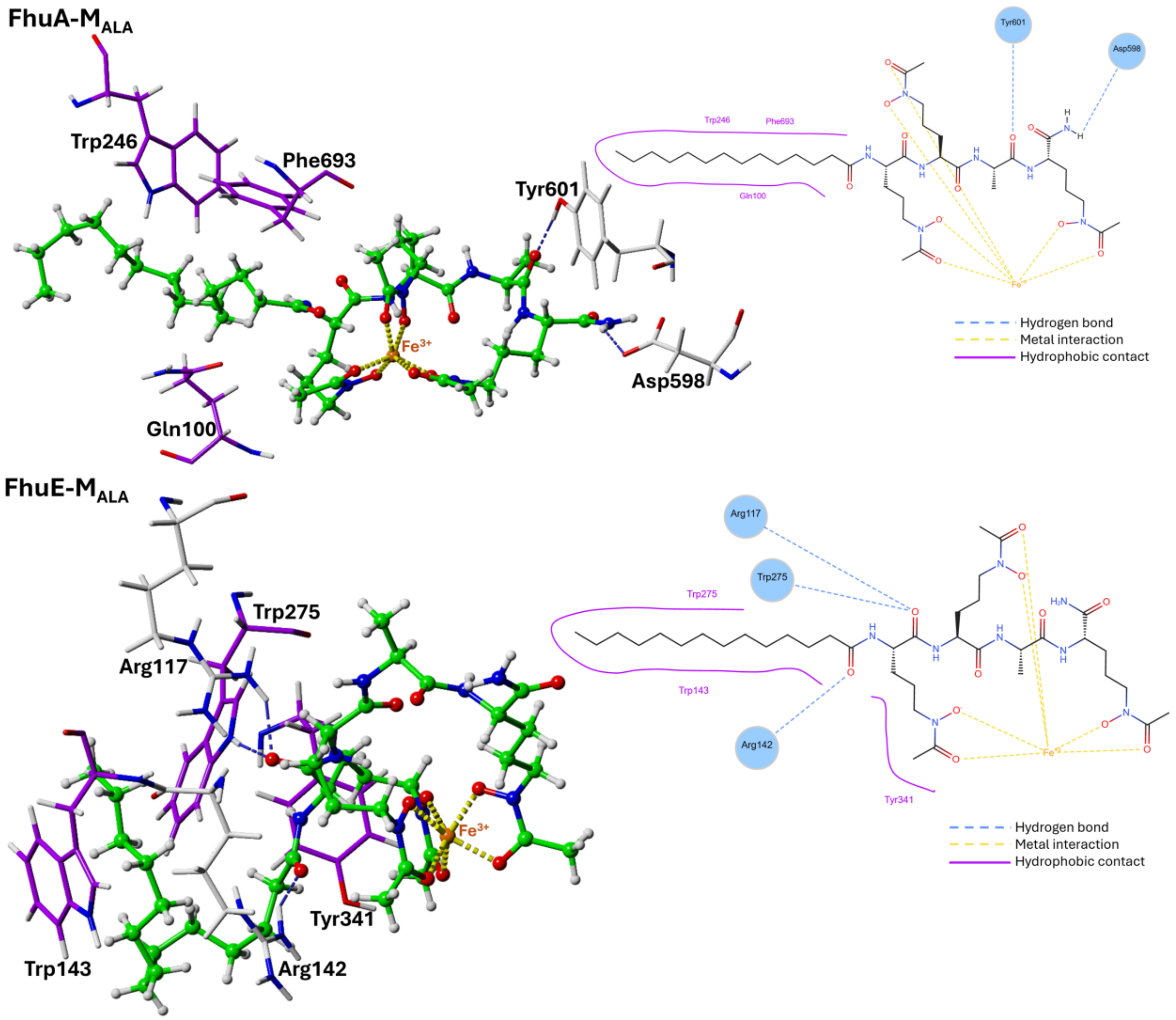


**Supplementary Figure S14.** 3D and 2D depictions of interactions between FhuA (top) or FhuE (bottom) and top-scoring docking poses of $M_{ALA}$. The interactions are based on default definitions used in PoseEdit (Diedrich et al., 2023). Amino acid residues participating in hydrophobic contacts are in purple.

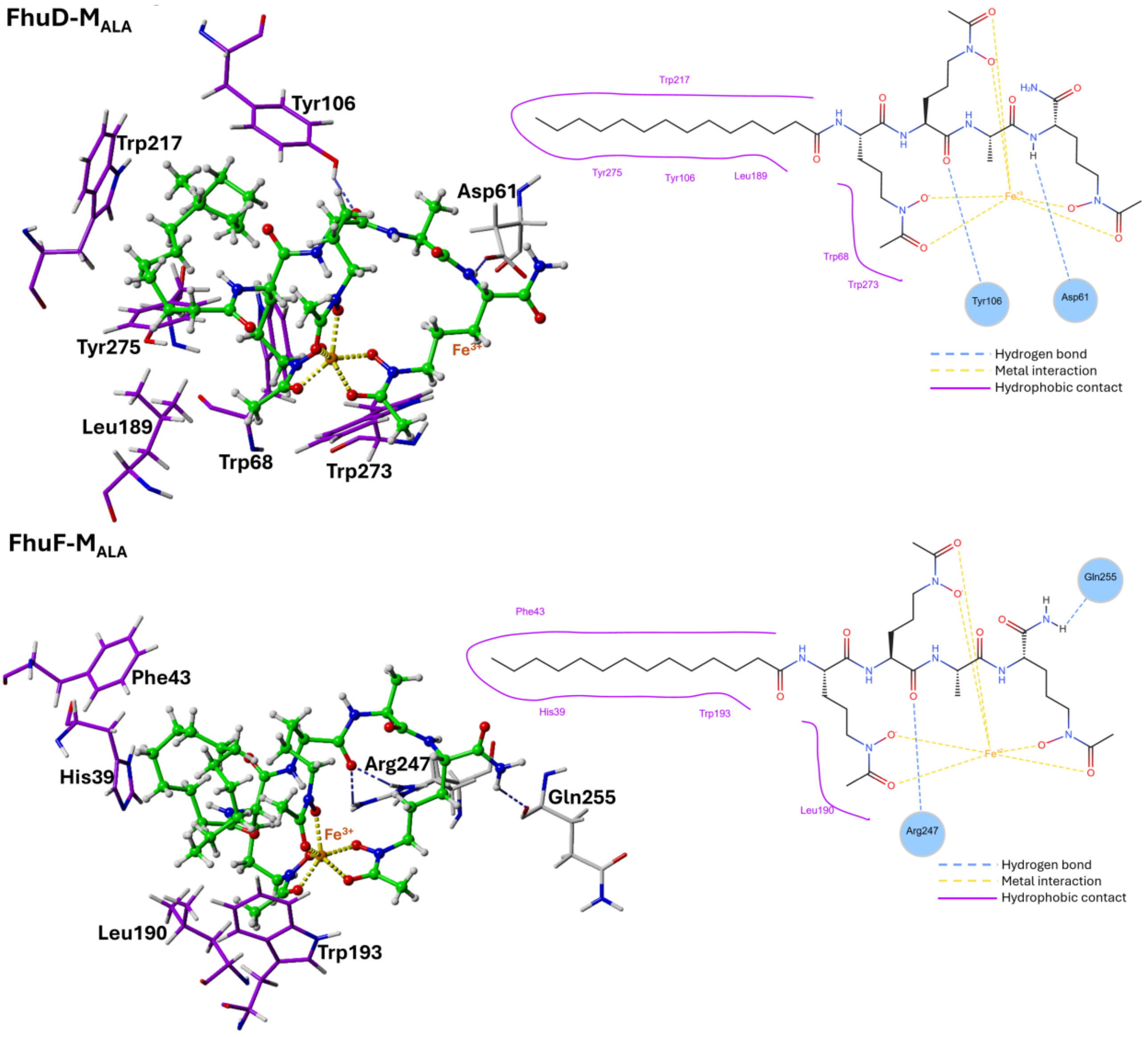


**Supplementary Figure S15.** 3D and 2D depictions of interactions between FhuD (top) or FhuF (bottom) and top-scoring docking poses of M$_{ALA}$. The interactions are based on default definitions used in PoseEdit (Diedrich et al., 2023). Amino acid residues participating in hydrophobic contacts are in purple.

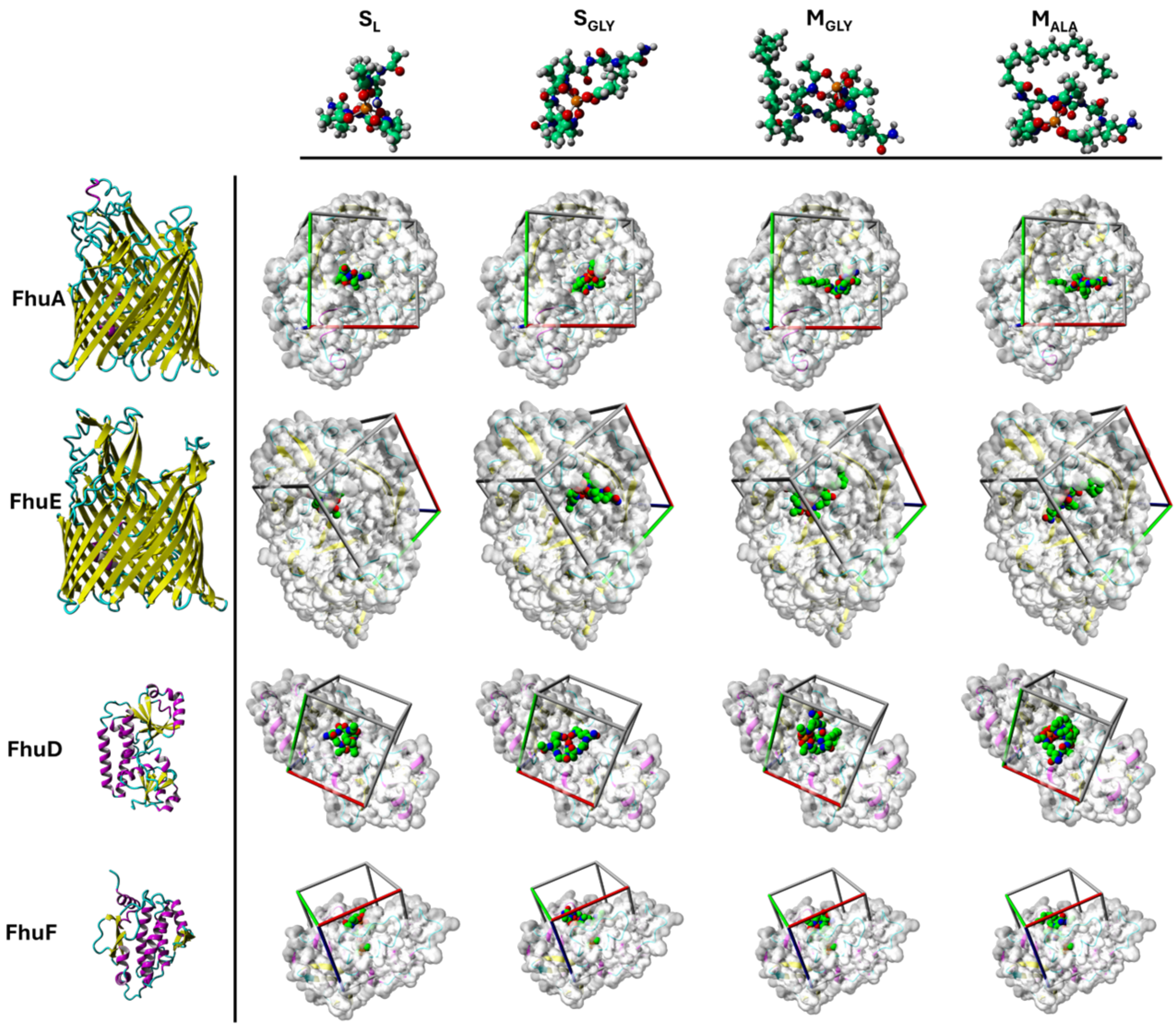


**Supplementary Figure S16.** Overview of docking results. Top-scoring docking poses of synthetic siderophores (columns) docked to target proteins (rows). Hydrogen atoms of the siderophores are omitted from the representations of the protein-ligand complexes for clarity. The cubes depict the boundaries of the docking boxes. A white semi-transparent surface depicts the van der Waals surface of the proteins.

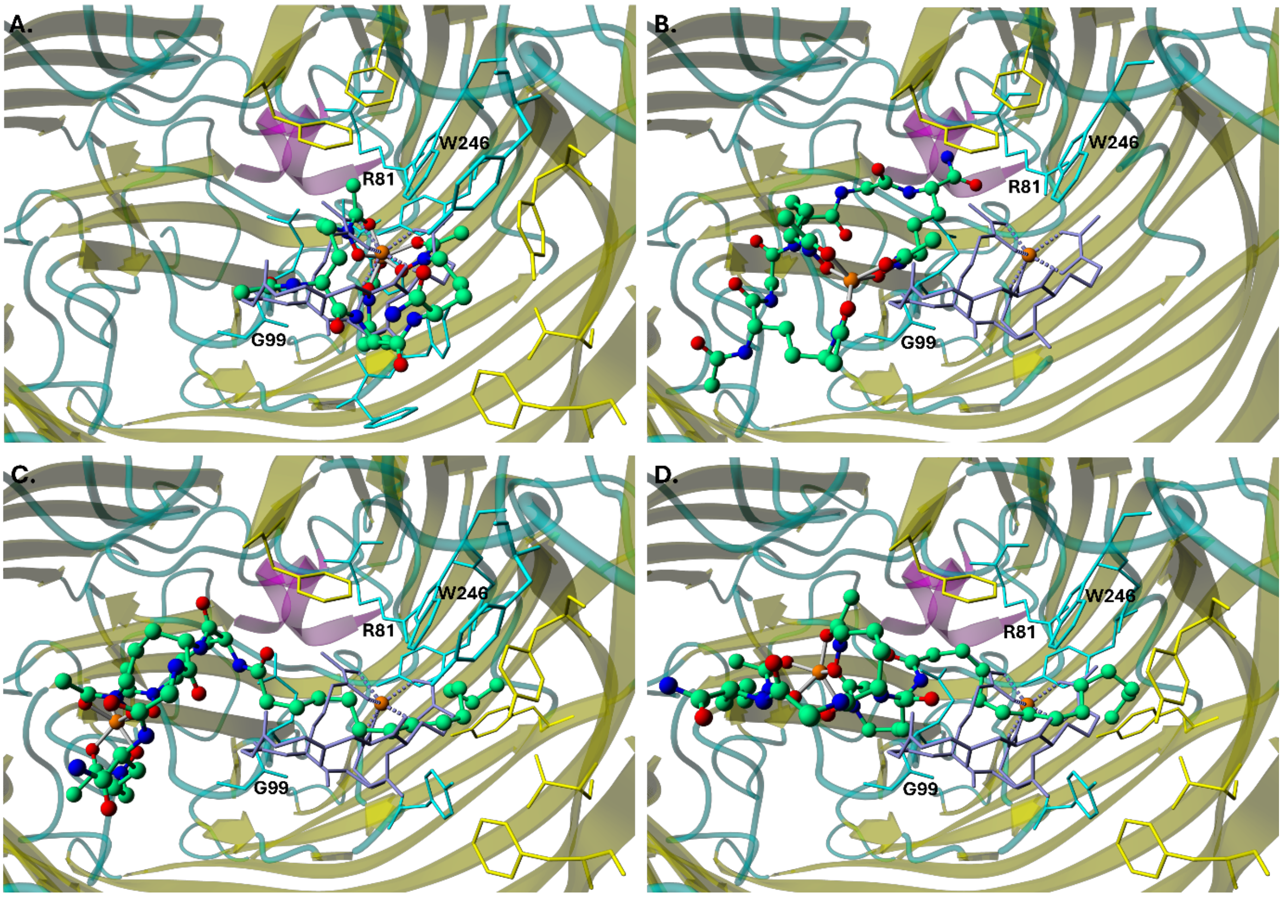


**Supplementary Figure S17.** Top-scoring docking poses of (A) $S_L$, (B) $S_{GLY}$, (C) $M_{GLY}$, and (D) $M_{ALA}$ docked to the FhuA crystal structure (PDB ID: 1by5) in a ball-and-stick representation (colored by atom type), relative to the crystallographic ligand (ferrichrome) in a blue-grey stick representation. Iron(III) ions are depicted as orange spheres. Dative metal coordination bonds are shown as thin solid grey lines for the docked ligands and as thin dotted blue-grey lines for ferrichrome. Amino acid residues within 4 Å of either the ferrichrome or the docked siderophore are shown in stick representation. These residues are colored based on the secondary structure to which they belong. Numbered residues are those involved in key noncovalent interactions with the crystallographic ligand and also interact with the investigated siderophores. Hydrogen atoms are omitted for clarity. The protein is shown as a ribbon model and colored according to its secondary structure based on PDB annotations (helices in magenta, β-sheets in yellow, turns and unstructured fragments in cyan).

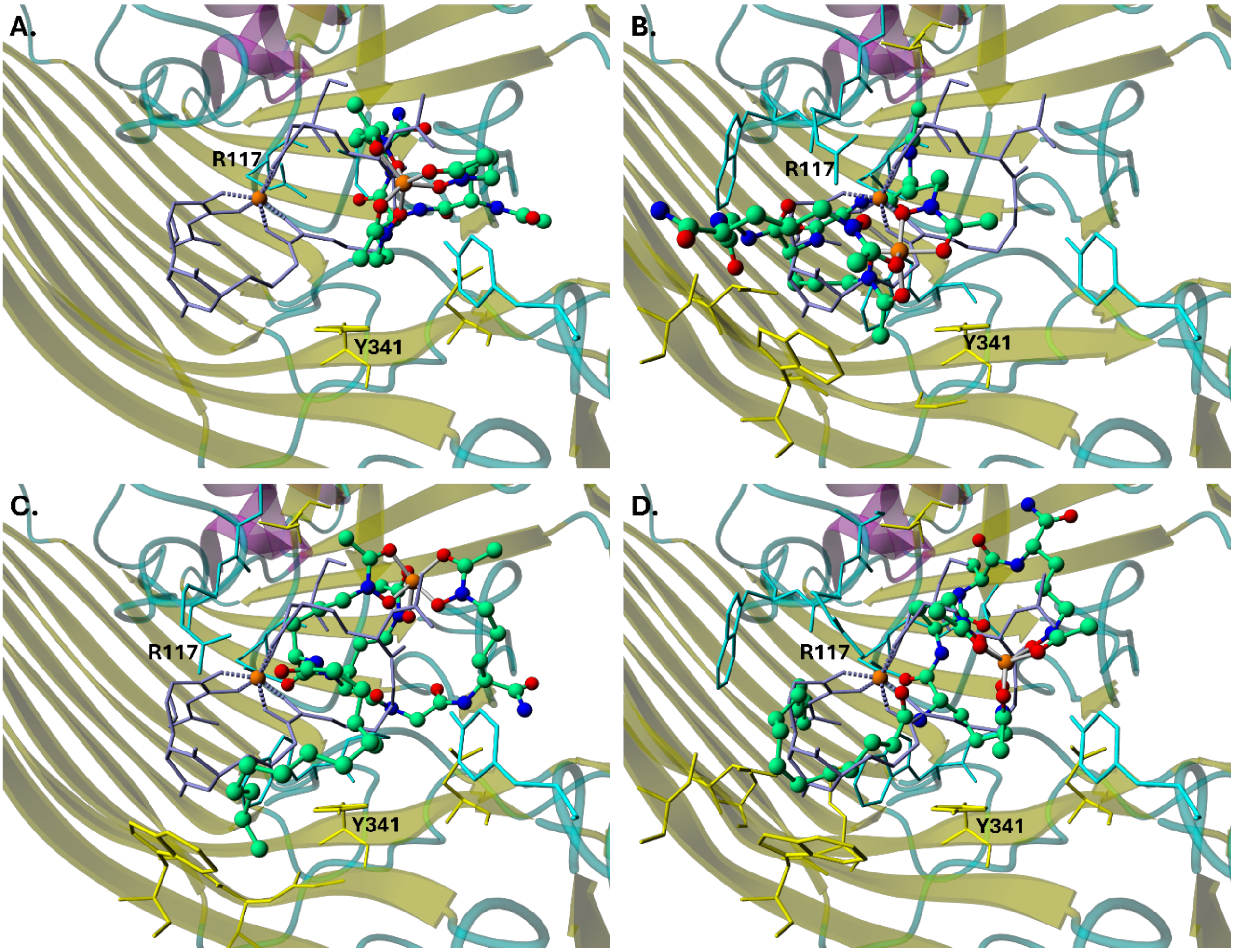


**Supplementary Figure S18.** Top-scoring docking poses of (A) $S_L$, (B) $S_{GLY}$, (C) $M_{GLY}$, and (D) $M_{ALA}$ docked to FhuE crystal structure (PDB ID: 6e4v) in a ball-and-stick representation (colored by atom type), relative to the crystallographic ligand (coprogen) in a blue-grey stick representation. Iron(III) ions are depicted as orange spheres. Dative metal coordination bonds are shown as thin solid grey lines for the docked ligands and as thin dotted blue-grey lines for coprogen. Amino acid residues within 4 Å of either the coprogen or the docked siderophore are shown in stick representation. These residues are colored based on the secondary structure to which they belong. Numbered residues are those involved in key noncovalent interactions with the crystallographic ligand and also interact with the investigated siderophores. Hydrogen atoms are omitted for clarity. The protein is shown as a ribbon model and colored according to its secondary structure based on PDB annotations (helices in magenta, β-sheets in yellow, turns and unstructured fragments in cyan).

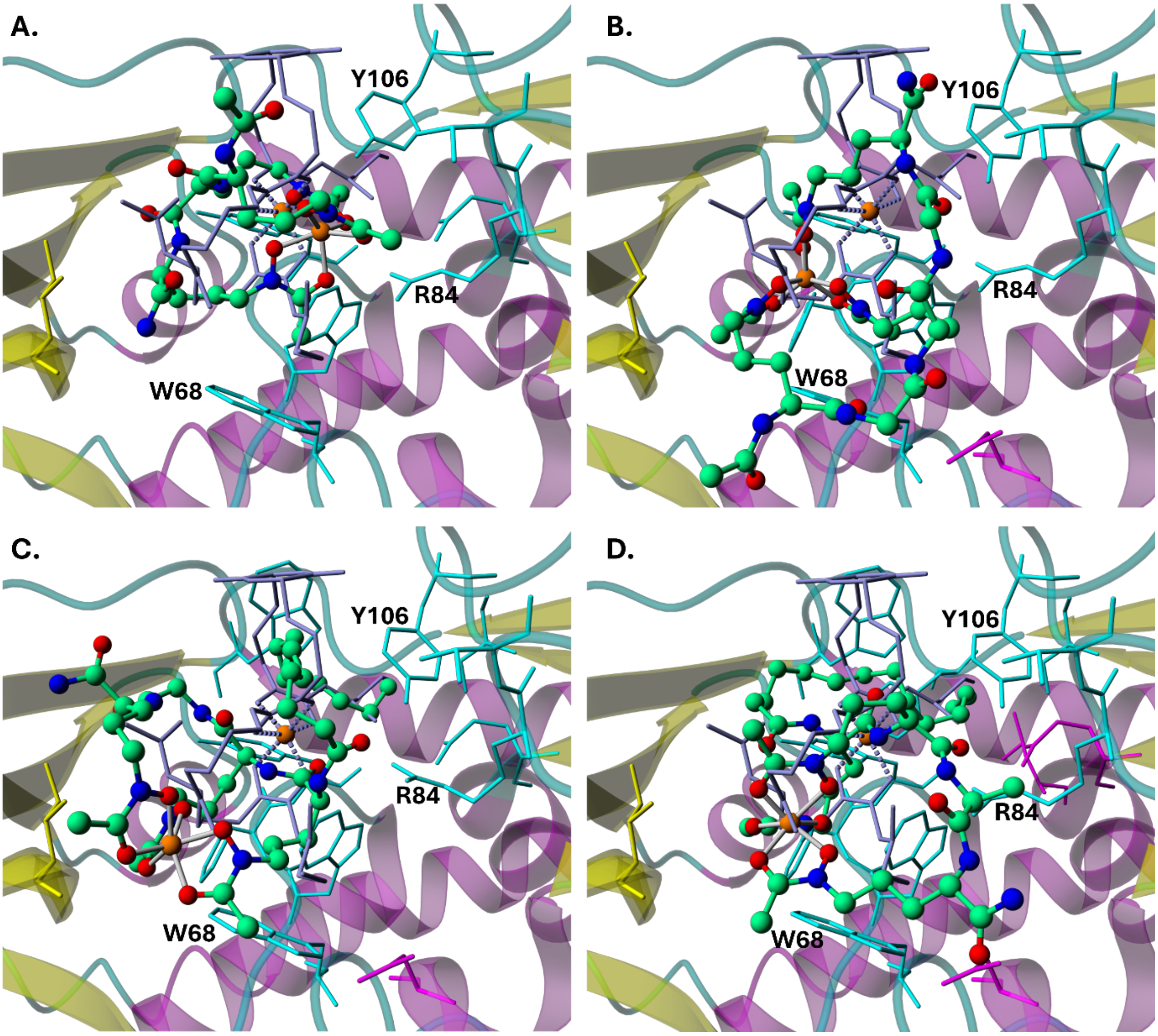


**Supplementary Figure S19.** Top-scoring docking poses of (A) $S_L$, (B) $S_{GLY}$, (C) $M_{GLY}$, and (D) $M_{ALA}$ docked to FhuD crystal structure (PDB ID: 1esz) in a ball-and-stick representation (colored by atom type), relative to the crystallographic ligand (coprogen) in a blue-grey stick representation. Iron(III) ions are depicted as orange spheres. Dative metal coordination bonds are shown as thin solid grey lines for the docked ligands and as thin dotted blue-grey lines for coprogen. Amino acid residues within 4 Å of either the coprogen or the docked siderophore are shown in stick representation. These residues are colored based on the secondary structure to which they belong. Numbered residues are those involved in key noncovalent interactions with the crystallographic ligand and also interact with investigated siderophores. Hydrogen atoms are omitted for clarity. The protein is shown as a ribbon model and colored according to its secondary structure based on PDB annotations (helices in magenta, β-sheets in yellow, turns and unstructured fragments in cyan).

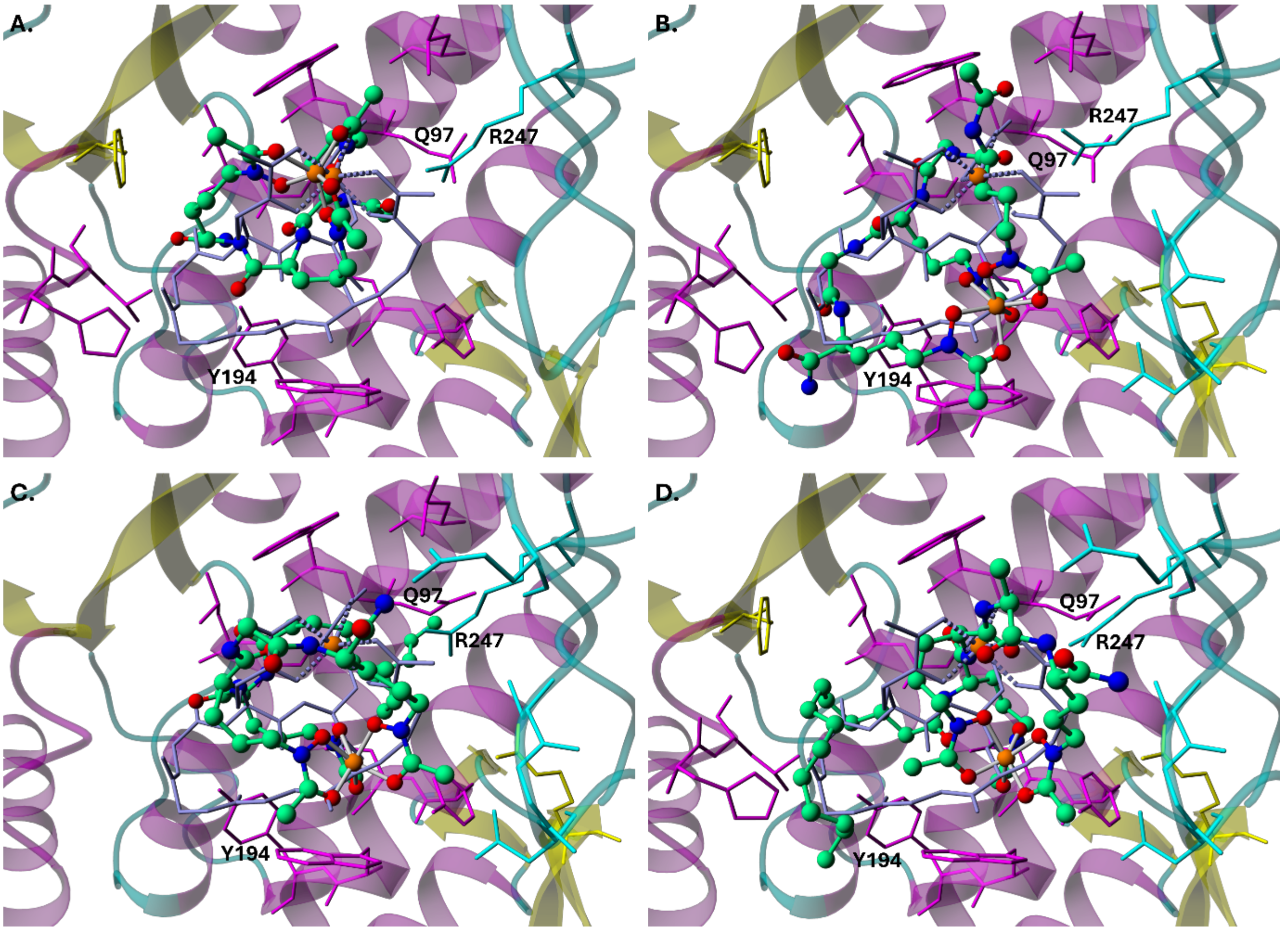


**Supplementary Figure S20.** Top-scoring docking poses of (A) $S_L$, (B) $S_{GLY}$, (C) $M_{GLY}$, and (D) $M_{ALA}$ docked to FhuF crystal structure (PDB ID: 7q5p) in a ball-and-stick representation, relative to physiological ligand obtained in the earlier docking experiment (ferrichrome) in a blue-grey stick representation. Iron(III) ions are depicted as orange spheres. Dative metal coordination bonds are shown as thin solid grey lines for the docked ligands and as thin dotted blue-grey lines for ferrichrome. Amino acid residues within 4 Å of either the ferrichrome or the docked siderophore are shown in stick representation. These residues are colored based on the secondary structure to which they belong. Numbered residues are those involved in key noncovalent interactions with the crystallographic ligand and also interact with investigated siderophores. Hydrogen atoms are omitted for clarity. The protein is shown as ribbon model and colored according to its secondary structure based on PDB annotations (helices in magenta, β-sheets in yellow, turns and unstructured fragments in cyan).

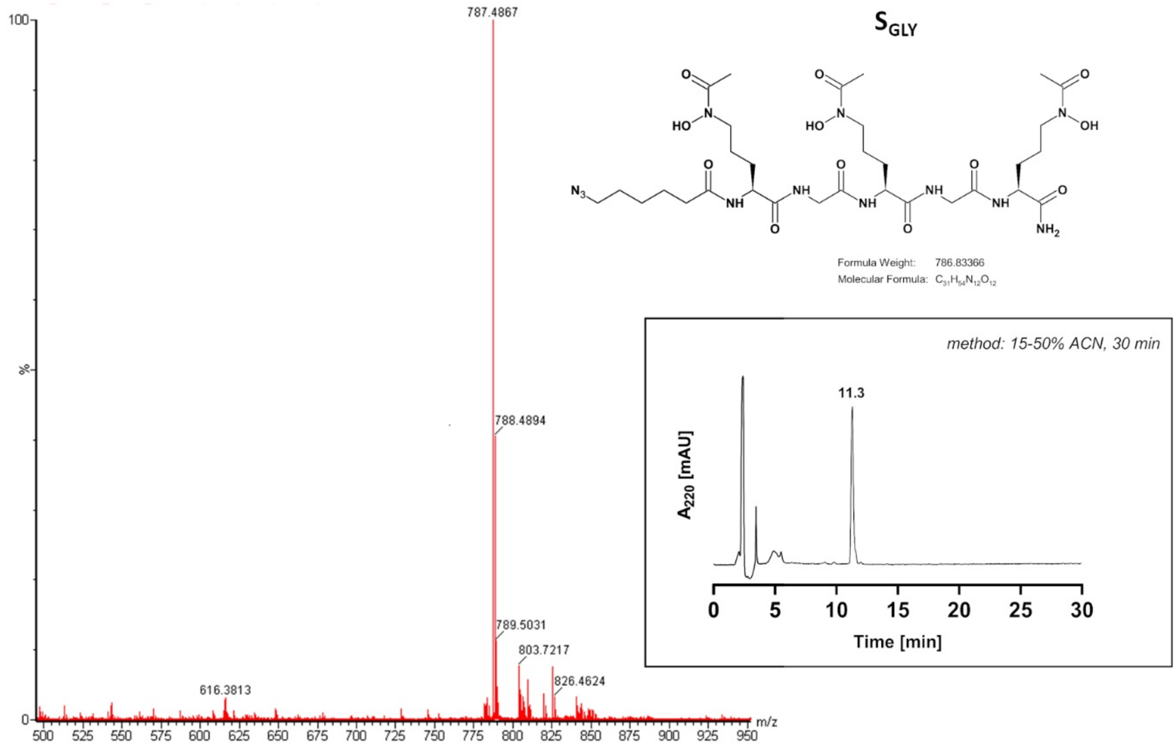


**Supplementary Figure S21.** The MS spectrum and HPLC chromatogram of $S_{GLY}$.

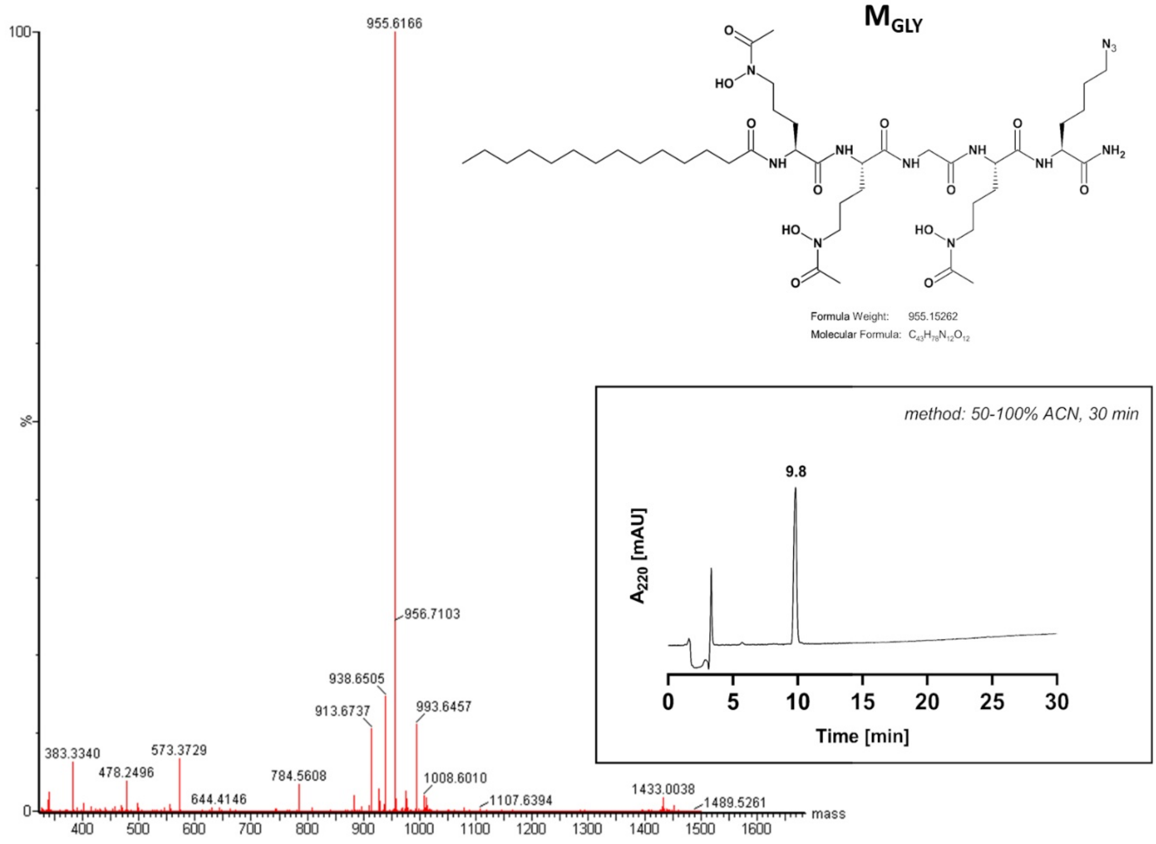


**Supplementary Figure S22.** The MS spectrum and HPLC chromatogram of $M_{GLY}$.

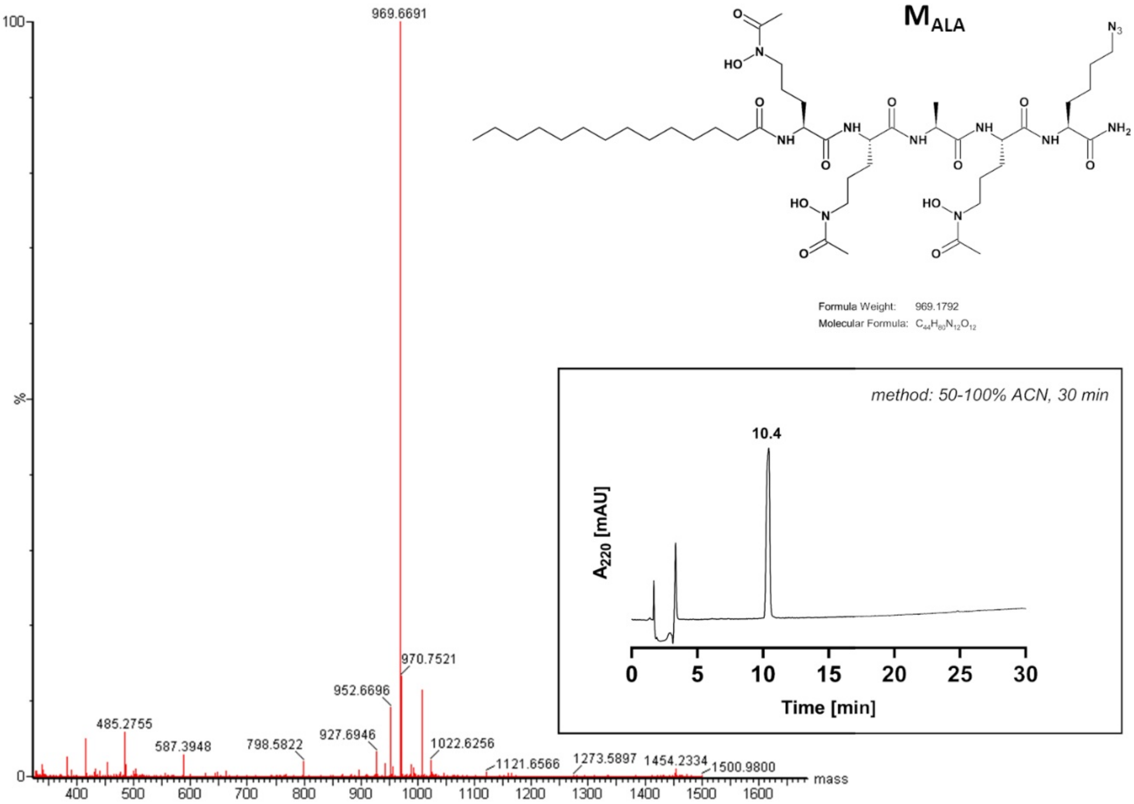


**Supplementary Figure S23.** The MS spectrum and HPLC chromatogram of $M_{ALA}$.

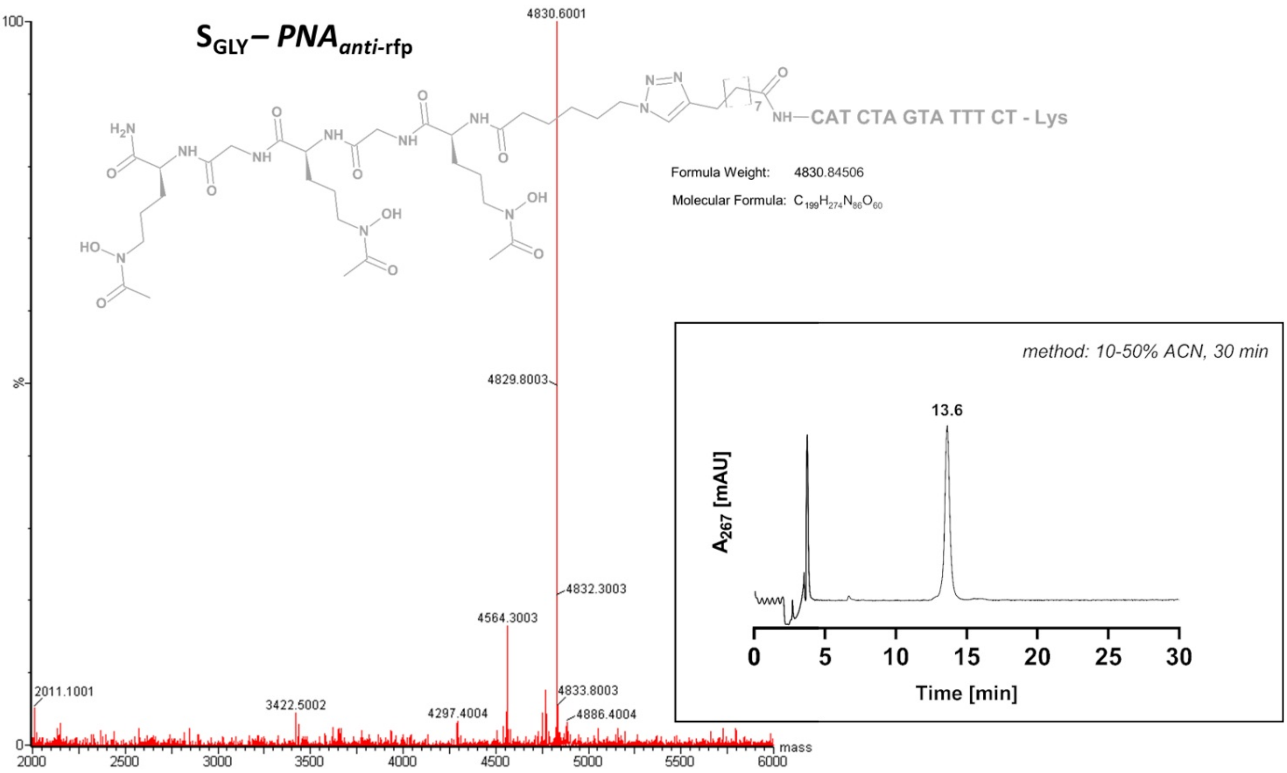


**Supplementary Figure S24.** The MS spectrum and HPLC chromatogram of the $S_{GLY}$ – $PNA_{anti\text{-}rfp}$ conjugate.

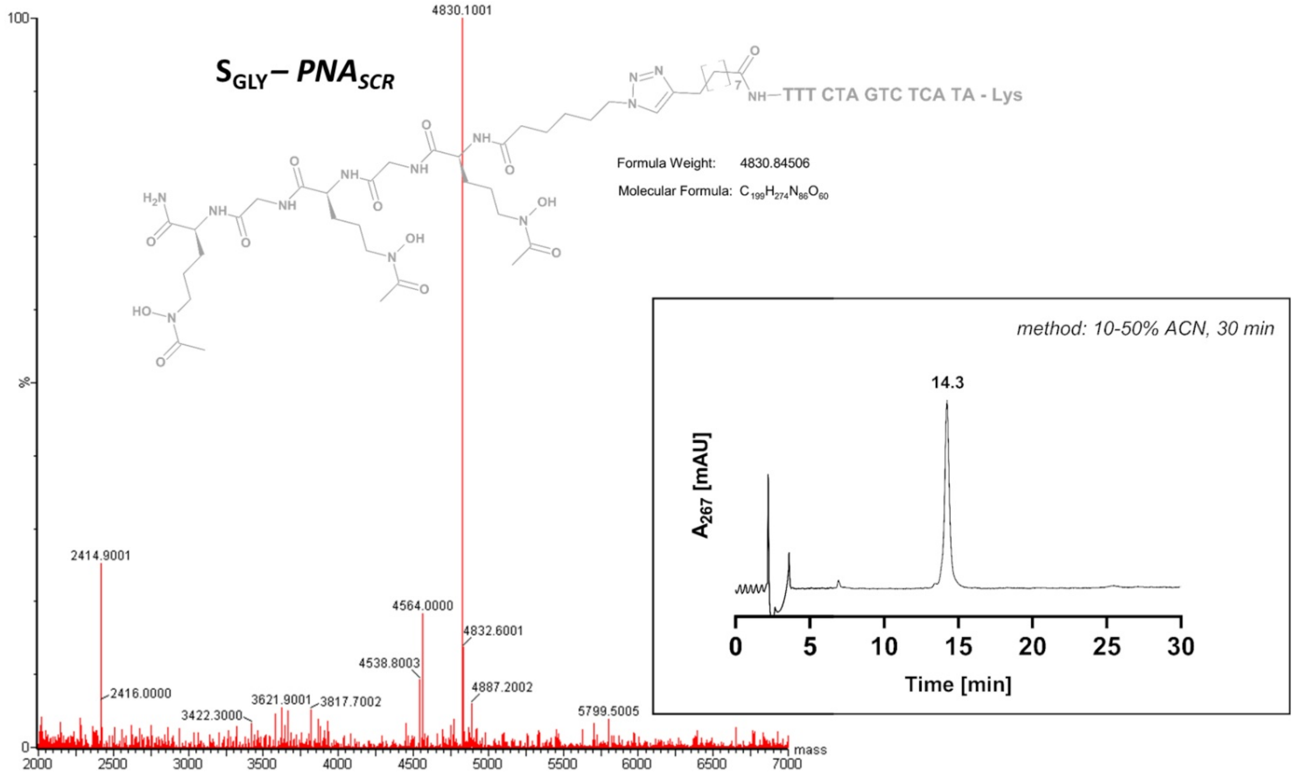


**Supplementary Figure S25.** The MS spectrum and HPLC chromatogram of the $S_{GLY}$ – $PNA_{SCR}$ conjugate.

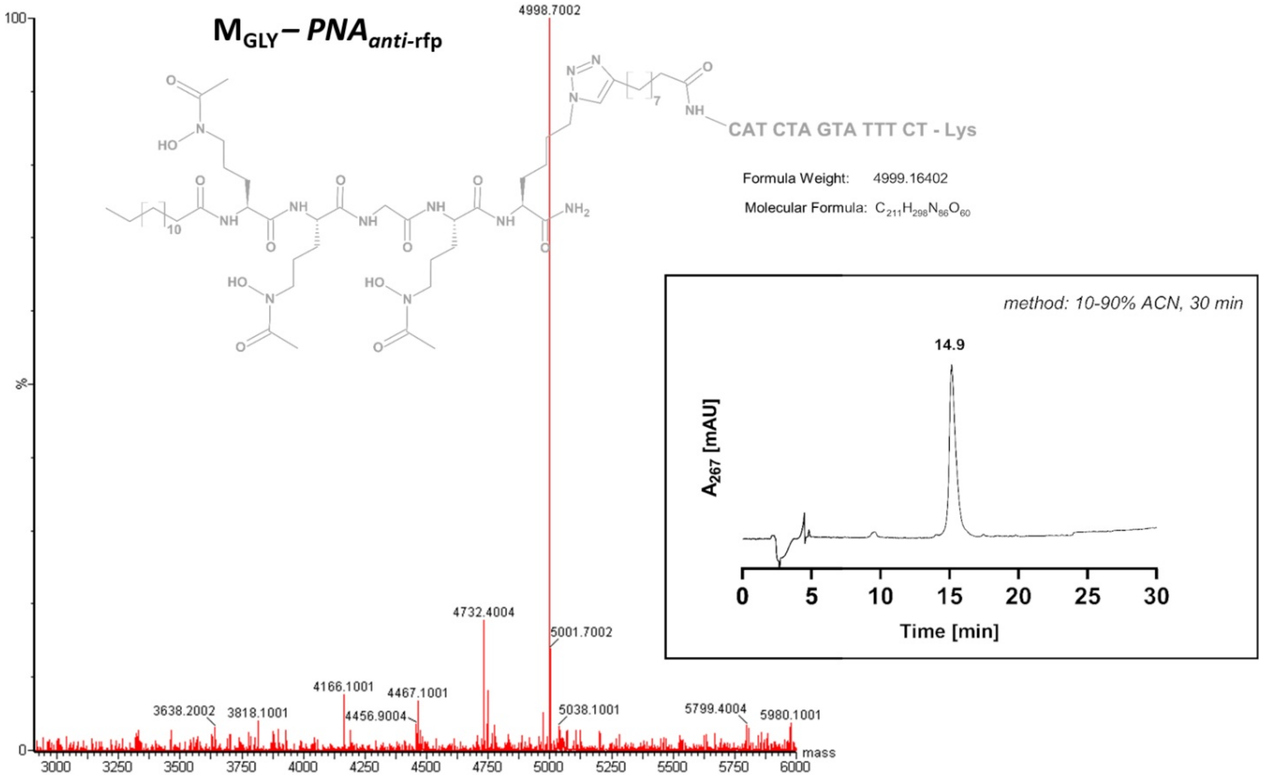


**Supplementary Figure S26.** The MS spectrum and HPLC chromatogram of the $M_{GLY}$ – $PNA_{anti\text{-}rfp}$ conjugate.

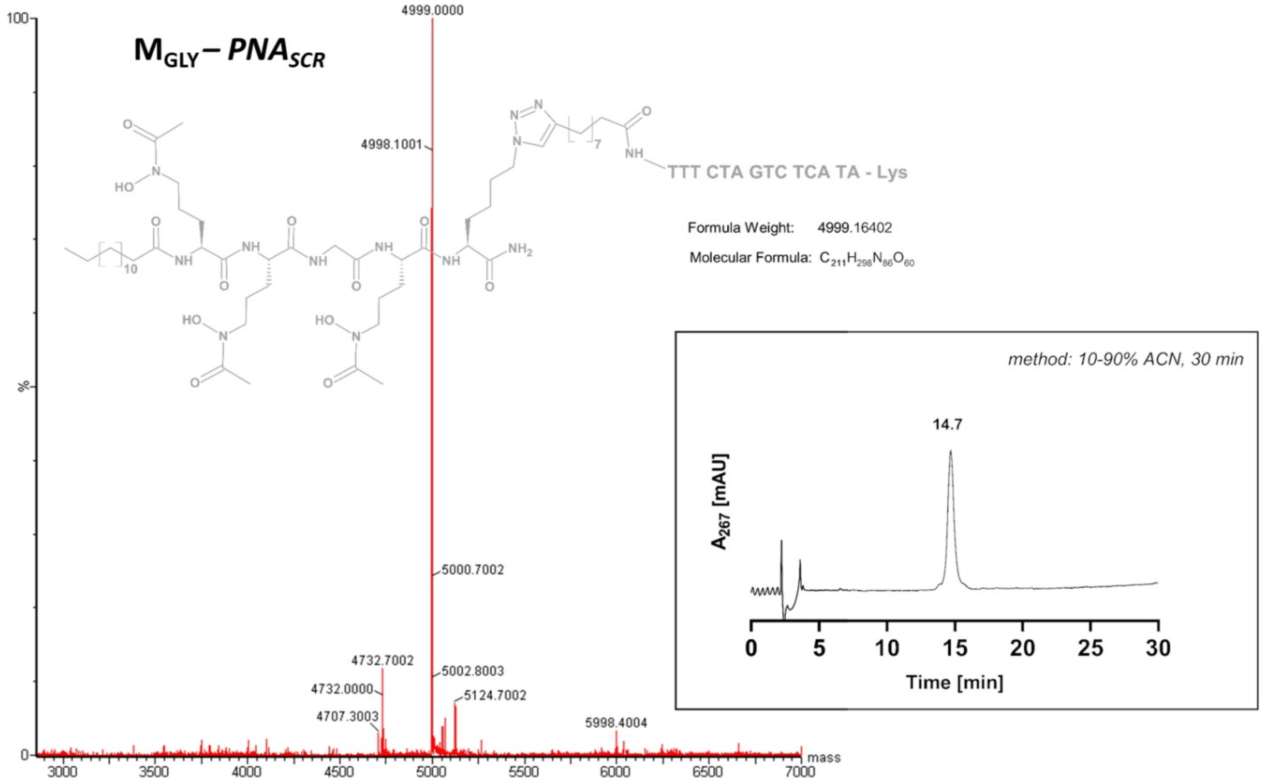


**Supplementary Figure S27.** The MS spectrum and HPLC chromatogram of the $M_{GLY}$ – $PNA_{SCR}$ conjugate.

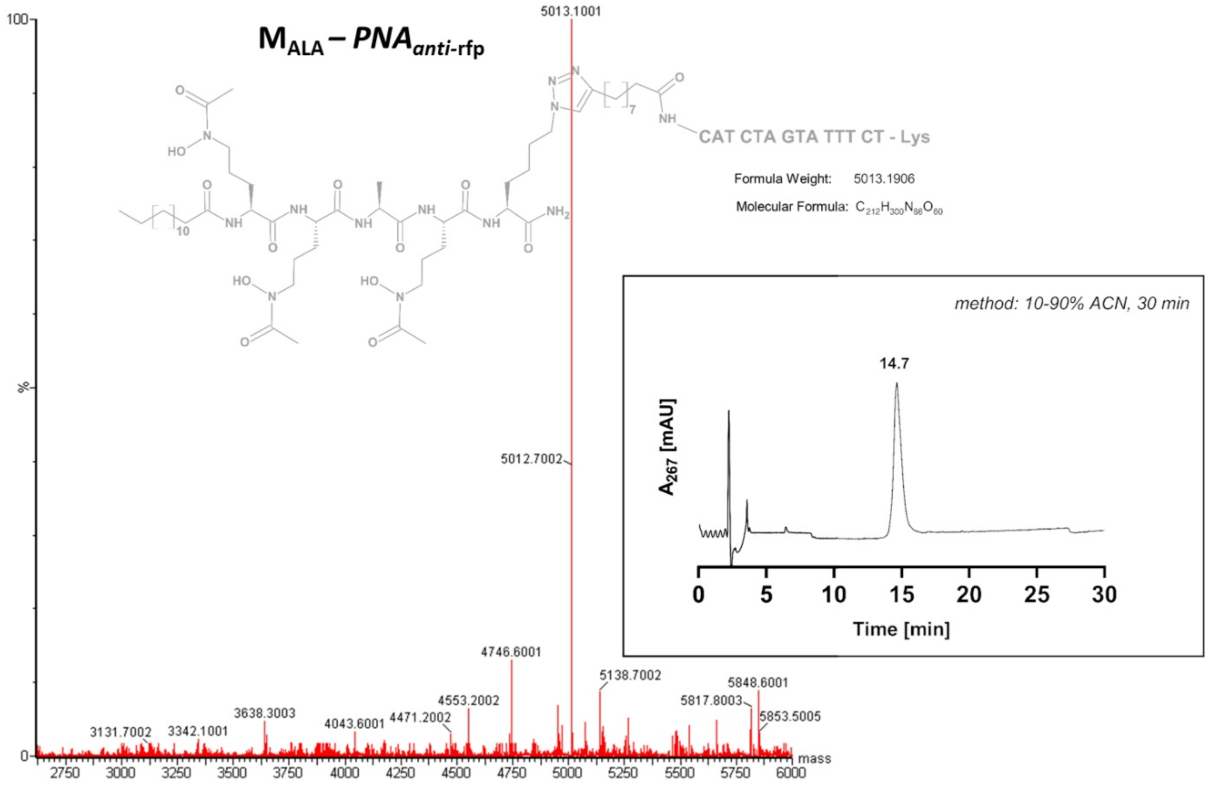


**Supplementary Figure S28.** The MS spectrum and HPLC chromatogram of the $M_{ALA}$ – $PNA_{anti\text{-}rfp}$ conjugate.

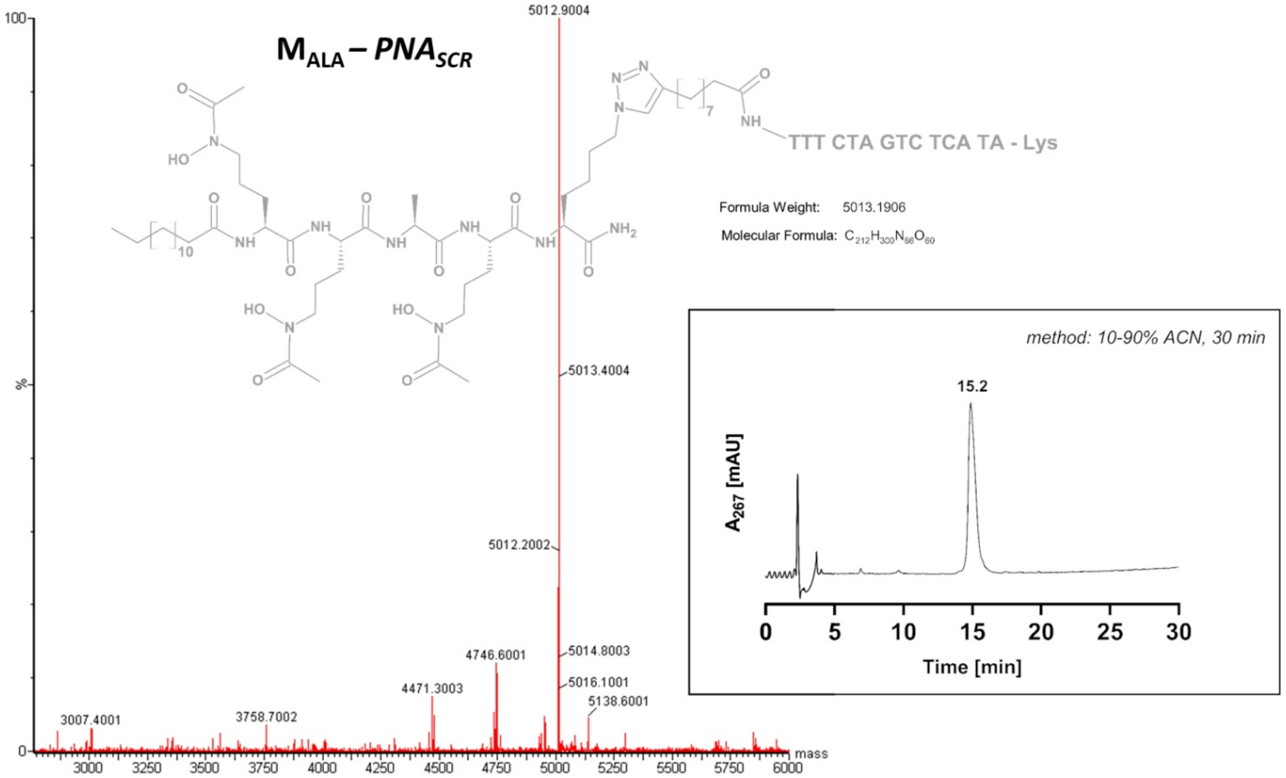


**Supplementary Figure S29.** The MS spectrum and HPLC chromatogram of the $M_{ALA}$ – $PNA_{SCR}$ conjugate.

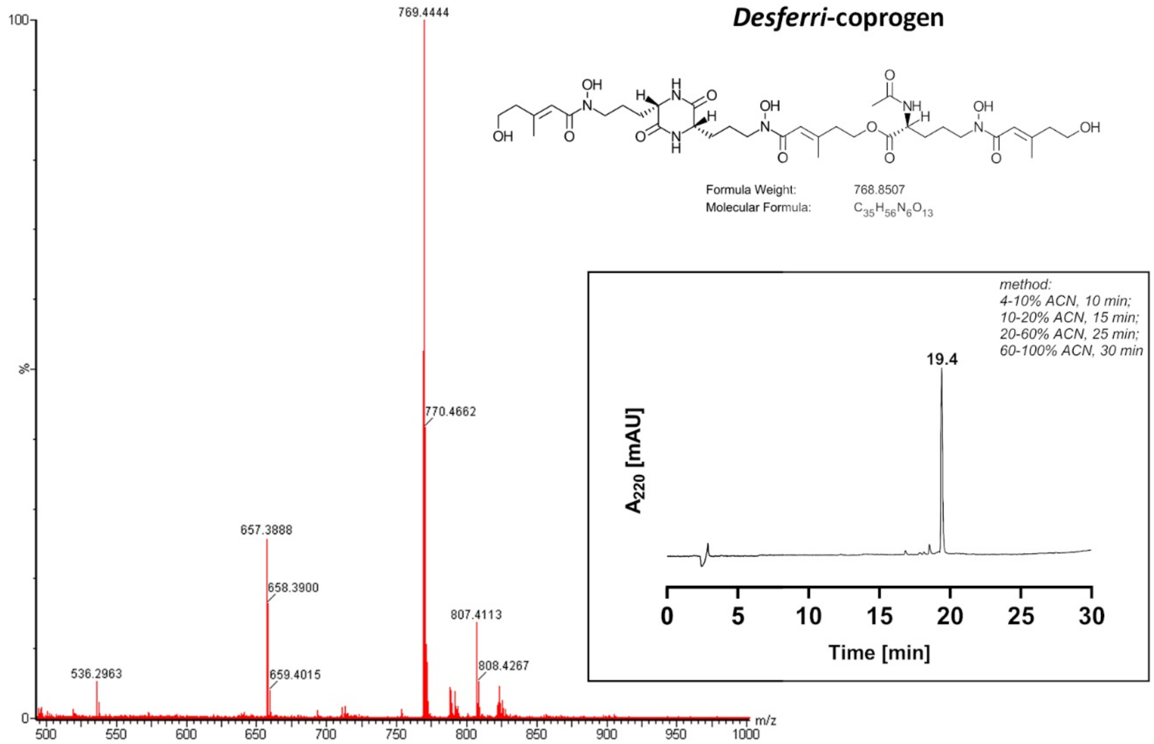


**Supplementary Figure S30.** The MS spectrum and HPLC chromatogram of *desferri*-coprogen.

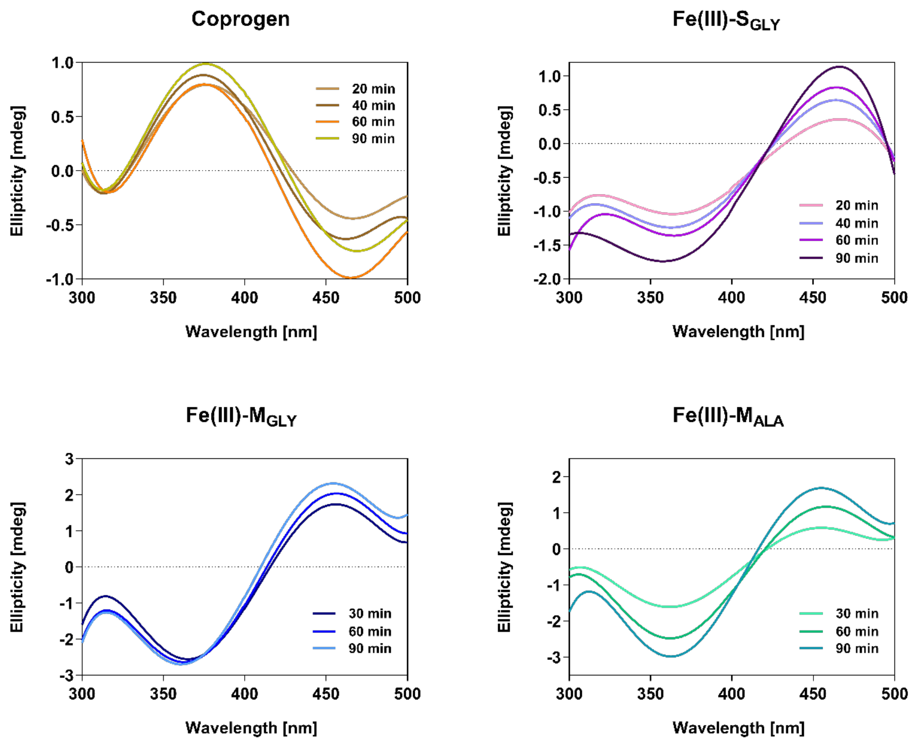


**Supplementary Figure S31.** CD spectra of coprogen and ferric complexes with $S_{GLY}$, $M_{GLY}$, and $M_{ALA}$ recorded after different incubation times at a 1:1 $Fe^{3+}$:siderophore molar ratio. Ferric complexes of coprogen and $S_{GLY}$ were prepared in the phosphate buffer (pH = 7.0). $M_{GLY}$ and $M_{ALA}$ ferric complexes were dissolved in the phosphate buffer/acetonitrile (1:1, v/v) solution.

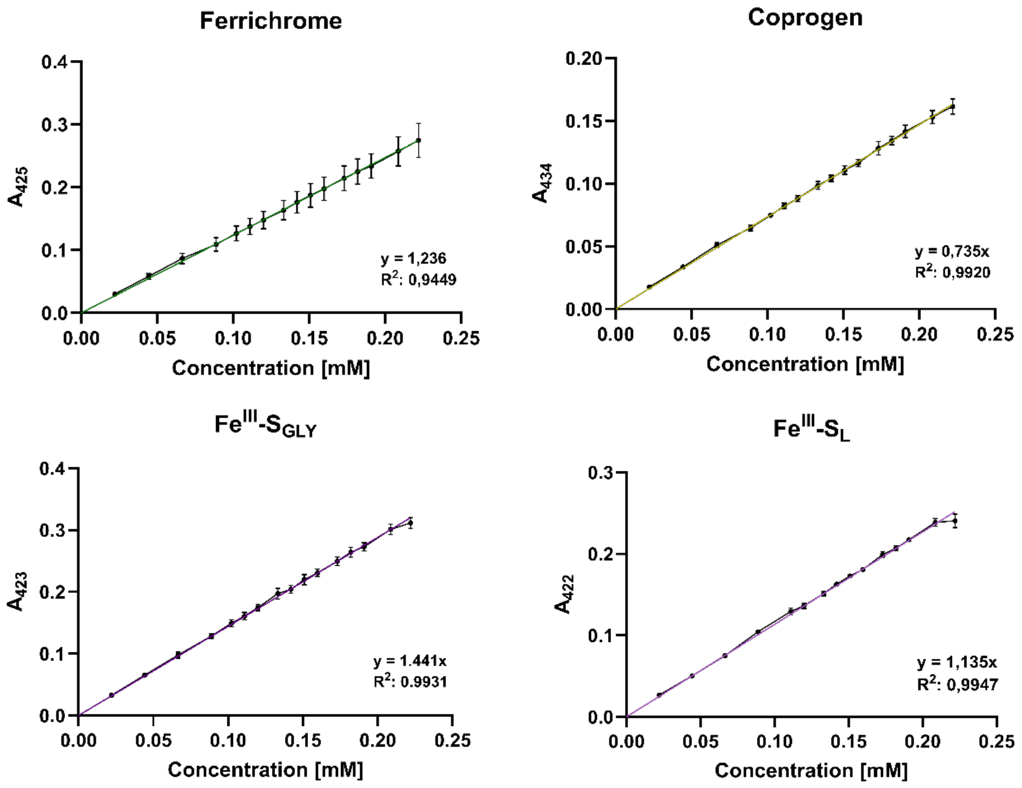


**Supplementary Figure S32.** Calibration curves of ferrichrome, coprogen, and ferric complexes of $S_{GLY}$ and $S_L$ at their respective absorption maximum wavelengths. Solutions were prepared in phosphate buffer (pH = 7.4). Data show averages with SD from three separate experiments (n = 3), each performed in duplicate.

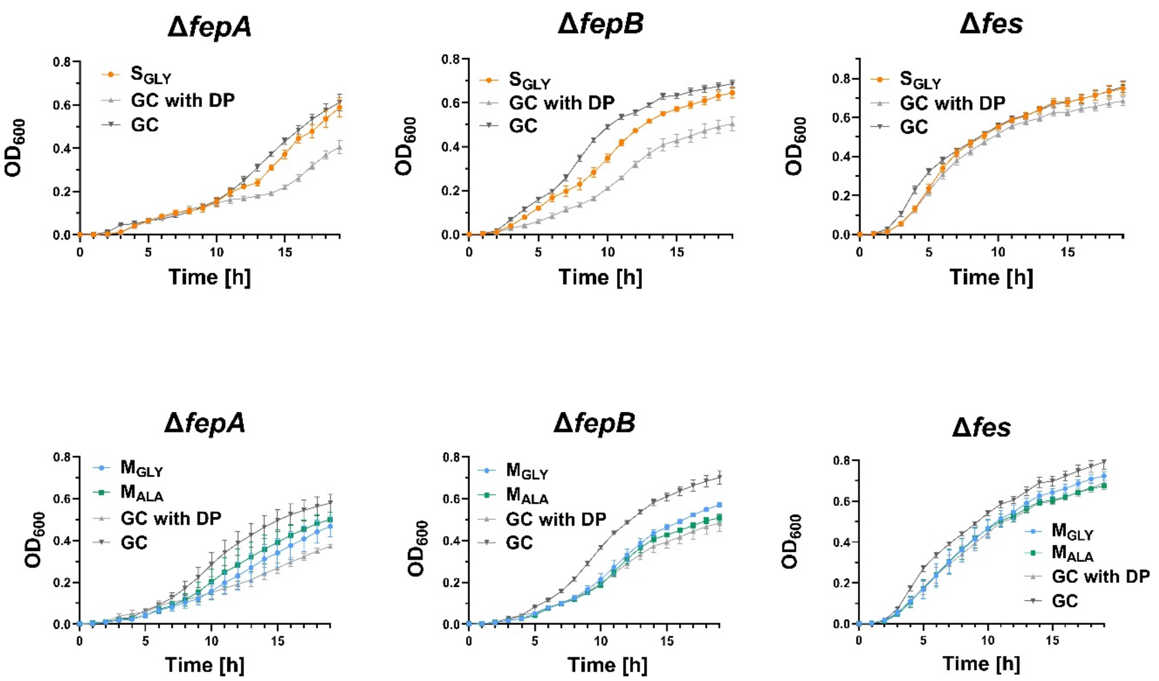


**Supplementary Figure S33.** Growth curves from the growth recovery assay for the Δ*fepA*, Δ*fepB* and Δ*fes* mutant strains cultured under iron-limiting conditions (with DP) without or with $S_{GLY}$, $M_{GLY}$ or $M_{ALA}$ (each at a concentration of 16 µM). $M_{GLY}$ and $M_{ALA}$ were dissolved in DMSO, which was also included and maintained at 3% (v/v) in the corresponding growth control (GC) cultures. The growth was normalized to the $OD_{600}$ measured in the absence of siderophores. Data are shown as mean ± SEM from three independent biological replicates (n = 3), each performed in duplicate.

### Section S1. Steps used during molecular dynamics simulations performed with the LJ 12-6-4 model for iron-siderophore interactions

If mentioned, positional harmonic restraints were applied to non-water heavy atoms, which constitute the solute composed of the siderophore and iron(III) ion.

Energy minimization:

- 3000 steps of steepest descent minimization
- 3000 steps of conjugate gradient minimization with 5 kcal·mol$^{-1}$·Å$^{-2}$ harmonic restraints

Thermalization:

- 1 ns of heating of the system at constant volume from 0.15 K to 310.15 K with 5 kcal·mol$^{-1}$·Å$^{-2}$ restraints. The temperature was increased by 10 K every 15000 simulation steps (corresponding to 30 ps) in 30 stages. Thermalization in final 10 K increase (at 310.15 K) lasted 100 ps.

Equilibration:

- 1 ns of NVT equilibration with 5 kcal·mol$^{-1}$·Å$^{-2}$ restraints
- 1 ns of NPT equilibration with 3 kcal·mol$^{-1}$·Å$^{-2}$ restraints
- 1 ns of NPT equilibration with 1.5 kcal·mol$^{-1}$·Å$^{-2}$ restraints
- 1 ns of NPT equilibration with 1 kcal·mol$^{-1}$·Å$^{-2}$ restraints
- 1 ns of NPT equilibration with 0.5 kcal·mol$^{-1}$·Å$^{-2}$ restraints
- 1 ns of NPT equilibration with 0.1 kcal·mol$^{-1}$·Å$^{-2}$ restraints
- 2.5 ns of NPT equilibration with no restraints

Production runs were performed in the NPT ensemble with no restraints.

**Section S2. Calculations and protocol used for thermodynamic integration (TI)**

The target value of Gibbs free energy Δ*G* for TI was calculated from the formula:

$$\Delta G = -RT \ln K_a$$

where *R* is the gas constant, *T* is the temperature, and $K_a$ is the association constant.

Under the conditions used, Gibbs free energy was approximated by the Helmholtz free energy Δ*F*, calculated using Gaussian quadrature from the following equation:

$$\Delta F = \int_0^1 \left\langle \frac{\partial U(\lambda)}{\partial \lambda} \right\rangle_\lambda d\lambda = \sum_i \left\langle \frac{\partial U(\lambda)}{\partial \lambda} \right\rangle_i w_i$$

where *U*(λ) is the energy of the system dependent on the parameter λ, which describes a degree of alchemical transformation from the unperturbed system to the fully transformed one, $w_i$ is the weight assigned to a specific value of λ. In our system, λ describes changes in the interaction energy occurring while the iron(III) is transformed into a non-interacting pseudo-atom.

The λ values and their corresponding weights $w_i$ are given in Table S3. The twelve λ values (12 windows) were used in one thermodynamic integration calculation. All steps were calculated with pmemd.cuda, with positional harmonic restraints imposed on all non-water heavy atoms with the indicated force constant. In one λ window, the calculations were performed according to the following protocol:

- 6000 steps of steepest descent minimization with 5 kcal·mol$^{-1}$·Å$^{-2}$ restraints
- heating of the system at constant volume from 0.15 K to 293.15 K over 0.5 ns with 5 kcal·mol$^{-1}$·Å$^{-2}$ restraints
- 1 ns of NPT equilibration with 2 kcal·mol$^{-1}$·Å$^{-2}$ restraints
- 2 ns of NPT equilibration with no restraints
- 5 ns of production in the NPT ensemble with no restraints.

An integration time step of 2 fs was used with SHAKE (Miyamoto & Kollman, 1992; Ryckaert et al., 1977) for bonds involving hydrogen atoms. For long-range electrostatic interactions, Particle Mesh Ewald (PME) summation (Cheatham et al., 1995; Darden et al., 1993; Petersen, 1995) was used. For non-bonded interactions, a cutoff of 9 Å was applied. A Langevin thermostat was used for temperature control, with a collision frequency of 1 ps$^{-1}$. In the steps with pressure control, a Monte Carlo barostat, set at 1 atm, was used. Smoothstep functions (Lee et al., 2020) with default settings were used to parametrize alchemical transformations to achieve smoother changes of the λ coupling parameter.

**Supplementary Table S3.** The λ values and their corresponding weights $w_i$ used in Gaussian quadrature for the 12 windows of the thermodynamic integration protocol.

| λ | | $w_i$ |
|---|---|---|
| 0.00922 | 0.99078 | 0.02359 |
| 0.04794 | 0.95206 | 0.05347 |
| 0.11505 | 0.88495 | 0.08004 |
| 0.20634 | 0.79366 | 0.10158 |
| 0.31608 | 0.68392 | 0.11675 |
| 0.43738 | 0.56262 | 0.12457 |

**Supplementary Table S4.** Chosen set of $C_{ij}$ parameters that reproduces the calculated free energy from the experimental association constant and the binding geometry. Atom names are consistent with those in Supplementary Table S2. * Default values calculated from polarizabilities contained in the lj_1264_pol.dat AMBER file.

| Atom | $C_{ij}$ value [$Å^4$ × kcal/mol] |
|---|---|
| OZ | 174.167590* |
| NE | 333.642659* |
| OH | 330 |
| OW (oxygen atoms of water molecules) | 442* |

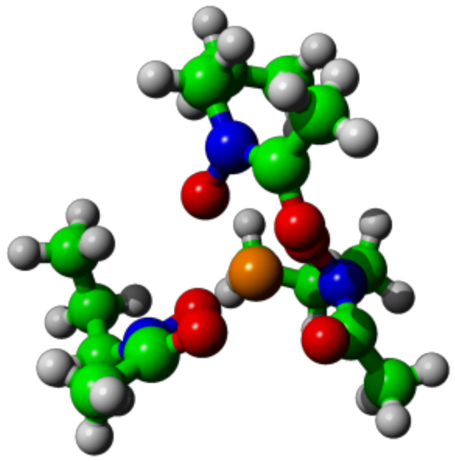

**Supplementary Figure S34.** The input structure for the MCPB.py force constant calculations. The system consisted of AHO sidechains and an iron(III) ion derived from the ferrichrome crystal structure (CCDC ID: 1154895).

### Section S3. Modifications in the force field

The force field modification file was obtained using the MCPB.py program. Parameters for bonds, angles, and dihedrals that do not involve the M1 iron are the same as in ff19SB supplemented by GAFF2. M1 uses nonbonded parameters from the IOD set for iron(III). M and Y were used as atom names to follow the default naming convention given in MCPB.py for the electron acceptor and donor, respectively.

```
REMARK GOES HERE, THIS FILE IS GENERATED BY MCPB.PY
MASS
M1 55.85                           Fe ion
Y1 16.000         0.434
Y2 16.000         0.434
Y1 16.000         0.434
Y2 16.000         0.434
Y1 16.000         0.434
Y2 16.000         0.434

BOND
M1-Y1   82.6      1.9690
M1-Y2   46.1      2.0845
C -Y2  570.00   1.229
N -Y1  380.90   1.218

ANGL
M1-Y1-N     77.31       113.02
M1-Y2-C     84.98       112.02
Y1-M1-Y1    46.81       93.43
Y2-M1-Y1    55.88       114.20
Y2-M1-Y2    76.78       90.47
C -N -Y1  103.590       116.410
CT-C -Y2   80.000       120.400
CT-N -Y1   98.004       121.730
N -C -Y2   80.000       122.900

DIHE
C -Y2-M1-Y1 3     0.00        0.00   3.0
C -Y2-M1-Y2 3     0.00        0.00   3.0
CT-C -N -Y1 4     10.0        180.0   2.0
CT-CT-N -Y1 6     0.0         0.0   2.0
CT-N -C -Y2 4     10.0        180.0   2.0
HC-CT-C -Y2 1     0.8         0.0  -1.0
HC-CT-C -Y2 1     0.0         0.0  -2.0
HC-CT-C -Y2 1     0.08        180.0   3.0
M1-Y1-N -C  3     0.00        0.00   3.0
M1-Y1-N -CT 3     0.00        0.00   3.0
M1-Y2-C -CT 3     0.00        0.00   3.0
M1-Y2-C -N  3     0.00        0.00   3.0
N -Y1-M1-Y1 3     0.00        0.00   3.0
N -Y1-M1-Y2 3     0.00        0.00   3.0
Y1-M1-Y1-N  3     0.00        0.00   3.0
Y1-M1-Y2-C  3     0.00        0.00   3.0
Y1-N -CT-H1 6     0.0         0.0   2.0
Y2-C -N -Y1 4     10.0        180.0   2.0
Y2-M1-Y1-N  3     0.00        0.00   3.0
Y2-M1-Y2-C  3     0.00        0.00   3.0

IMPR
X -X -C -Y2       10.5        180.        2.
C -CT-N -Y1       1.1         180.0       2.0
CT-N -C -Y2       10.5        180.0       2.0

NONB
  M1        1.3860  0.0135709700
  Y1        1.6612  0.2100
  Y2        1.6612  0.2100
  Y1        1.6612  0.2100
  Y2        1.6612  0.2100
  Y1        1.6612  0.2100
  Y2        1.6612  0.2100
```

**Supplementary Table S5.** Names, atom types, and partial charges of atoms in the myristic acid residue.

| Atom name | Atom type | Charge [e] | Structure |
|---|---|---|---|
| C12 | cD | -0.166449 | |
| H2S, H2R | hL | 0.053162 | |
| C13 | cD | -0.002433 | |
| H3S, H3R | hL | 0.018641 | |
| C14 | cD | -0.027370 | |
| H4S, H4R | hL | 0.020121 | |
| C15 | cD | -0.016726 | |
| H5R, H5S | hL | 0.010455 | |
| C16 | cD | -0.028181 | |
| H6R, H6S | hL | 0.011342 | |
| C17 | cD | -0.004460 | |
| H7R, H7S | hL | 0.006707 | |
| C18 | cD | -0.026863 | |
| H8S, H8R | hL | 0.011047 | |
| C19 | cD | -0.014901 | |
| H9R, H9S | hL | 0.010060 | |
| C110 | cD | -0.021288 | |
| H10S, H10R | hL | 0.009863 | |
| C111 | cD | -0.032236 | |
| H11R, H11S | hL | 0.012526 | |
| C112 | cD | -0.028485 | |
| H12S, H12R | hL | 0.016964 | |
| C113 | cD | 0.017260 | |
| H13R, H13S | hL | 0.007397 | |
| C114 | cD | -0.113230 | |
| H14R, H14S, H14T | hL | 0.025151 | |
| C | C | 0.589019 | |
| O | O | -0.575679 | |

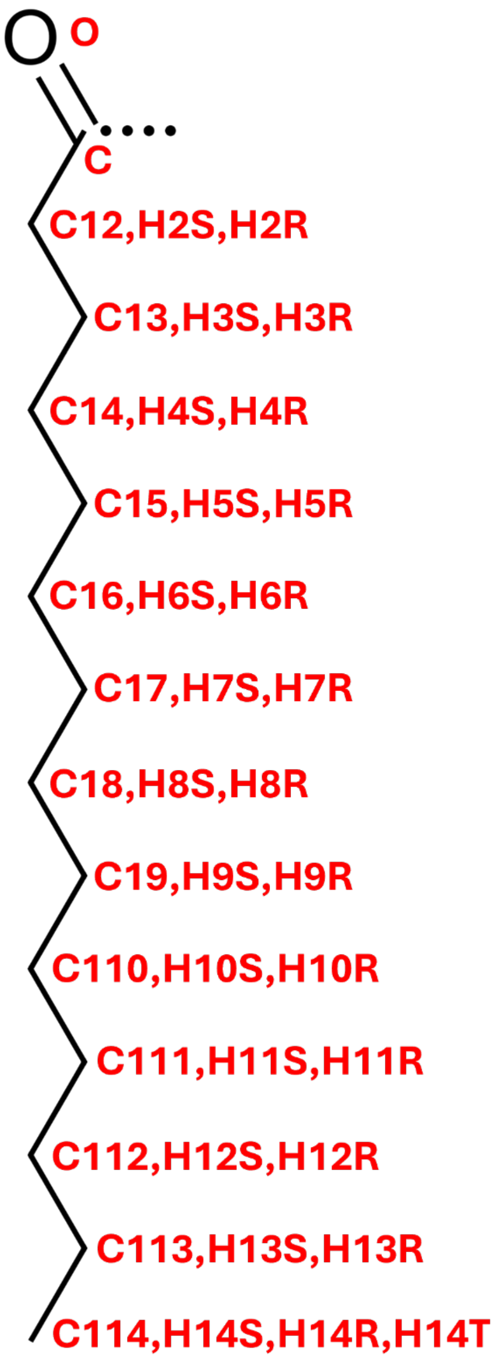

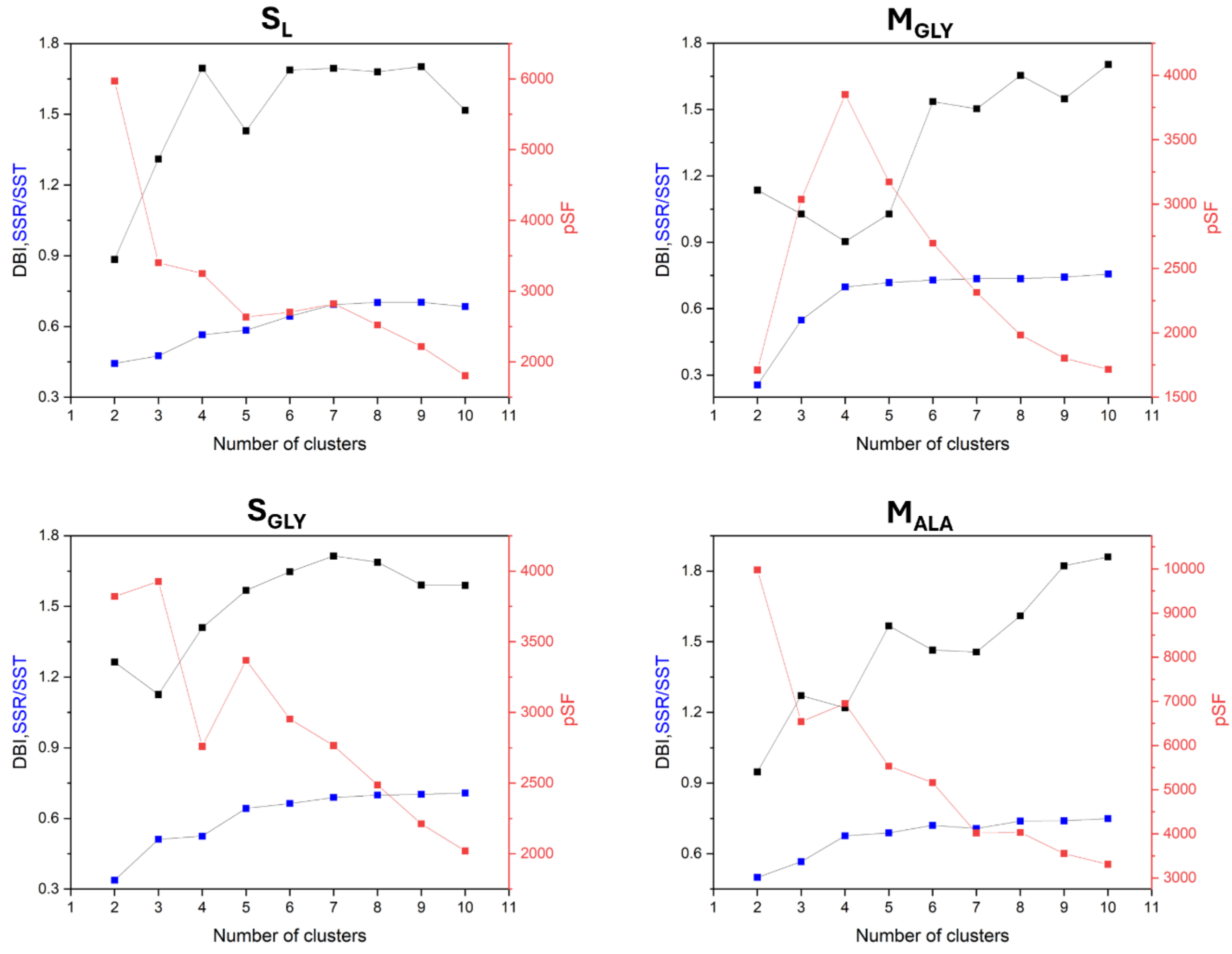


**Supplementary Figure S35.** Clustering metrics for the simulations of siderophores using the nonbonded LJ 12-6-4 model. Abbreviations: Davies-Bouldin index (DBI), pseudo-F statistic (pSF), sum of squares regression (SSR), and total sum of squares (SST). Based on these metrics, the number of clusters was selected as two for $S_L$ and $M_{ALA}$, three for $S_{GLY}$, and four for $M_{GLY}$.

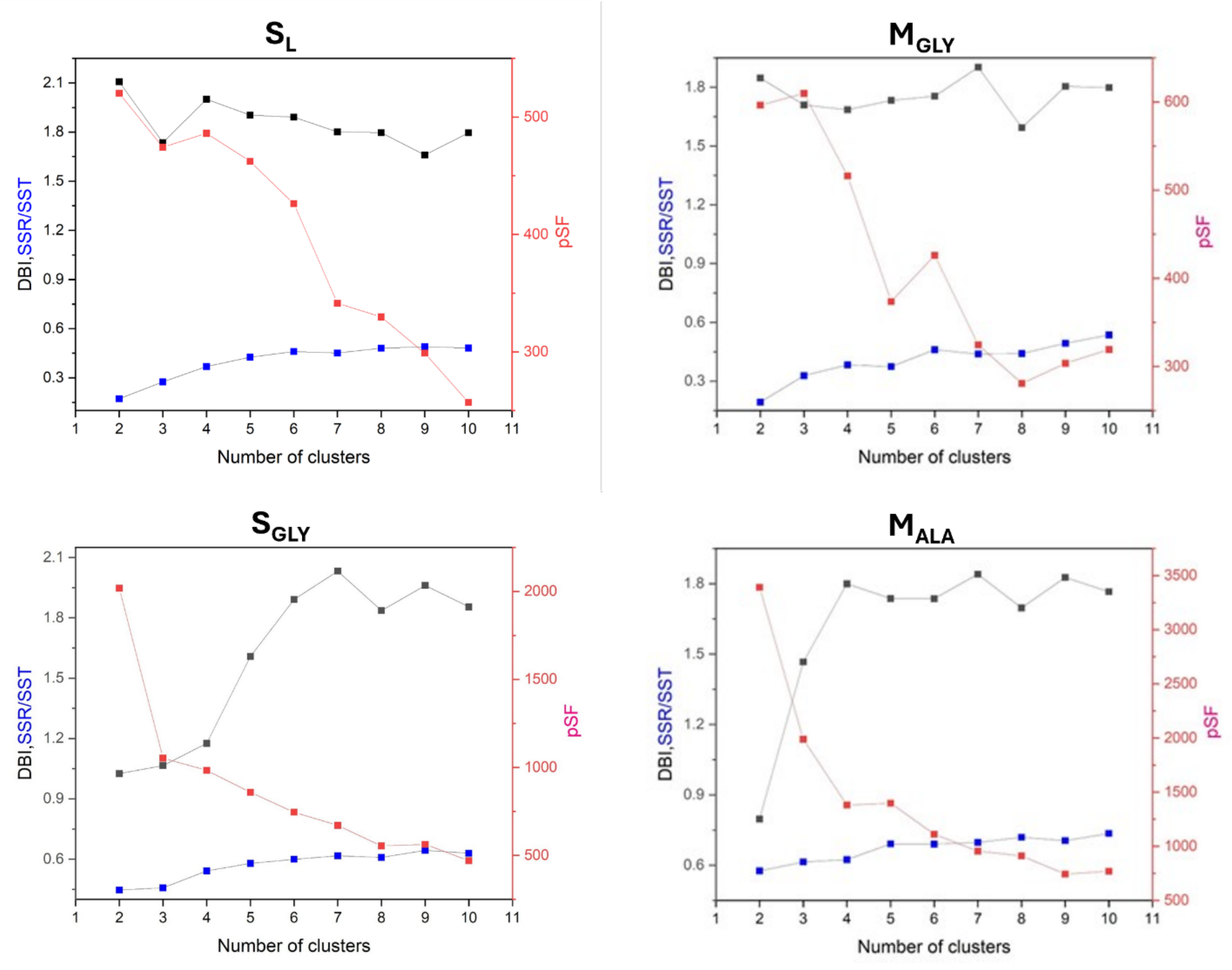


**Supplementary Figure S36.** Clustering metrics for the simulations of siderophores using the covalent bonded model for iron-siderophore interactions. Abbreviations: Davies-Bouldin index (DBI), pseudo-F statistic (pSF), sum of squares regression (SSR), and total sum of squares (SST). Based on these metrics, the number of clusters was selected as two for $S_{GLY}$ and $M_{ALA}$, and three for $S_{L}$ and $M_{GLY}$.
